# Breakthrough in Dicot Prime Editing: Enabling Heritable Desired Edits in Tomato and *Arabidopsis*

**DOI:** 10.1101/2024.02.11.579803

**Authors:** Tien Van Vu, Ngan Thi Nguyen, Jihae Kim, Young Jong Song, Thu Hoai Nguyen, Jae-Yean Kim

**Author notes:** These authors contributed equally to this article (Tien Van Vu, Ngan Thi Nguyen). Correspondence: Tien Van Vu; Jae-Yean Kim.

## Abstract

Prime editing (PE) enables almost all types of precise genome editing in animals and plants. It has been successfully adapted to edit several plants at variable efficiency and versatility. However, this technique is inefficient for dicots for unknown reasons. Here, by employing novel combinations of PE components, including an RNA chaperone and modified epegRNAs driven by a PolII-PolIII composite promoter and a viral replicon system, we obtained up to 9.7% of the desired PE efficiency at the callus stage assessed by targeted deep sequencing. Subsequently, we identified that up to 38.2% of transformants contained desired PE alleles in tomatoes and *Arabidopsis*, marking the first successful heritable PE transmission in dicots. Our PE tools also showed high accuracy, specificity, and multiplexing capability, which unlocked the potential for practical PE applications in dicots, paving the way for transformative advancements in plant sciences.

## Introduction

Precise and efficient genome editing is highly demanded for accelerating plant breeding to cope with cropping conditions such as climate changes and arable land shortage^1^. Prime editing (PE) is one of the CRISPR-Cas-based gene editing approaches that allow *in vivo* installation of multiple types of precise DNA changes with low off-targeting activities^2–4^. The PE technique has been successfully applied and improved in monocots^5–13^. However, the PE performance in dicots was extremely low and practically unsuccessful in the whole plant level^14–17^.

Several strategies derived from animal and plant research have been proposed to address the inefficiency and inconsistency of PE. Notable methods include the PE2max configuration, ‘flip and extension’ pegRNA protection, RNase H-truncated RT variant, nucleocapsid RNA chaperone use, Primer Binding Site (PBS) design within an optimal Tm range, and composite promoter use for driving pegRNA transcription^8, 10, 12, 13, 15, 18^. Despite these advancements, the application of PE in dicots, specifically tomatoes, has been challenging due to efficiency issues potentially caused by factors such as self-complementarity of the PBS and spacer sequence, RTT complexity, PE protein stability, unfavorable reaction conditions, and pegRNA stability^4^.

In this study, we have successfully overcome the barriers to PE performance in dicots by combining multiple approaches regarding the PE complex load and favorable conditions for PE reaction, such as a modified epegRNA variant, novel combinations of PE protein components, overproduction of pegRNA by a U6 composite promoter, a geminiviral replicon for amplifying the PE tool, and heat treatment. Our enhanced PE tools resulted in high PE efficiency and inheritability in tomatoes and *Arabidopsis* that has yet to be reported earlier, even all the dicots. Further, we reported that the paired pegRNA approach helped to ensure consistent PE efficiency at various sites, and multiplexed PE approaches showed simultaneous PE activities at several sites in tomatoes. Our data fill in the gap of PE applications in dicots, exploiting the power and versatility of the technology for precision genome editing in plants.

## Results

### Breaking the PE efficiency barrier in tomatoes

We reasoned that the low PE efficiency in dicots might be due to the variation in pegRNA stability with self-complementarity and the reduced catalytic activity of the nCas9(H840)-RT fusion. Both the bottlenecks require a more robust accumulation of active pegRNAs and PE proteins in dicots^4^. We, therefore, combined the following approaches in our initial PE experiment: (1) a nCas9(H840A) of the PE2max variant^3^; (2) an engineered MMLV RT configuration containing the NC and *ΔRNaseH* RT^12^ (Fig. 1a and Extended Data Fig. 1a); (3) a geminiviral replicon cargo^19^ (Extended Data Fig. 1b); (4) an altered version of the epegRNA ^20^; (5) an U6 composite promoter (pU6cm) for enhancing pegRNA transcription^5^ (Fig. 1a and Extended Data Fig. 1a); and (6) a double terminator for the recombined PE protein transcription termination^21^.

**Fig. 1:**
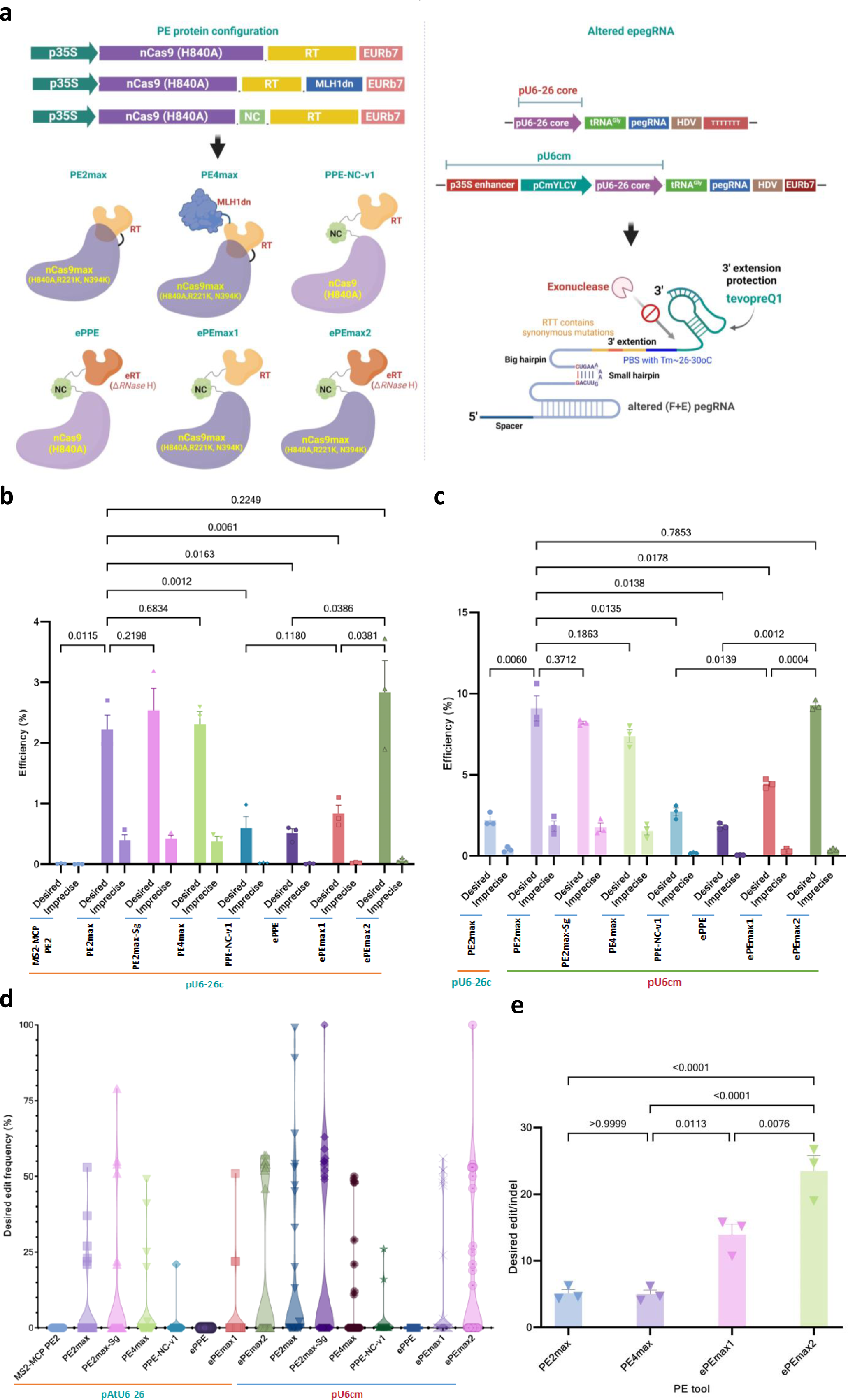
Recombining the PE tool for breaking the efficiency barriers in tomato. **a,** The PE protein configurations (left panel) and altered pegRNA variant (right panel). **b-c,** Performance of the recombined PE tools using pU6-26c (**b**) and pU6cm (**c**) for driving the altered pegRNA transcription at the *SlALS1* site. **d,** Transformants screened by Sanger sequencing and ICE Synthego analysis with the frequency of desired PE edits. **e,** Desired edit/indel ratios.

We conducted experiments using the proposed PE configurations and pegRNAs at the *SlALS1* locus to modify an amino acid (P186S) that conferred herbicide tolerance in plants (Extended Data Fig. 1 and Supplementary Table 1). Using targeted deep sequencing analysis, we found that the novel PE tools returned up to 678-fold higher desired PE efficiency compared to our previous PE system (MS2-MCP PE2)^15^ and reached up to 9.2% with the ePEmax2 protein (Fig. 1b-c and Extended Data Fig. 2). Further, compared to the AtU6-26 core promoter (pU6-26c), pU6cm enhanced 3.2 to 5.3 folds of desired PE efficiency with all the PE proteins tested (Fig. 1c and Supplementary Table 2). Among the PE protein configurations (Fig. 1a), PE2max, PE4max and ePEmax2 resulted in significantly higher desired PE efficiency than that of the PPE-NC-v1, ePPE and ePEmax1 with both the pU6-26c (Fig. 1b) and pU6cm (Fig. 1c). The PE2max using a supportive gRNA (PE2max-Sg) near the pegRNAs did not help to improve PE efficiency (Fig. 1b-c). Similarly, the PE efficiency was not significantly changed when a dominant-negative MLH1 protein was added to the PE4max configuration (Fig. 1b-c). The data also indicates that replacing the nCas9(H840A) of ePPE by nCas9max (H840A, R221K, N394K) to form ePEmax2, 3 times enhanced the desired PE efficiencies (Fig. 1b-c and Supplementary Table 2).

At the plant stages, the frequency of transformants carrying the desired PE edit was 1.1-to 2.7-fold higher using pU6cm compared with pU6-26c (Supplementary Table 3). Up to 38.2 % of analyzed transformants carrying the desired PE allele were obtained with ePEmax2 (with pU6cm), while the same PE protein configuration using the pU6-26c resulted in 33.3 % efficiency (Fig. 1d and Supplementary Table 3), resulting in the lowest fold change (1.1-fold). The other PE configurations evidenced more improvement in PE efficiency. However, only the pU6cm usage generated events containing homozygous PE alleles with PE2max, PE2max-Sg, and ePEmax2 (Fig. 1d, Extended Data Fig. 3a and Supplementary Table 3). Targeted deep sequencing validated most of the edited events obtained by PE2max and ePEmax2 delivered by the replicon carried a high frequency of the desired edit (Extended Data Fig. 3b).

In summary, combinations of novel PE protein configurations and pU6cm successfully broke the PE efficiency barrier in tomatoes via dramatically enhanced PE efficiency and accuracy.

### The major factors that helped break the efficiency barrier in tomato

#### Novel combinations of PE protein components enhanced desired PE efficiency and accuracy

In animal systems, PE2 protein exhibited high PE activity that was frequently associated with moderate levels of imprecise edits^9, 20, 22, 23^. We observed similar levels of imprecise PE edits (Fig. 1c and Supplementary Table 2) and desired/imprecise edit ratios (Fig. 1e) using the PE2max, PE2max-Sg, and PE4max (Fig. 1e). The majority of the imprecise PE edits were the incorporation of one to several nucleotides of the gRNA scaffold sequence adjacent to the RTT that was also reported in animals^22^ and plants^9^. However, the novel combinations of the PE components in ePEmax2 maintained high activity of PE reaction while dramatically reducing the imprecise frequency (Fig. 1b-c), and thus, its desired/imprecise PE ratio was much higher compared with the PEmax variants (Fig. 1e). Therefore, the ePEmax2 can be employed for editing reactions that favor high precision over the total PE activity.

#### The replicon-based delivery dramatically enhanced PE efficiency

Earlier, we constructed a Bean Yellow Dwarf Virus (BeYDV) replicon system that autonomously amplified its cargo DNAs to enhance gene targeting efficiency in tomatoes^19^. Here, the replicon system amplified the PE tool-encoding DNA, resulting in the breakthrough outcomes of PE performance in tomatoes (Fig. 1b-c). The replicon system resulted in 6.6- to 7.8-fold higher desired PE efficiency than the T-DNA, with the highest enhancement for the ePEmax2 protein configuration (Fig. 2a and Supplementary Table 4). This indicates that the higher expression was one of the key factors for breaking the efficiency barrier in the tomatoes.

**Fig. 2:**
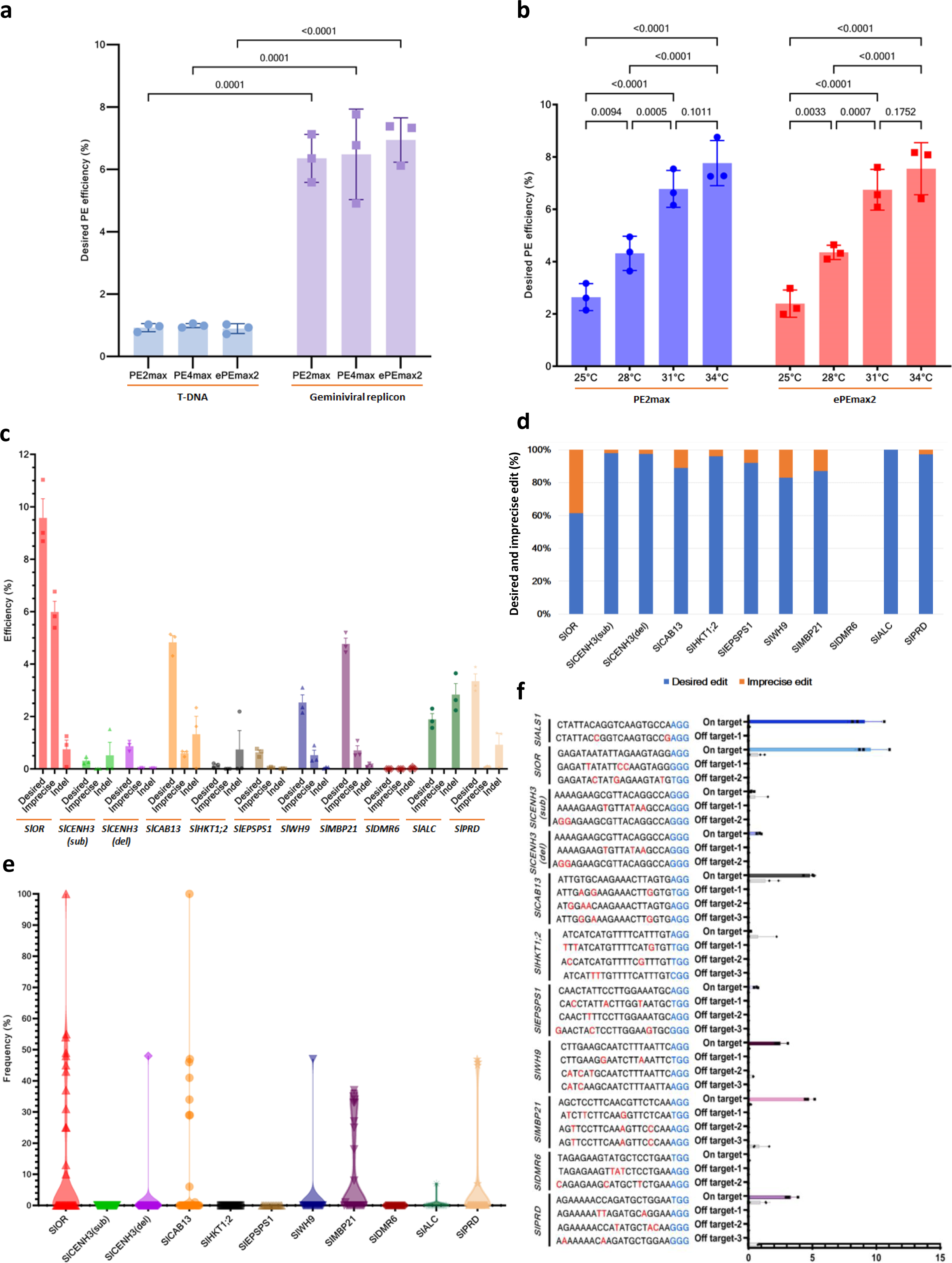
Impacts of the delivery and heat treatment on PE performance at the *SlALS1* and other targets. **a-b,** Comparison of T-DNA and the geminiviral replicon (**a**) and heat treatments (**b**) at the *SlALS1* site using the PE2max-based tool. **c-d,** Callus-stage PE efficiency revealed by targeted deep sequencing (**c**) and the portions of desired and imprecise edit at ten tomato loci (**d**). **e,** PE events and desired PE frequency obtained at various tomato loci. **f,** Off-target activity of the enhanced PE tool.

#### Heat treatment significantly improved PE activity

It is well-known that heat treatment could significantly enhance the cleavage activity of the CRISPR-Cas9^24^ and Cas12a^19, 25^ in dicots. We therefore conducted the experiment at 25, 28, 31, and 34°C using the PE2max- and ePEmax2-based PE tools. The desired PE efficiency was gradually significantly enhanced when the temperature was increased from 25 to 34°C (Fig. 2b and Supplementary Table 5). Compared to 25°C, the desired PE efficiency was improved 2.9- and 3.2-fold at 34°C with PE2max and ePEmax2, respectively (Supplementary Table 5).

### The PE tools worked robustly and precisely at various loci in tomatoes

To exploit the applicability and versatility of the enhanced PE tools developed, we employed the PE2max configuration to precisely introduce various edit types at 11 sites of 10 loci that are potentially agronomically significant in tomatoes (Supplementary Table 1). Except for the *SlDMR6*, all the tested sites exhibited desired PE edits with efficiency ranges from 0.1 (*SlHKT1*;2) to 9.6% (*SlOR*) (Fig. 2c and Supplementary Table 6). Five of eleven sites showed less than 1% desired PE efficiency at the callus stage (Fig. 2c and Supplementary Table 6). At most sites, the proportion of desired edits was much higher than the imprecise edit (Fig. 2d). Only the SlOR site appeared to be the least precise (Fig. 2d and Supplementary Table 6).

As a result, analyzing fewer than 50 transformants revealed desired PE events from 7/11 sites (Fig. 2e and Table 1). The PE T0 events carried homozygous (ho), heterozygous (he), or chimeric (chi) edited alleles (Extended Data Fig. 4a, b and Table 1). We obtained homozygously edited events in the first generation of transformants at the *SlOR* and *SlCAB13* loci, indicating the PE tool worked robustly at these sites. Indeed, validating the edited events by targeted deep sequencing revealed desired base changes in most of the events (Extended Data Fig. 4c). However, some events were also associated with high frequency of scaffold sequence incorporations (imprecise (Imp) edits) at the *SlOR* (3 events out of 13) and *SlMBP21* (1/8) sites (Extended Data Fig. 4c). The relatively imprecise frequency at the SlOR site correlated with the high imprecise frequency at the callus stage (Fig. 2c and Supplementary Table 6). Our observation at various tomato loci led to a conclusion that despite working well at the majority of the tested loci, the performance of the PE tool was locus-dependent (Fig. 3, Table 1 and Supplementary Table 6), as observed earlier^4, 9, 22^ and thus, more improvement of the PE tool is required for the PE-inactive sites.

**Fig. 3:**
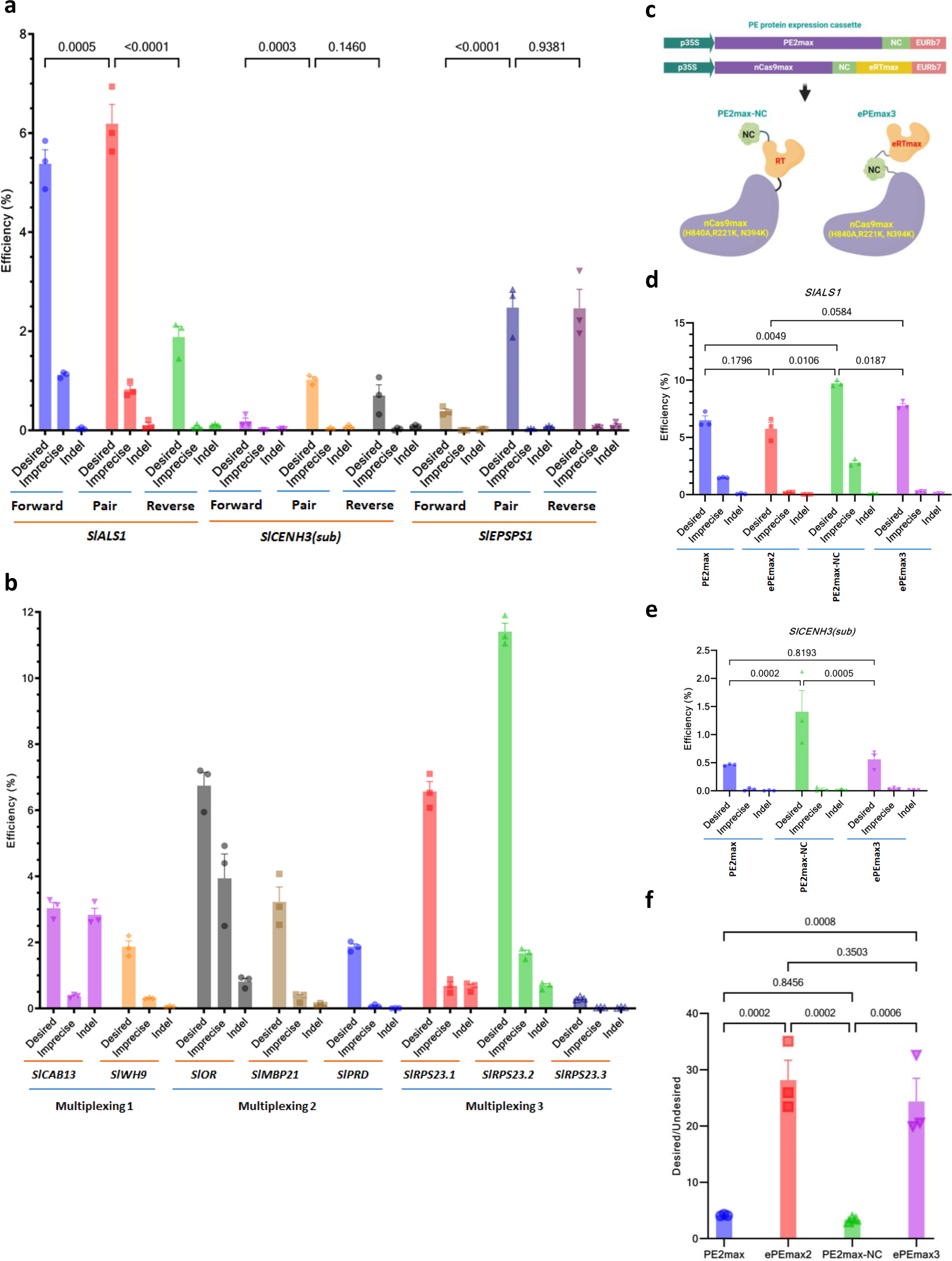
PE performance using paired pegRNAs, multiplexing, and the NC RNA chaperone. **a,** The paired pegRNA approach. **b,** Multiplexing PE using several pegRNAs or single pegRNA**. c-f,** improving the PE tools using the NC RNA chaperone.

**Table 1.**
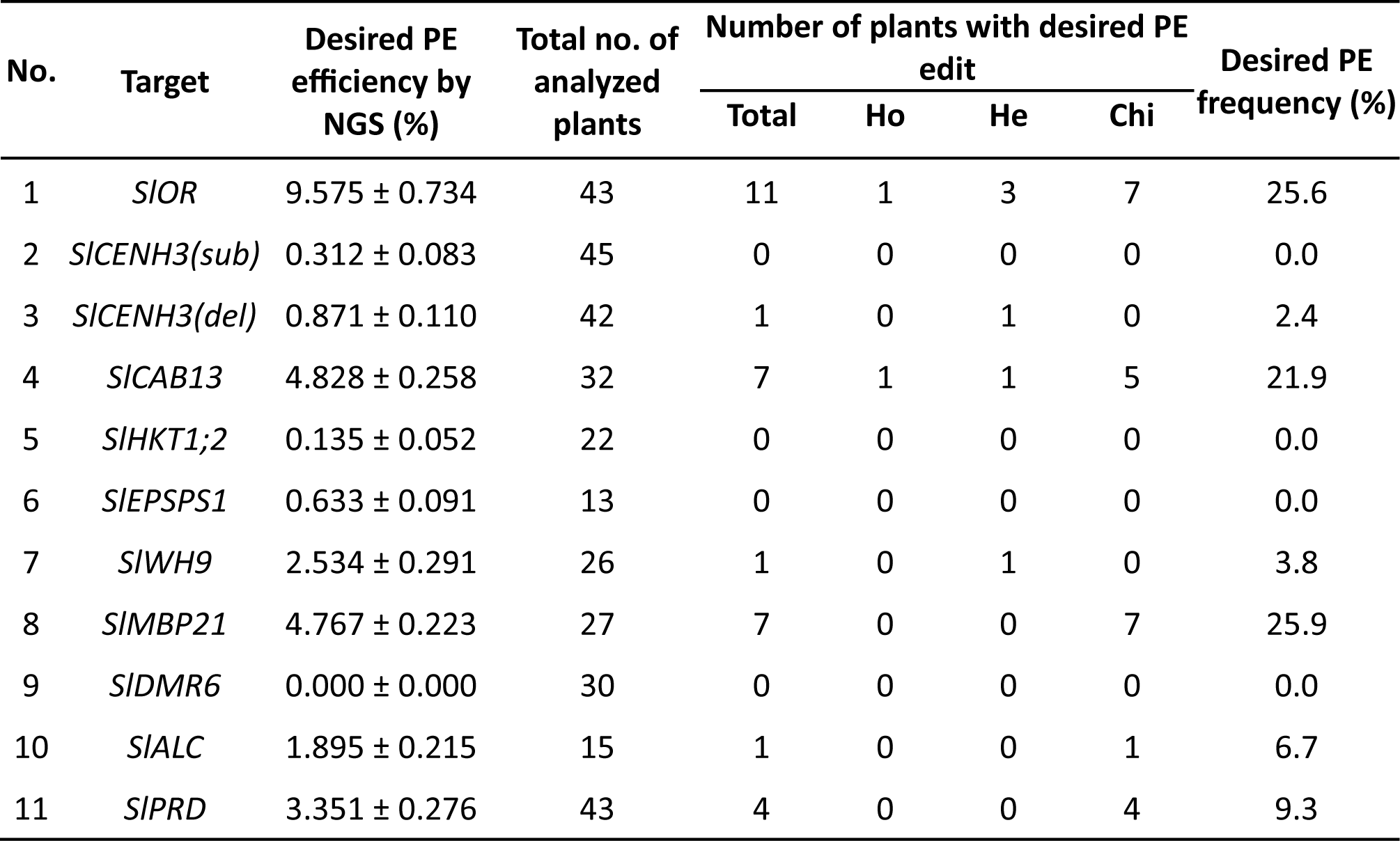
The PE performance on various tomato targets at the plant stage.

### The enhanced PE tools did not have significant off-target activities

One important characteristic of the PE tool is the low off-target activity thanks to the absence of double-stranded break (DSB) formation^9, 22, 26^. The activity of the PE tools at the potential off-target sites (Supplementary Table 7) was analyzed by targeted deep sequencing. We could not find any significant modifications among the potential off-target sites of the PE-transformed explants compared with WT samples (Fig. 2f and Supplementary Table 8), indicating the PE tool is highly precise.

### The paired pegRNA use assured the highest PE efficiency in tomatoes

Using a pair of pegRNAs to install desired edits at a site might help to avoid the flap competition step, thereby enhancing PE efficiency in plants^4, 8^. We tested if the dual-pegRNA strategy could improve desired PE efficiency in tomatoes at the *SlALS1*, *SlEPSPS1,* and *SlCENH3(sub)* sites. The PE2max protein was used for the test (Fig. 1a). The desired PE efficiency was the highest with paired pegRNAs at all the tested sites while the imprecise products were slightly reduced (Fig. 3a and Supplementary Tables 9). In each tested site, one pegRNA demonstrated significantly lower efficiency than its counterpart. This observation suggests that the dual-pegRNA approach might combine the activities of individual pegRNAs, potentially ensuring the highest achievable PE efficiency at each targeted site.

### Multiplexed PE is feasible with the PE2max in tomatoes

We assessed the capacity of multiplexing in tomatoes for simultaneous editing of two (*SlCAB13* and *SlWH9*) and three (*SlOR*, *SlMBP21,* and *SlPRD*) sites (Supplementary Table 1). A single pegRNA was also used to simultaneously edit three tomato homologs of the ribosomal protein S23 (*RPS23*) gene for archiving a conservative allele (K60R, corresponding to K59R in tomatoes) that potentially enhances the fidelity of protein translation^27^ (Supplementary Table 1). We found that the multiplexed PE tools worked well at all the targeted sites, albeit the desired PE efficiency was slightly reduced (Fig. 3b and Supplementary Table 10) compared with each of the single pegRNA constructs (Fig. 2c). Interestingly, using single pegRNA, we could simultaneously edit three homologs of the *SlRPS23* gene. The desired PE efficiency reached 6.6, 11.4, and 0.3% at *SlRPS23.1*, *SlRPS23.2,* and *SlRPS23.3 sites*, respectively (Supplementary Table 10).

### The PE protein configurations could be further improved by the RNA chaperone

Though the barriers to PE efficiency have been successfully overcome and reached the levels observed in monocots (summarized by Vu et al.^4^), the locus dependence and low efficiency at some tested loci demand further improvement of the current PE tools. In the previous experiment, using NC in the ePEmax1 and ePEmax2 protein configurations efficiently improves the PE performance over the PPE-NC-v1 and ePPE, respectively (Fig. 1b-c). We investigated whether adding the NC to the PE2max configuration and reconstructing the ePEmax2 with the PE2max-based ΔRNaseH RT codons (eRTmax), named ePEmax3 (Fig. 3c), would enhance the tools further. Intriguingly, adding NC enhanced 1.5 folds of desired PE efficiency at the *SlALS1* site. Similarly, ePEmax3 improved 1.4 folds of desired PE efficiency at the same site compared to ePEmax2 (Fig. 3d and Supplementary Table 11). The enhancement was higher, 3.1-fold, using PE2max-NC at the inactive site (*SlCENH3(del)*), Supplementary Table 1) than PE2max (Fig. 3e and Supplementary Table 11). At the *SlCENH3(del)* site, PE2max-NC also slightly enhanced the desired PE efficiency compared to PE2max. The desired PE efficiency reached up to 9.7% and 1.4% at the *SlALS1* and *SlCENH3(del)*, respectively. Even though the PE activity was significantly enhanced, the accuracy of the PE2max-NC and ePEmax3 was not significantly altered at either of the sites (Fig. 3f and Supplementary Table 11).

### The newly developed PE tools worked efficiently in *Arabidopsis*

The exploration of PE in dicots has been limited in several species because of their inefficiency^4^. We asked if our enhanced PE tools could be employed to improve PE performance in *Arabidopsis*, which exhibited low PE efficiency earlier^17^. Using the PE2max-NC variant with the p35S-tHSP promoter-terminator cassette (Fig. 4a), we constructed PE tools to target *AtALS* and *AtPDS3* (Supplementary Table 12). We first conducted PE experiments with the optimal temperature (22°C) for *Arabidopsis* growth and analyzed T1 plants by targeted deep sequencing using pooled samples that revealed desired PE edits at both the sites with stronger activity at the *AtPDS3* (Supplementary Table 13). Analyzing individual T1 plants of the *AtPDS3* constructs obtained three events carrying more than 10% desired edits out of 96 plants (Fig. 4b and Extended Data Fig. 5). This indicates that PE events carrying desired edits could be obtained in *Arabidopsis* at the normal growth conditions. Subsequently, we employed heat treatment (Fig. 4c) to improve the PE performance up to seven folds, up to 21.7%, at the *AtPDS3* sites (Fig. 4b and Extended Data Fig. 5). These data indicate that our enhanced PE tool could be efficiently applicable to other dicots.

**Fig. 4:**
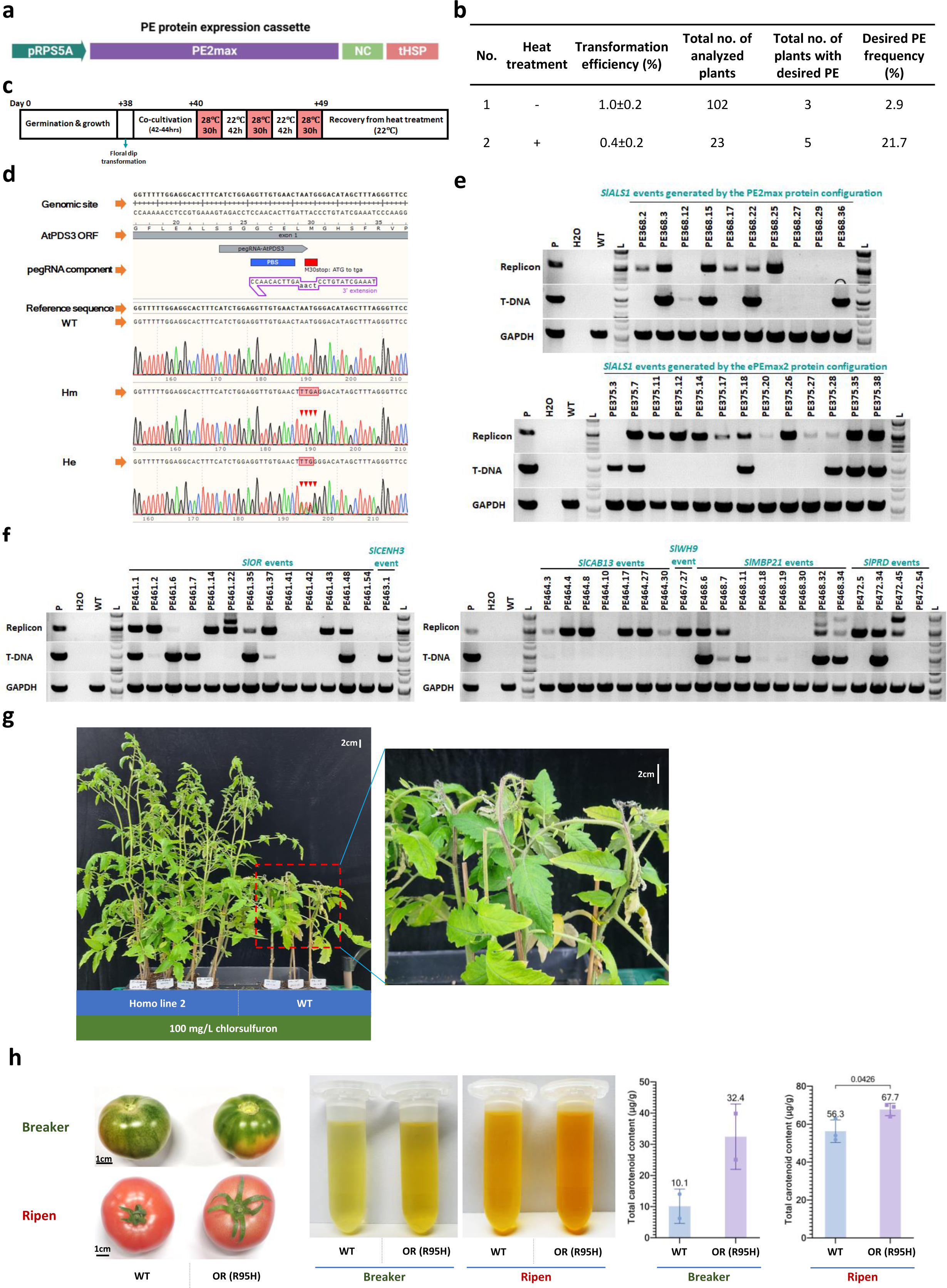
Application of the enhanced PE tool in *Arabidopsis* and characterizing the PE-edited alleles. **a-d,** Design and employment of the enhanced PE tool in *Arabidopsis.* **a,** The expression cassette of the PE protein used for *Arabidopsis* editing. **b,** The PE efficiency obtained with and without heat treatment. **c,** The heat treatment conditions. **d,** Chromatogram alignment showing the T2 lines carrying homozygous (Ho) and heterozygous (He) *AtPDS3* KO allele compared with WT. **e-f,** Assessment of the presence of T-DNA and replicon in the T0 events. **g,** Herbicide tolerance of the T1 plants carrying homozygous *SlALS1* (P186S) allele. **h,** The enhancement of total carotenoid production in tomato carrying homozygous *SlOR* (R95H) allele.

### Transgene-free PE-edited plants could be obtained in the first and next generations of the transformants

Editing without remaining the editing tools within the cells is important for plant breeding^28, 29^. We analyzed the occurrence of the T-DNA and replicon in some representative PE events obtained from the study by PCR specified for the T-DNA and circularized form of replicons released from the binary plasmids (Extended Data Fig. 1b). Initially, the PE events obtained by using the PEmax and ePEmax2 at the *SlALS1* site were checked to reveal the occurrence of either T-DNA or replicon in most of the plants (Fig. 4e). Interestingly, we found two events (PE368.27 and 29) of the PEmax-based tool without a T-DNA or replicon band, representing for 8.7% frequency among a total of 23 analyzed plants (Fig. 4e). T-DNA and replicon-free plants could later be obtained from offspring of the events containing replicon (PE368.17) or both the T-DNA and replicon (PE375.18) (Extended Data Fig. 6a). Subsequently, we assessed thirty-four T0 events obtained from other targeted sites/loci *(SlOR*, *SlCENH3*, *SlCAB13*, *SlWH9*, *SlMBP21* and *SlPRD*) using the PE2max-based tool and found several PE events (PE461.41, PE461.42, PE461.54, PE464.10, PE468.30 and PE472.54) that did not contain a band of T-DNA and replicon (Fig. 4f), which averages at 17.6% frequency. T-DNA and replicon-free plants were subsequently obtained in the next generation of the T0 events (Extended Data Fig. 6b).

### The PE alleles were stably inherited and conferred dominant phenotypic appearance

To assess the inheritance of the desired PE edits in the next generation of the PE events, we self-pollinated the plants, harvested and germinated the seeds, and analyzed seedlings for calculating the desired edit frequency. We found that all the desired edits were stably inherited in the next generation in tomato (Extended Data Fig. 7a). Using the tomato seedlings carrying homozygous *SlALS1* (P186S) allele for chlorsulfuron tolerance tests, we found that they were resistant against up to 100 mg/L chlorsulfuron herbicide spraying while all the WT apical shoots were dead two weeks’ post-spraying (Fig. 4g and Extended Data Fig. 7a). In addition, fruits of the tomato event carrying homozygous alleles of *SlOR* (R95H) produced higher total carotenoid contents compared with WT controls at both the breaker and ripen stages (Fig. 4d). In *Arabidopsis*, the PE-installed knockout allele of the *AtPDS3* gene was stably inherited in the T2 generation (Fig. 4d and Supplementary Table 14) and the inheritance of a homogeneous pair (Ho) of the *AtPDS3* knockout alleles resulted in albino phenotypes (Extended Data Fig. 7c).

## Discussion

The PE is a breakthrough technique for precision gene editing and modern plant breeding^26^. However, its applications have been limited in dicots due to the efficiency barrier^4^. We employed synthetic combinations of PE components developed in mammals and monocots to generate novel PE tools that could break the efficiency barrier in tomatoes (Fig. 1) and *Arabidopsis* (Fig. 4a-d) and achieved high efficiencies of inheritable desired PE edits.

Here, compared to our previous PE system (MS2-MCP PE2)^15^, we obtained a dramatical improvement of the PE efficiency up to 678 folds using modified PE tools with U6cm promoter-driven modified epegRNA expression (Fig. 1). The BeYDV replicon system was well-known for the possibility of enhancing up to 80-fold mRNA transcription and up to 10-fold protein translation in plants^30^. It was efficiently used to support our CRISPR-Cas-based gene targeting applications in tomatoes^19, 31^. We obtained dramatical improvement of the PE efficiency up to 7.9 folds using the replicon from the T-DNA baseline (Fig. 2a), indicating that the amplification strengths of the replicon might maximize the transcription of the modified epegRNAs and translation of the PE proteins.

Our results show that by adding NC to the PE2max configuration and reconstructing ePEmax2 with PE2max-based ΔRNaseH RT codons (eRTmax), we can significantly enhance the efficiency and accuracy of PE (Fig. 3c-f). Recent studies have shown that including nucleocapsid (NC) between nCas9(H840A) and RT domains significantly enhances the PE process in monocots, providing a foundation for the enhancement of our PE tools^12^. Subsequently, our enhanced PE tool exhibited efficient editing in *Arabidopsis*, which was also enhanced seven folds with the heat treatment (Fig. 4a-d), albeit the tool could be further improved at the other loci, e. g., using the paired pegRNA approach.

Interestingly, our results also showed that these enhanced PE tools could generate transgene-free PE-edited plants in the first generation of transformants (Fig. 4e, f). This is a crucial aspect of plant breeding as it eliminates concerns about foreign DNA integration and potential regulatory issues. The fact that an average of 14% transgene-free plants were obtained in the first generation (Fig. 4e, f) and segregated plants in the next generation (Extended Data Fig. 6) suggests that the improved PE tools can be effectively utilized in plant breeding programs, although this finding requires further validation in other plants.

Our study also evidences that the desirable PE edits were stably inherited and showed strong phenotypic performance in herbicide tolerance (Fig. 4g and Extended Data Fig. 7b), enhanced carotenoid production in tomato (Fig. 4h), and impaired chlorophyll and carotenoid production in *Arabidopsis* (Extended Data Fig. 7c), signifying the reliability and usefulness of these tools in generating inheritable modifications.

In conclusion, our research presents significant improvements to the PE tools in dicot plants, yielding enhanced efficiency and accuracy, transgene-free plants, and stably inherited edits. These advancements hold great promise for the future of plant gene editing, potentially enabling more precise and efficient modifications, even in species where conventional editing tools have been less effective. Further studies are needed to validate these tools in other plant species, including monocot plants, and investigate their applicability in various agricultural and horticultural contexts.

## Online Methods

### Combinations of PE protein components and construction of plasmids

For combining the bioparts to generate PE proteins tested in this study, we cloned PE2max and PE4max, the nCas9(H840A) of ePEmax1, ePEmax2, and the nCas9 and eRT of ePEmax3 from the plasmid pCMV-PEmax-P2A-hMLH1dn (Addgene #174828). The nCas9 of PPE -NC-v1 was cloned from nCas9-PPE plasmid (Addgene #140445), and its NC and RT were cloned from pH-ePPE plasmid (Addgene #183097). The ePPE was cloned from pH-ePPE plasmid. The NC sequence used in ePEmax1, ePEmax2, ePEmax3, and PE2max-NC was cloned from the pH-ePPE plasmid. For the basic PE expression cassettes, a CaMV 35S promoter (p35S) (Addgene #50267) and an EU + Rb7 double terminator (EURb7)^21^ (Supplementary sequences) were used to control the transcription.

The epegRNA (F+E SpCas9 scaffold with a tevopreQ1 pseudoknot for 3’ terminal protection^20^) was modified by introducing the base changes from the altered SpCas9 scaffold^32^ (Supplementary sequences). The altered epegRNA was placed between a tRNA(gly) and an HDV ribozyme sequences^13^. For driving the transcription of the pegRNAs, either the pU6-26c or pU6cm promoter^13^ was used (Supplementary sequences). The edited sites/loci, gRNA, and RTT information were selected and detailed in Supplementary Table 1. The pegRNAs used in pairs were designated as “Forward” and “Reverse” pegRNAs (Supplementary Table 1). The pegRNAs were amplified by PCRs using the primers listed in Supplementary Table 14, cloned into the expression cassettes, and combined into the binary vectors (Extended Data Fig. 1) by Golden Gate assembly.

For the T-DNA-based PE tools, the expression cassettes of the selection marker (NptII, Addgene #51144), PE protein, and pegRNA were cloned into the pAGM4723 plasmid (Addgene #48015). When the geminiviral replicon system was used, the expression cassettes were cloned into the pLSL.R.Ly vector reported earlier^19^.

All the bioparts (Supplementary sequences) were domesticated into the Moclo^33^ level 0 plasmids and assembled into the PE protein combinations and binary vectors, as shown in Extended Data Fig. 1.

### *Agrobacterium*-mediated tomato transformation and assessment of PE efficiency

*Agrobacterium*-mediated transformation of tomato was carried out following our in-house protocol as detailed by Vu and coworkers^19^. The *Agrobacterium tumefaciens* strains used in this study were EHA105-based super *Agrobacterium* version 2 (EHA105spv2)^34^. In brief, seven-day-old cotyledons were excised and utilized for the transformation. The agrobacteria harboring the PE constructs were cultured, harvested by centrifugation, and subsequently resuspended in an ABM-MS solution^19^, supplemented with 200 µM acetosyringone to reach an optical density at 600 nm (OD_600_) of 0.8. The agrobacteria were activated by incubating at 28°C for one hour before transformation. The agrobacteria and cotyledon explants were mixed and incubated for 20 minutes at room temperature. Then, the transformed explants were co-cultivated for 2 days before washing and transferred onto the selection medium. The samples were incubated at 31 °C for 5 days and then 28 °C for 5 days before applying 25 °C incubation for the rest of the stages. Regenerated shoots were selected on a medium containing 80 mg/L kanamycin and rooted before transferring onto the soil. Subsequently, the hardened plants were assessed for PE activity. For temperature treatment experiments, after cocultivation, the explants were incubated at different temperatures for 5 days and then incubated at 28°C for 5 days before transferring to 25°C incubation for the later stages.

To evaluate the efficiency of PE, samples were collected at 16 days post-transformation (dpt) and subjected to targeted deep sequencing. At the plant stage, the leaves of transformants were gathered and screened for PE alleles using PCRs and Sanger sequencing. Targeted deep sequencing analysis was performed to validate some representative events carrying PE alleles.

### *Arabidopsis* transformation by the floral dip method

The floral dip method, with some minor adjustments, was used to transform *Arabidopsis thaliana* accession Col-0 plants. Germinated plants were grown under long-day conditions, with 16 hours of light and 8 hours of darkness. A. tumefaciens cells carrying PE constructs were grown overnight, harvested by centrifugation at 4000 rpm for 10 minutes, and then suspended in a 5% sucrose solution that contained 200 µM acetosyringone (Duchefa Biochemie, Haarlem, The Netherlands) and 0.02% of silwett L-77. At 38 days old, plants were transformed by dipping their floral parts into the A. tumefaciens suspension for ten seconds, three times, followed by co-cultivation in the dark at 22°C for 24 hours. After co-cultivation, plants were incubated at 28°C for 30 hours and then recovered at 22°C for 42 hours. This heat treatment was repeated two more times before the plants were grown at 22°C under long-day conditions (16 h light/8 h dark) until seed harvest (Fig. 4c). Transgenic T1 plants were chosen on half-strength MS agar plates supplemented with 1% sucrose, 100 mg/L of timentin, and 50 mg/L of kanamycin. The seeds harvested from edited plants carrying desired PE alleles were germinated on half-strength MS agar plates supplemented with 1% sucrose and 100 mg/L of timentin without additional antibiotic selections.

### Targeted deep sequencing

Genomic DNA (gDNA) was extracted from cotyledon explants or plant leaves using the CTAB method. The Illumina MiniSeq sequencing service (KAIST Bio-Core Center, Daejeon, Korea) was employed for the analysis. MiniSeq samples were prepared through three PCRs utilized primers listed in Supplementary Table 15. The third PCR was conducted with the manufacturer’s provided primers to assign sample IDs. The MiniSeq raw data FASTQ files were subsequently analyzed using the Cas-Analyzer^35^ and CRISPResso2^36^ tools with the parameters listed in Supplementary Table 16.

### Identification of transformants carrying PE alleles

Transformants that survived after the hardening step were subjected to screen for PE alleles. Three leaf fragments were collected from three different compound leaves of each plant and combined to isolate gDNA using the CTAB method. PCR reactions were employed for amplifying DNA sequences flanking the targeted sites, and PCR products were sequenced by the Sanger method. The resulting sequencing chromatograms were analyzed using the ICE Synthego tool^37^ to screen for potential events carrying PE alleles. Some representative PE events were subsequently validated by targeted deep sequencing.

### Assessment of T-DNA and replicon presence within PE events

To assess the presence or absence of T-DNA and replicon, gDNAs of PE plants were used as the template for PCR reactions using primer pairs specified to the right border (RB) of the T-DNA and the circularized forms of replicon (Supplementary Table 15). The replicon primers were designed only to amplify circularized DNA that was constructed to be released from the vectors^19^. PCR products were resolved on 1% agarose gel, and the absence of T-DNA and replicon was determined based on the absence of PCR bands at the sizes corresponding to T-DNA and replicon, respectively.

### Potential off-target analysis

The gRNA sequences of the pegRNAs (Supplementary Table 1) were subjected to the Cas-OFFinder tool for searching for potential off-target sites within the tomato genome database *Solanum lycopersicum* (SL2.4) that contain fewer than 4 mismatches to the gRNA sequences. The 16-dpt cotyledon explants that were transformed with pegRNAs having identified potential off-target sites (Supplementary Table 7) were subjected to analysis by targeted deep sequencing using primers listed in Supplementary Table 15.

### Herbicide resistance assay

Eight-leaf stage tomato seedlings carrying homozygous *SlALS1* (P186S) and WT were sprayed until fully wetted two times (a five-minute interval) with 0, 50, and 100 mg/L chlorsulfuron (Sigma-Aldrich, Massachusetts, US, CAS No.:64902-72-3) solution containing SilwettL-77 (0.02%). Sprayed plants were photographed two weeks post-spraying.

### Assessment of carotenoid contents in tomato fruits

Fresh fruits were collected at the breaker and ripening stages and sampled for assessing the carotenoid contents. The carotenoid assessment was conducted using UV-Vis spectroscopy described by Lichtenthaler and Buschmann^38^. One gram of fresh fruit tissue was ground to a fine powder in liquid nitrogen and 10 ml of acetone was added. The mixture was then transferred into a 15 ml falcon tube and vortexed in two minutes. Subsequently, the sample was centrifuged at 5000 rpm for 5 minutes, and 5 ml of the upper phase was transferred into a new tube to measure absorption spectra (A662, A645, and A470nm). The content of total carotenoids was calculated as the following:

Chlorophyll a: C_a_ = 11.24*A662 – 2.04*A645 (μg/ml)

Chlorophyll b: C_b_ = 20.13*A645 – 4.19*A662 (μg/ml)

Total carotenoids: C_x+c_ = 10*(1000*A470 – 1.90*C_a_ – 63.14*C_b_)/214 (μg/g)

### Data analysis and presentation

All the experiments were carried out at least in triplicates. Editing data, as well as any applicable statistical analyses and scatter plots, were subsequently processed using MS Excel and the GraphPad Prism 9.0 software. Multiple comparisons were executed utilizing the uncorrected Fisher’s LSD test.

## Author contributions

T.V.V. and J.Y.K. conceived and designed the research. T.V.V., N.T.N, J.K., S.Y.J., and T.H.N. conducted experiments. T.V.V., N.T.N. and J.Y.K. analyzed data. T.V.V. and N.T.N. wrote the manuscript. T.V.V. and J.Y.K. finalized the manuscript. All authors read and approved the manuscript.

## Extended Data Figure legends

**Extended Data Fig. 1:**
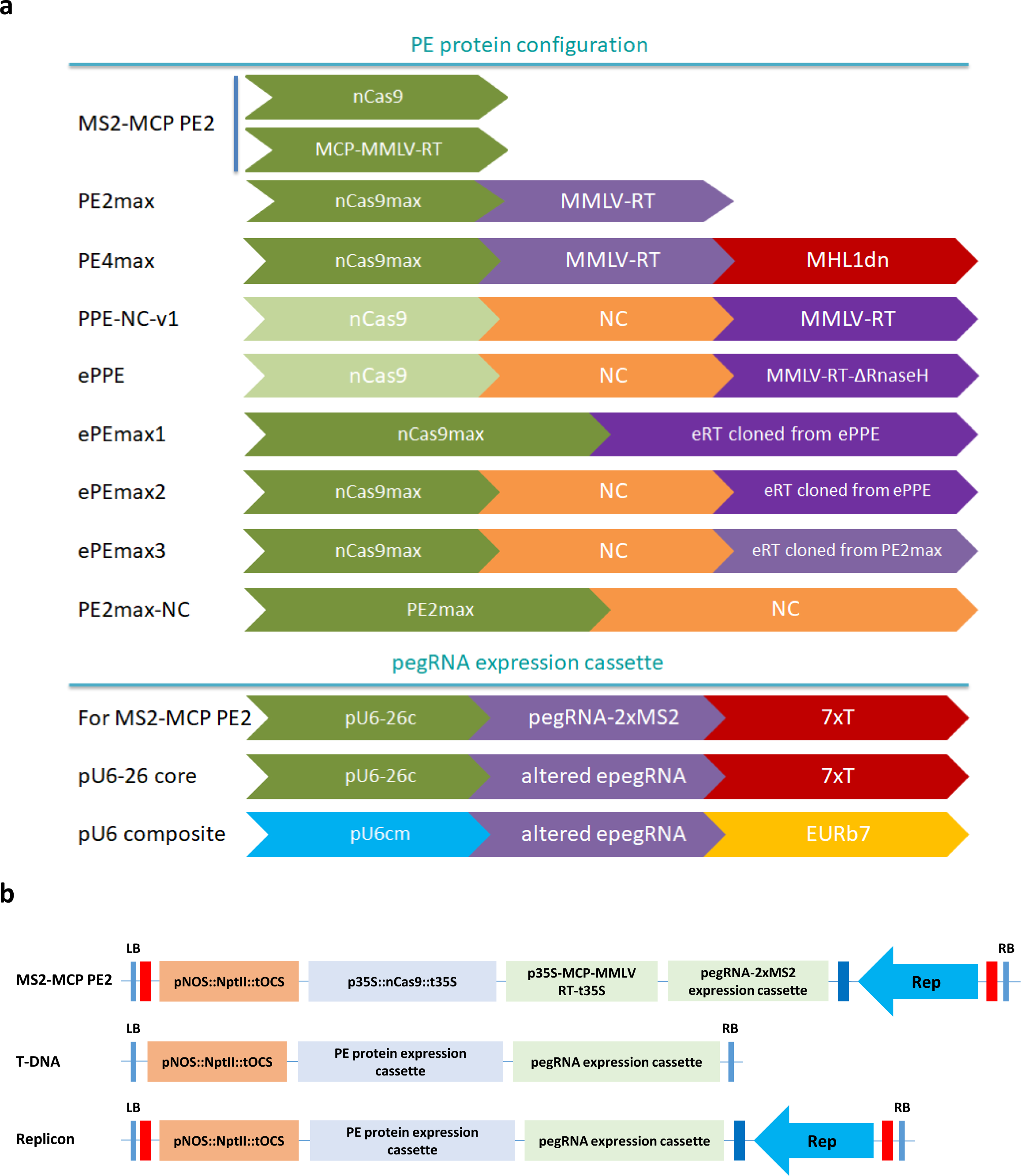
Construct design of the recombined PE tools. **a,** The arrangement of the PE protein components (upper panel) and pegRNA expression cassettes (bottom panel). The bioparts with the same color codes were cloned from the same sources of DNA materials. **b,** Binary vector maps of the T-DNA and geminiviral replicon tools.

**Extended Data Fig. 2:**
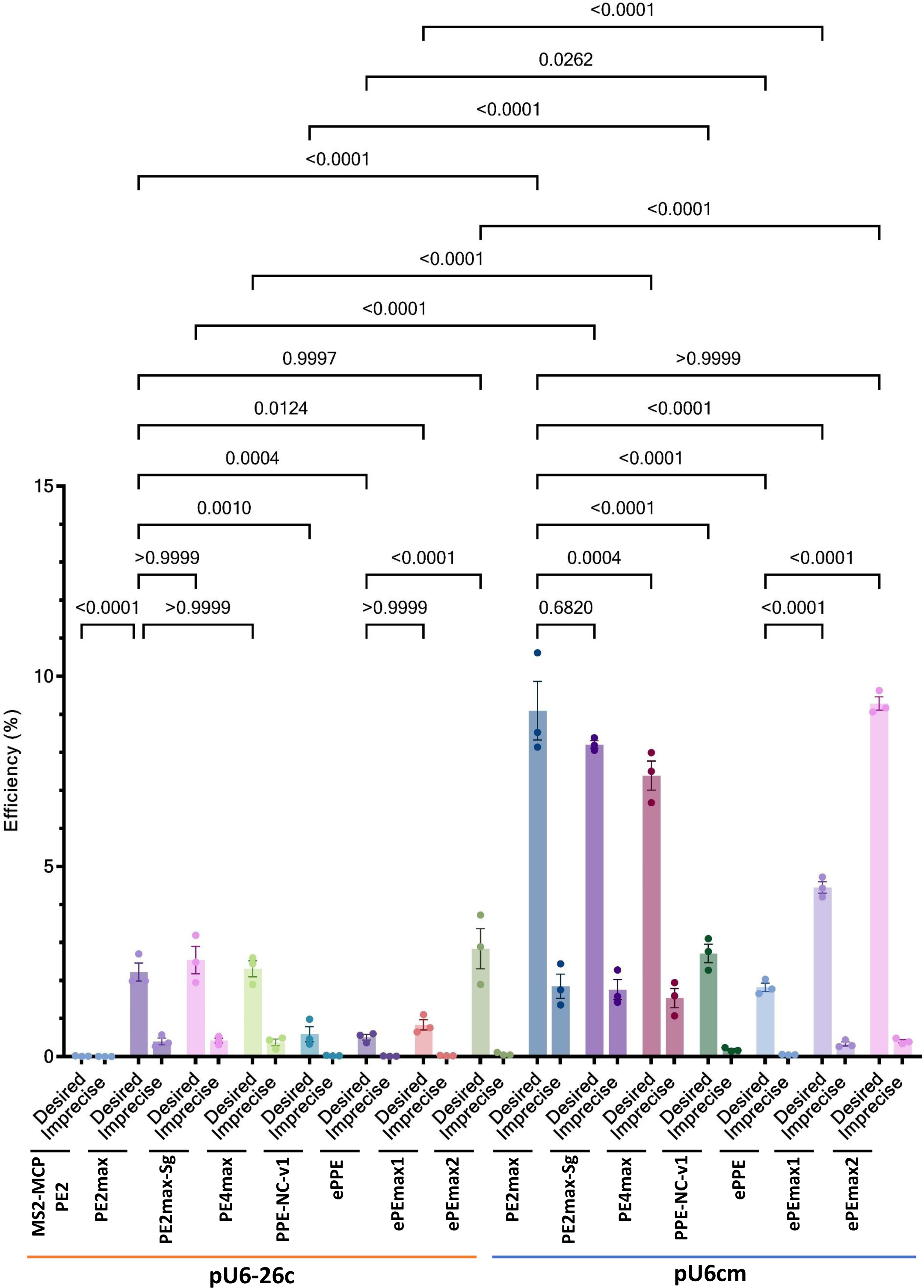
Callus-stage PE efficiency of the recombined PE tools with the pU6-26c and pU6cm obtained at the *SlALS1* site. All the constructs were transformed into thin-sliced cotyledon explants in triplicates, and 16-dpt samples were analyzed by targeted deep sequencing. The statistical analysis and plotting were performed using GraphPad Prism 9.0.

**Extended Data Fig. 3:**
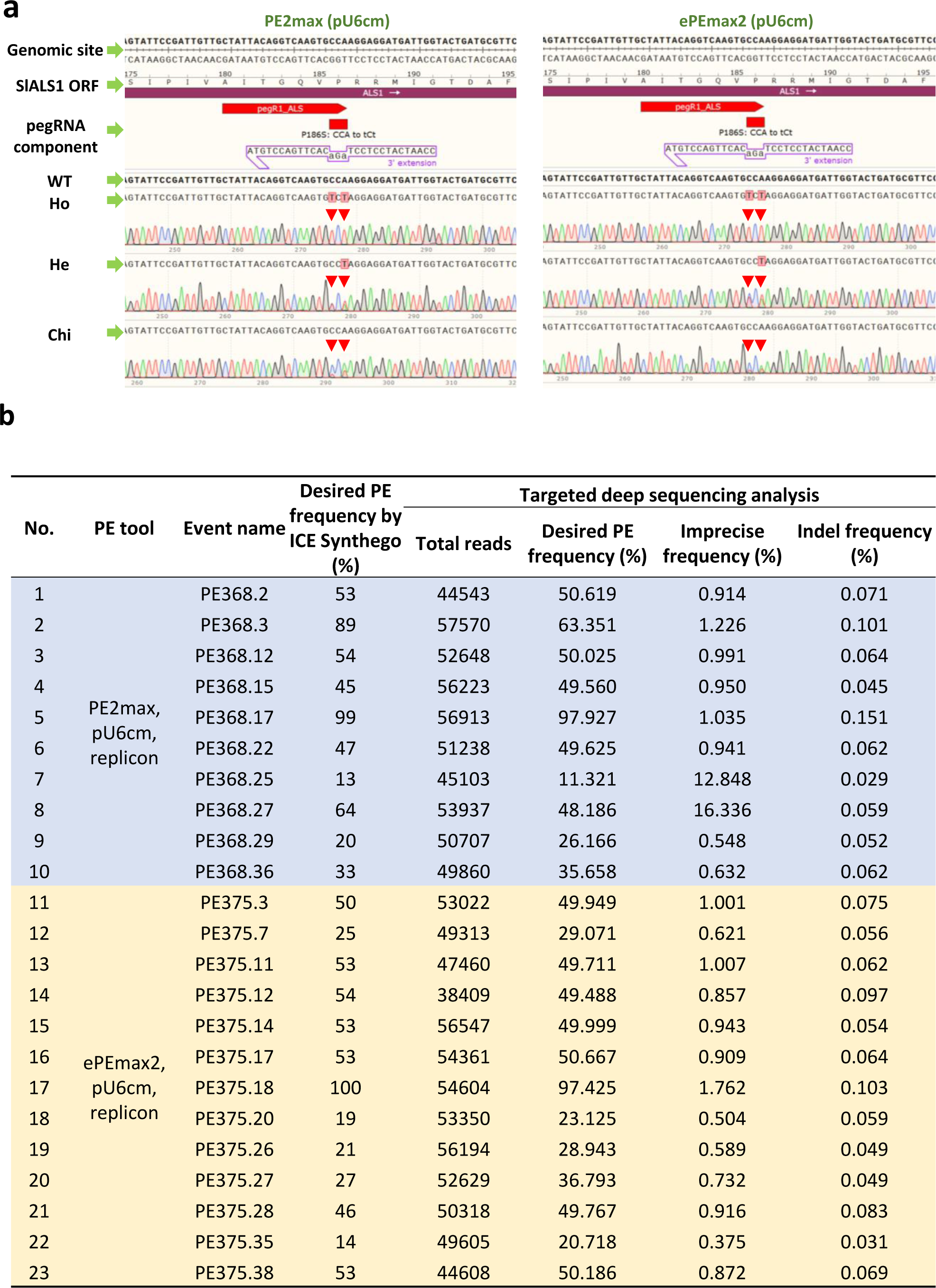
T0 events carrying desired edit (P186S) obtained at the *SlALS1* site. **a,** Alignments of Sanger sequencing chromatograms among the PE events carrying homozygous (Ho), heterozygous (He), and chimeric (Chi) forms of the desired edit generated by the PE2max and ePEmax2-based constructs. The red arrows denote the peaks containing desired base changes. **b,** Targeted deep sequencing analysis of the PE events obtained using the PE2max and ePEmax2-based constructs.

**Extended Data Fig. 4:**
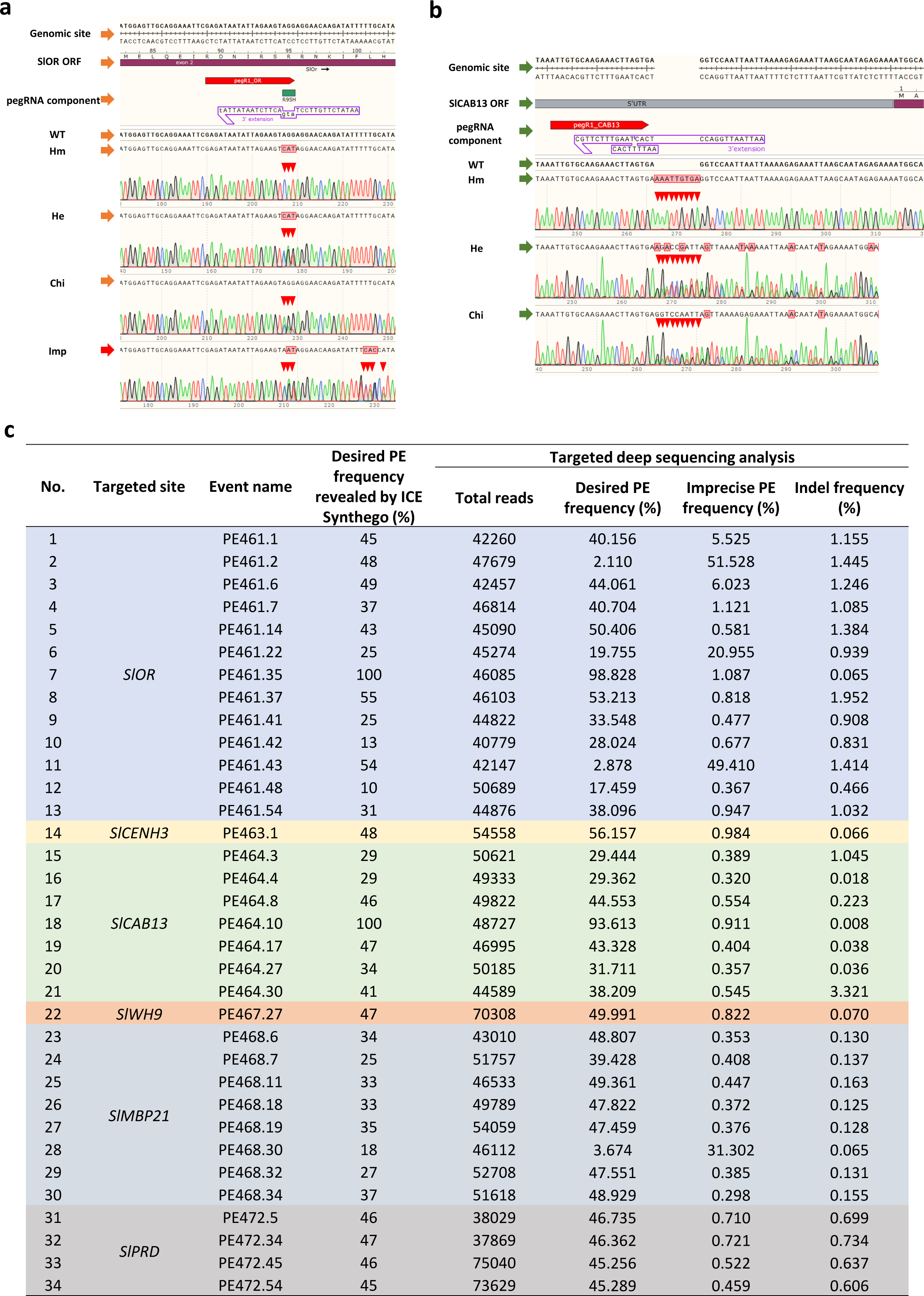
T0 events carrying desired edits obtained at the other targeted sites. **a,** Alignments of Sanger sequencing chromatograms among the PE events carrying homozygous (Ho), heterozygous (He), chimeric (Chi), and imprecise (Imp) forms of the desired edit at the *SlOR* and *SlCAB13* targets. The red arrows denote the peaks containing desired base changes. **b,** Targeted deep sequencing analysis of the PE events obtained at *SlOR*, *SlCENH3*, *SlCAB13*, *SlWH9*, *SlMBP21*, and *SlPRD* sites.

**Extended Data Fig. 5:**
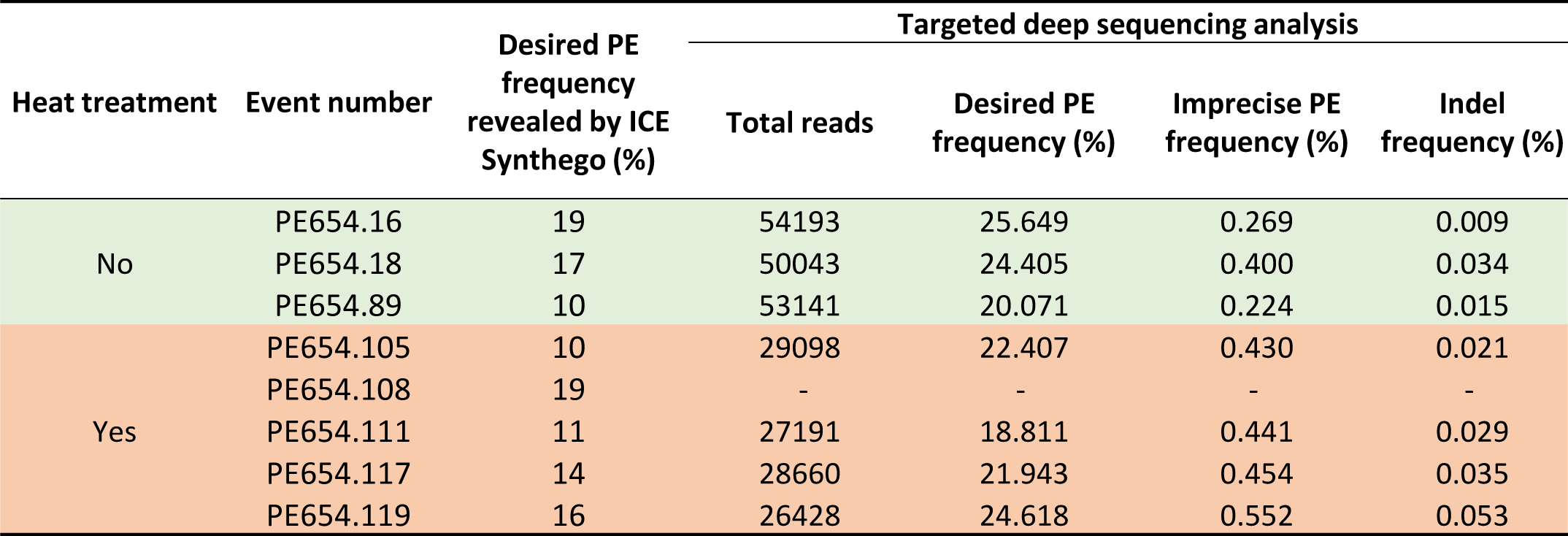
T0 events carrying desired edits obtained at the *AtPDS3* site in *Arabidopsis* without and with heat treatment.

**Extended Data Fig. 6:**
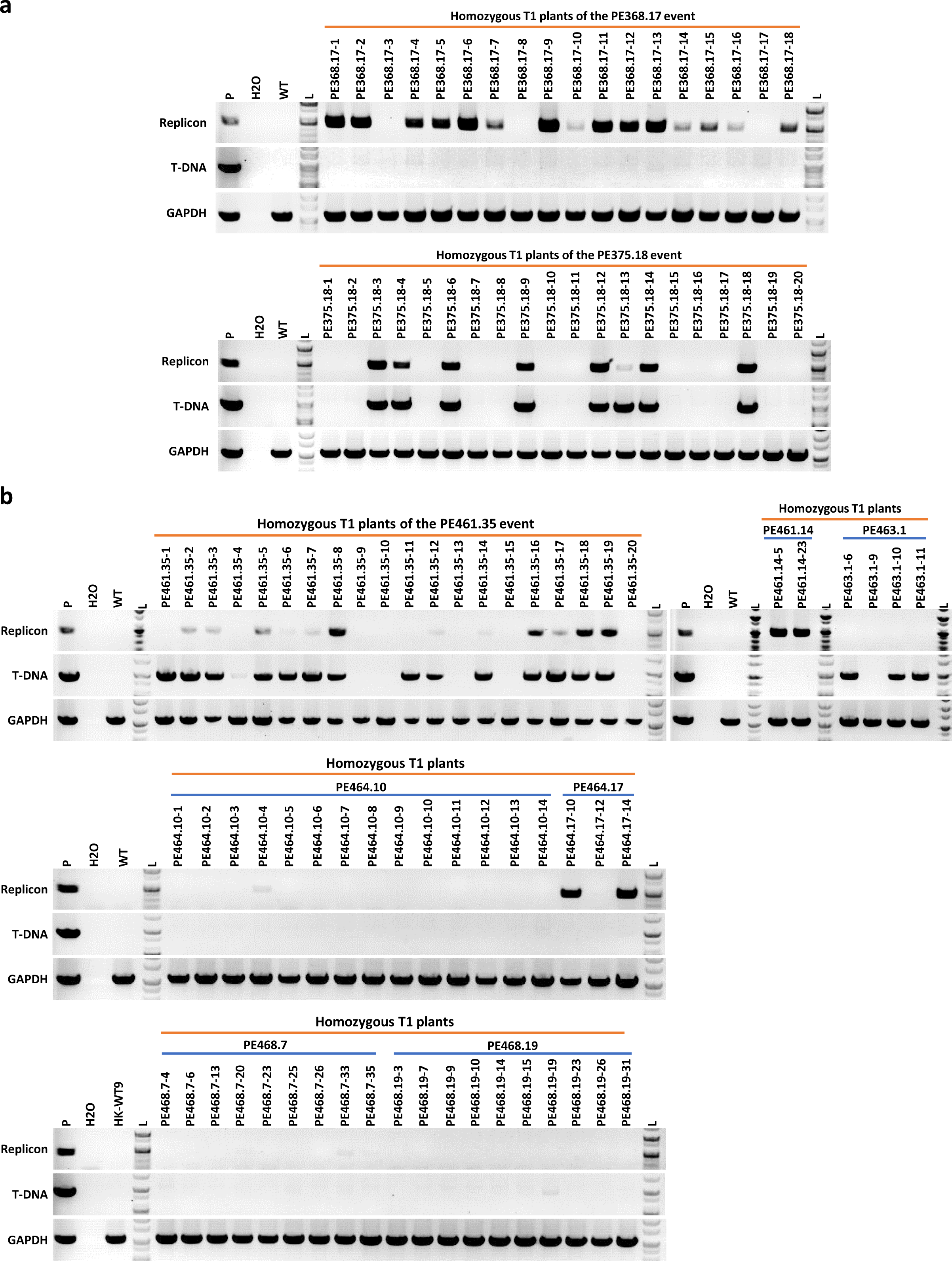
Assessment of the presence of T-DNA and replicon in the T1 events carrying homozygous alleles of the desired edits. **a,** Homozygous T1 offspring of the homozygous T0 events PE368.17 (PE2max) and PE375.18 (ePEmax2). **b,** Homozygous PE1 offspring of the T0 events at *SlOR* (PE461. 14 and PE461.35), *SlCENH3(del)* (PE463.1), *SlCAB13* (PE464.10 and PE464.17), and *SlMBP21* (PE468.7 and PE468.19).

**Extended Data Fig. 7:**
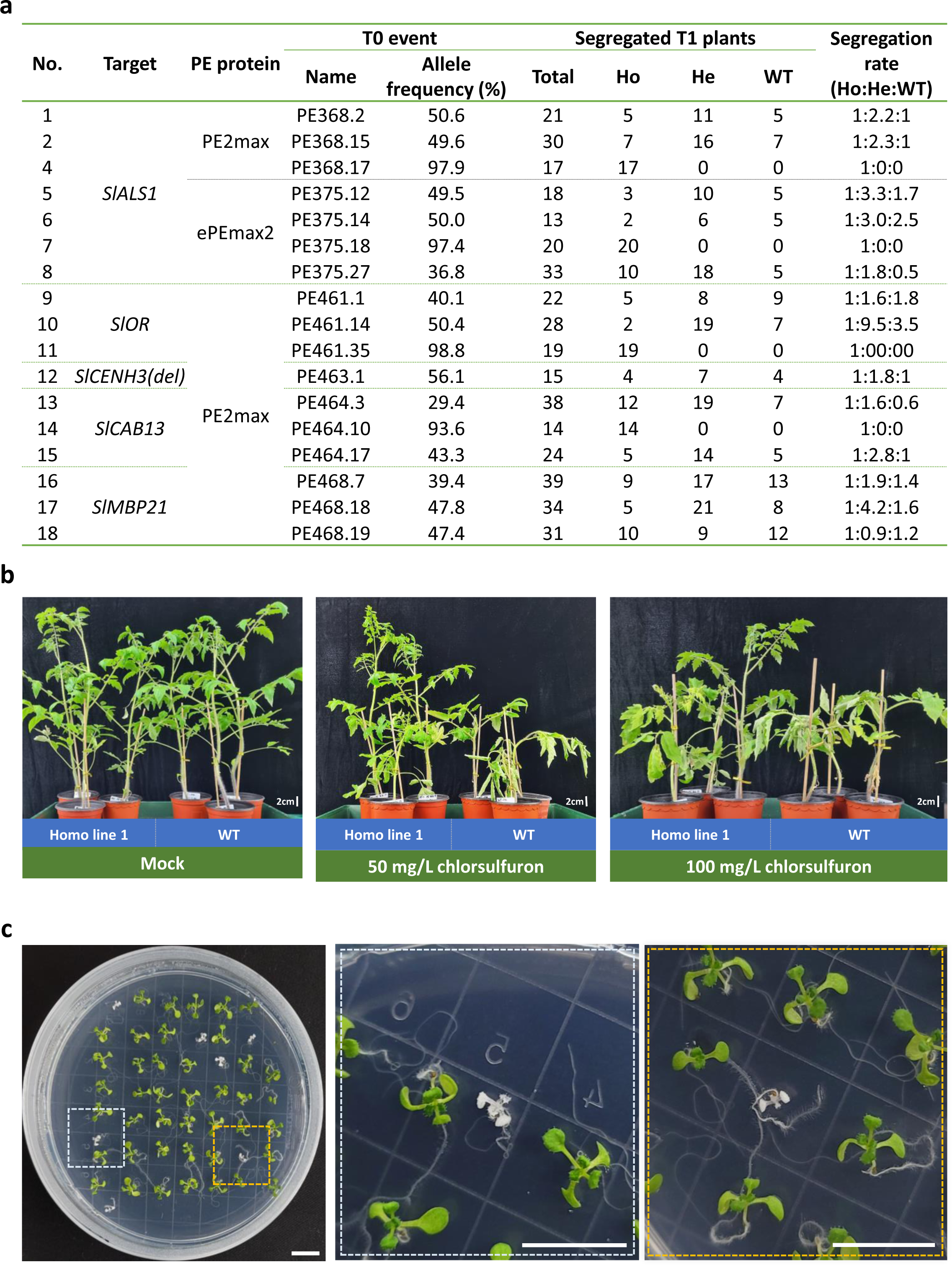
The inheritance of the desired PE edits in the next generation and their phenotypic performance. **a,** Analysis of T1 plants obtained from some representative T0 events of *SlALS1* (from PE2max and ePEmax2), *SlOR*, *SlCENH3(del), SlCAB13*, and Sl*M*BP21. **b,** Chlorsulfuron tolerance of the PE1 plants carrying homozygous alleles (P186S) of *SlALS1*. **c,** T2 seedlings of the AtPDS3 events carrying 19% of the knockout allele (*M30stop*) germinated on an antibiotic-free medium.

## Supplementary information

### Supplementary Tables

**Supplementary Table 1.**
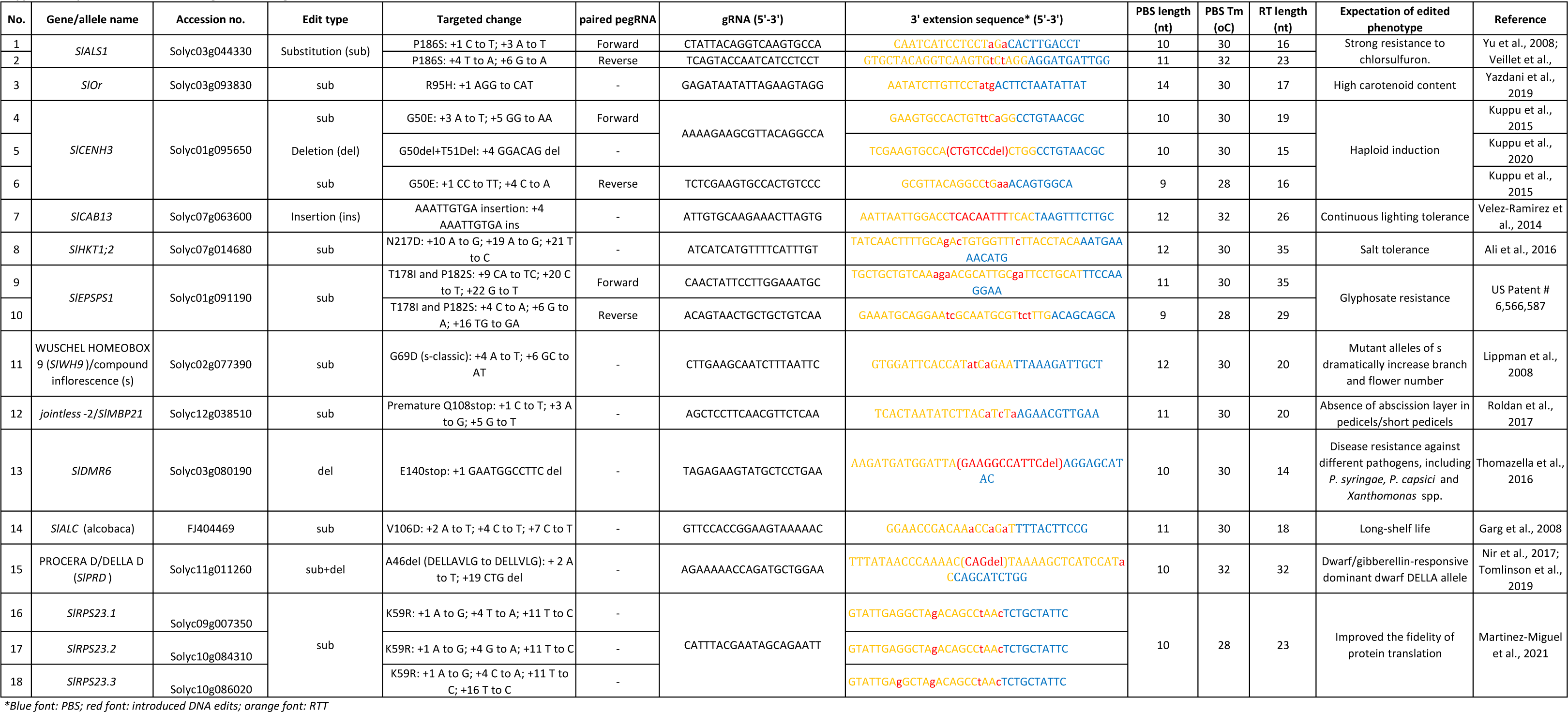
The selected target sites/loci, gRNA, and RTT information for tomatoes.

**Supplementary Table 2.**
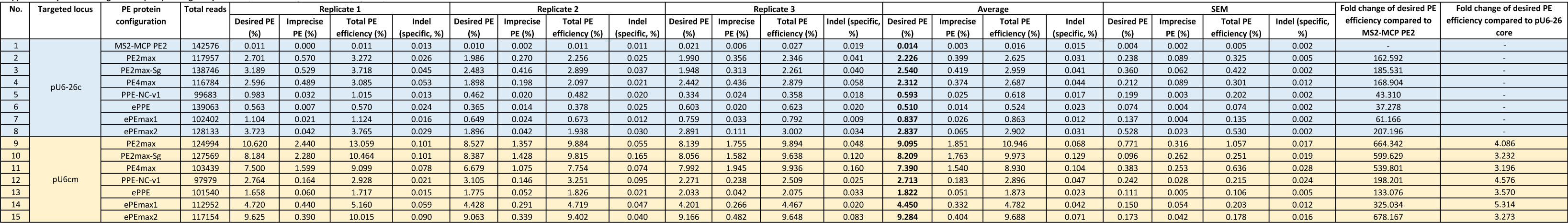
Targeted deep sequencing analysis of PE protein configurations and pegRNA promoters.

**Supplementary Table 3.**
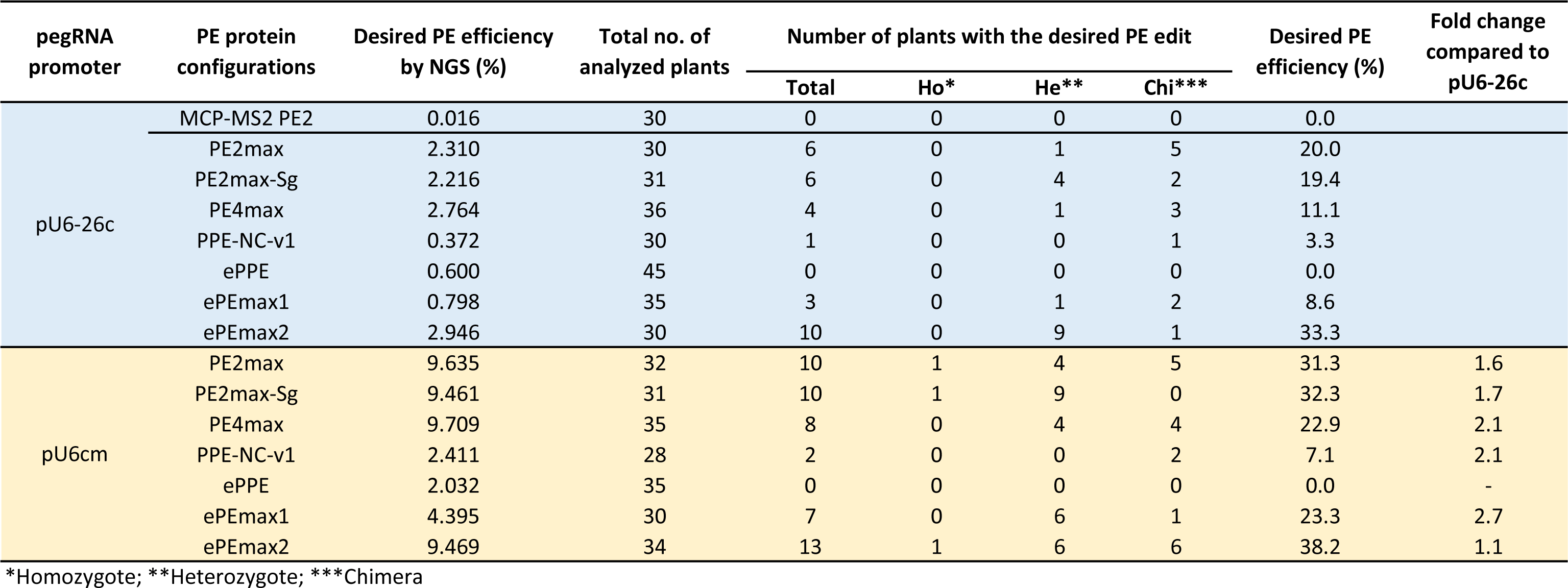
Summarized data of transformant screening for the PE protein configuration and pegRNA promoter combinations.

**Supplementary Table 4.**
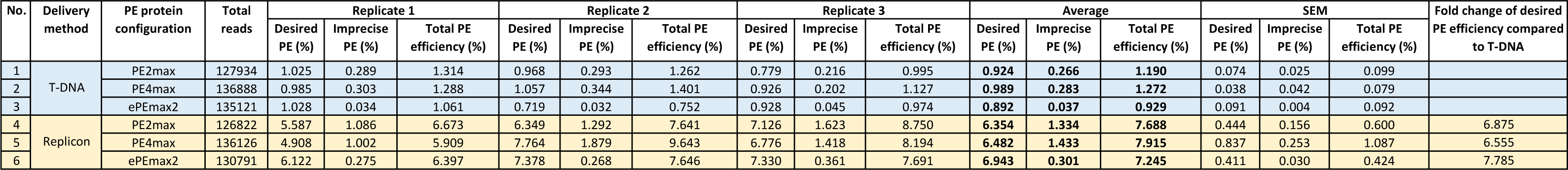
Targeted deep sequencing analysis for the PE tool delivery methods.

**Supplementary Table 5.**
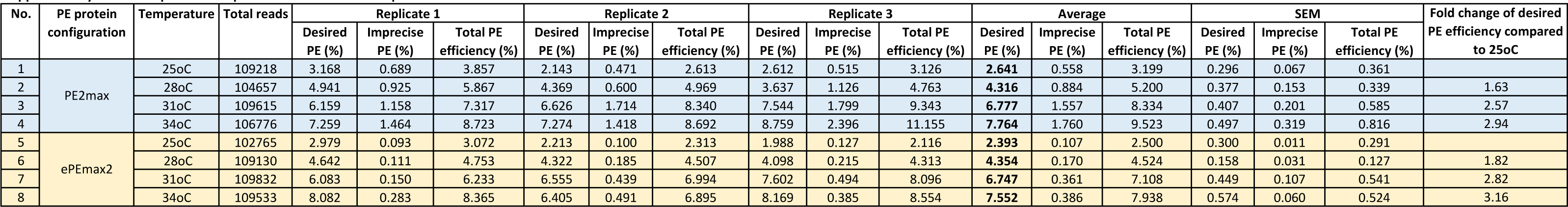
Impacts of temperature treatments on PE performance assessed by targeted deep sequencing at the callus stage.

**Supplementary Table 6.**
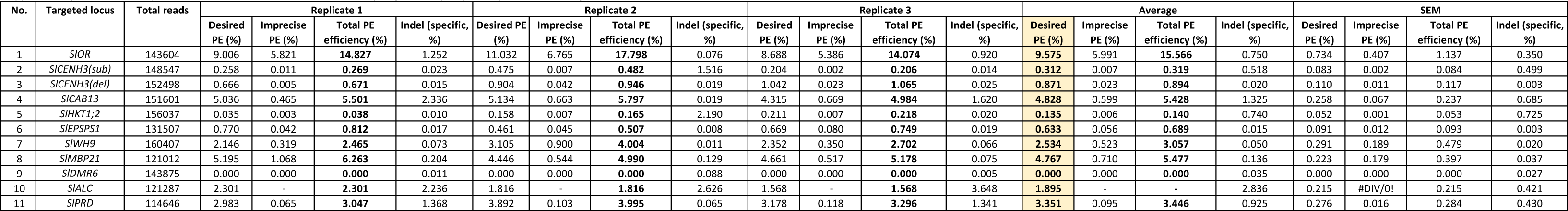
The PE performance on various tomato loci assessed by targeted deep sequencing at the callus stage.

**Supplementary Table 7.**
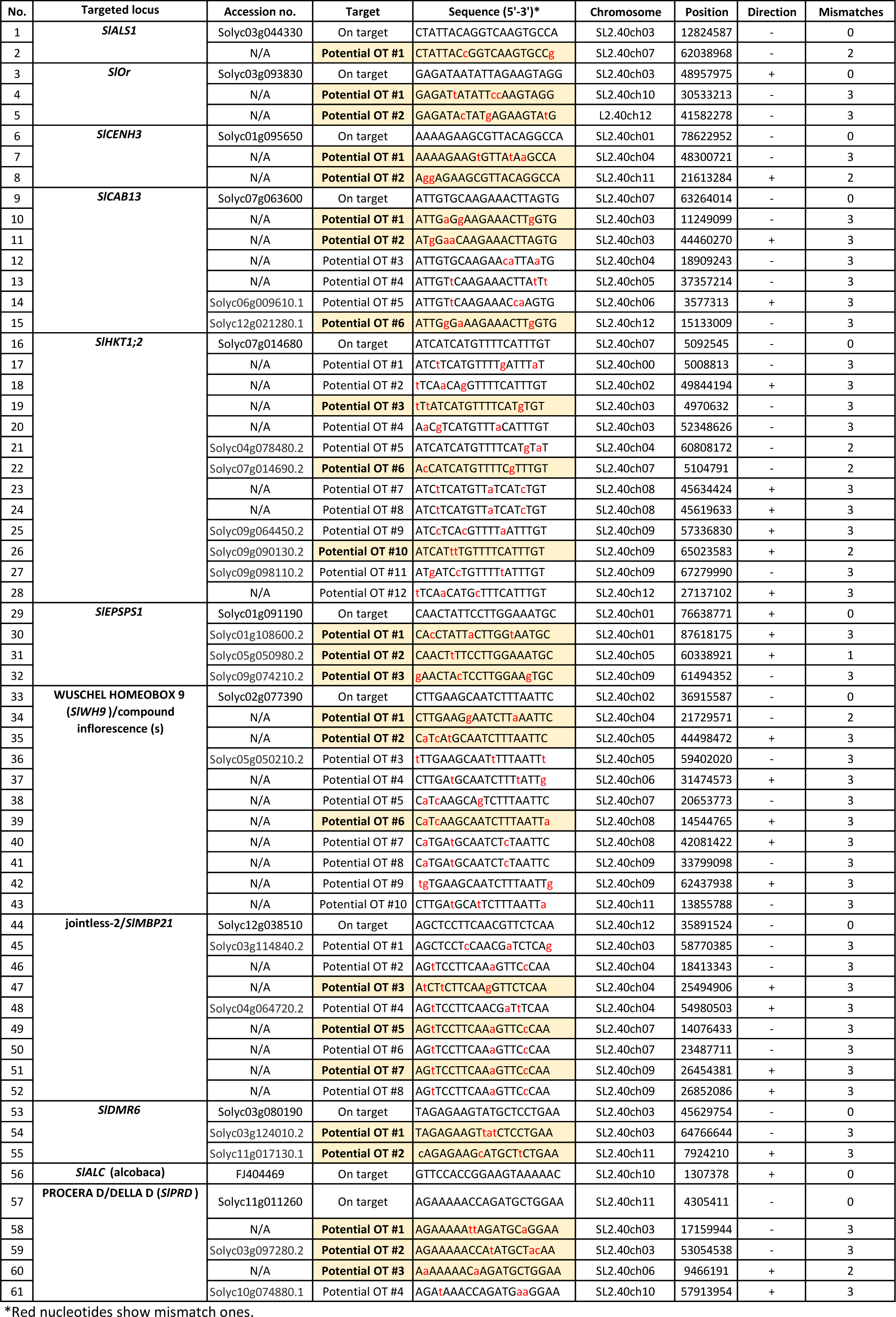
Information of potential off-target sites of the gRNAs used in the study.

**Supplementary Table 8.**
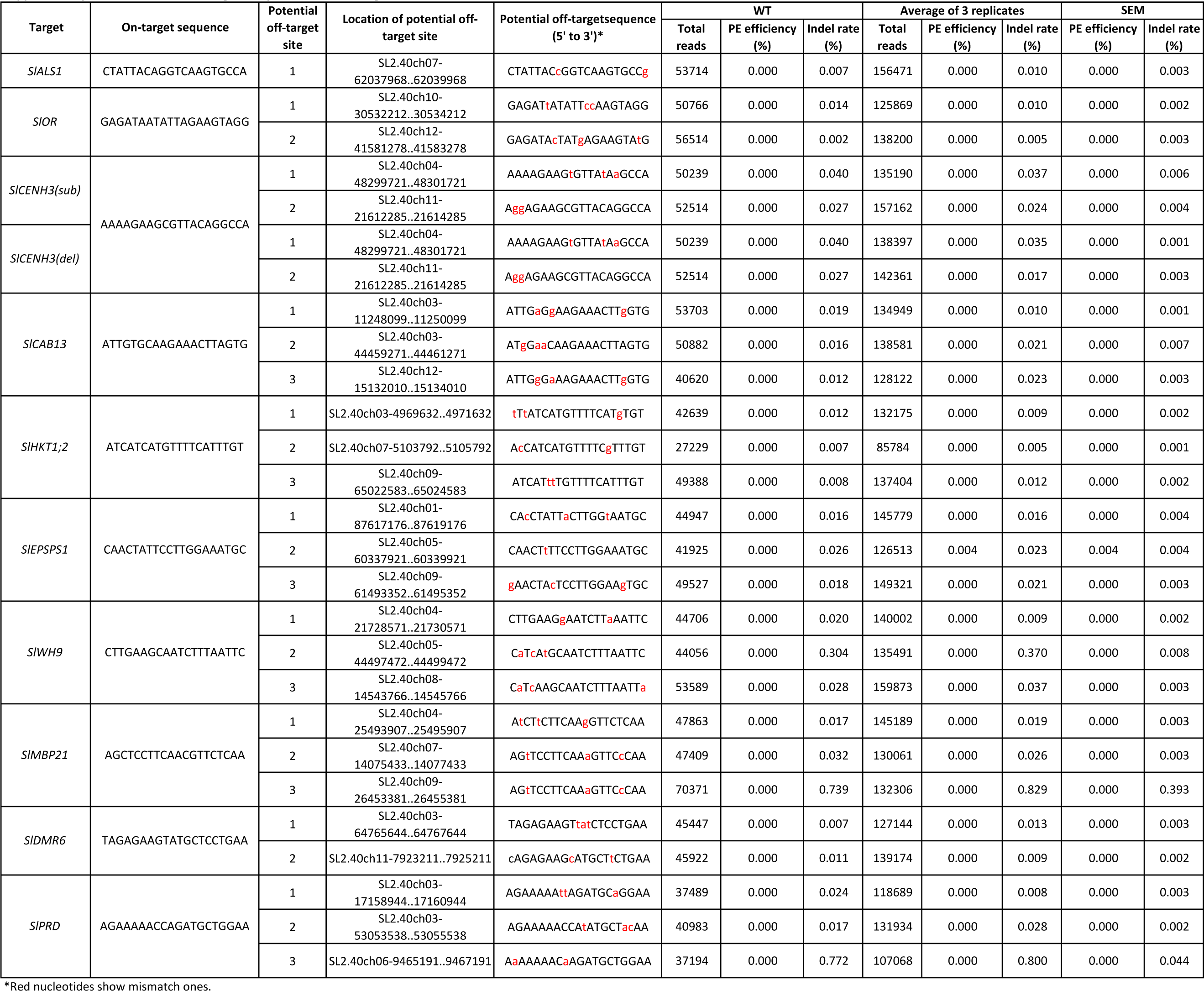
Potential off-target analysis at the callus stage.

**Supplementary Table 9.**
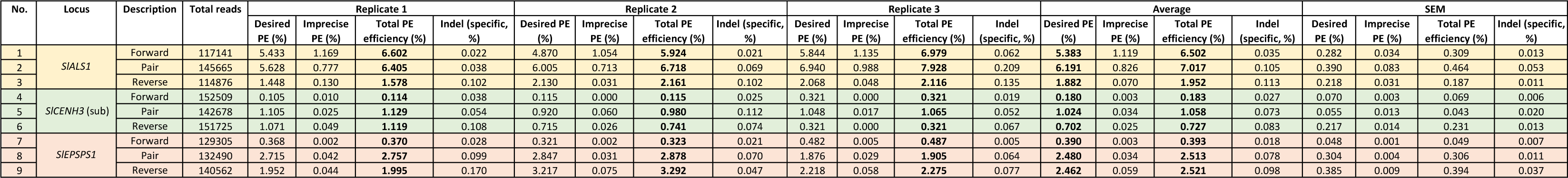
Improvement of PE performance using paired pegRNAs.

**Supplementary Table 10.**
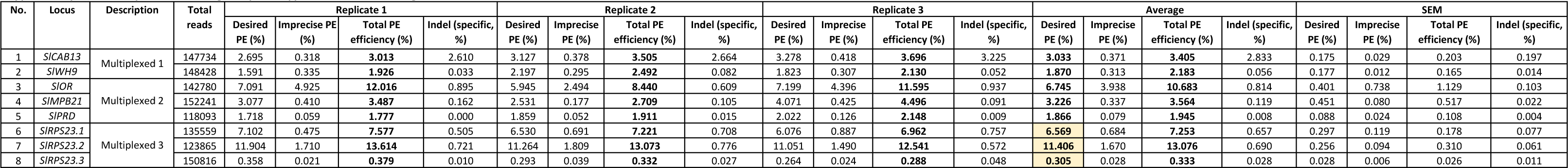
PE performance using multiplexed approaches at the callus stage.

**Supplementary Table 11.**
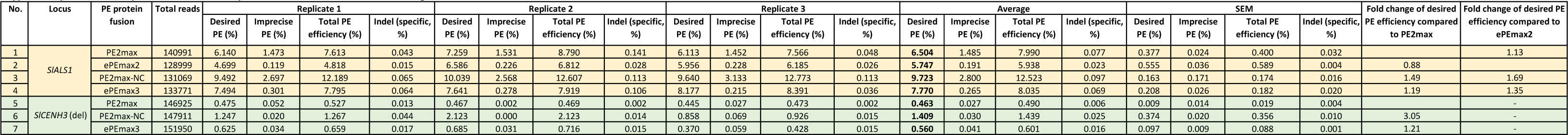
The impacts of NC on PE efficiency in PE2max-NC and ePEmax3 at the callus stage.

**Supplementary Table 12.**
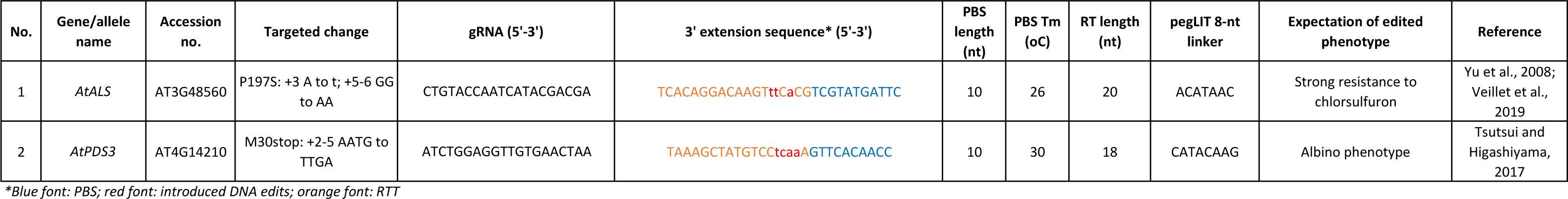
The selected target sites/loci, gRNA, and RTT information for *Arabidopsis*.

**Supplementary Table 13.**
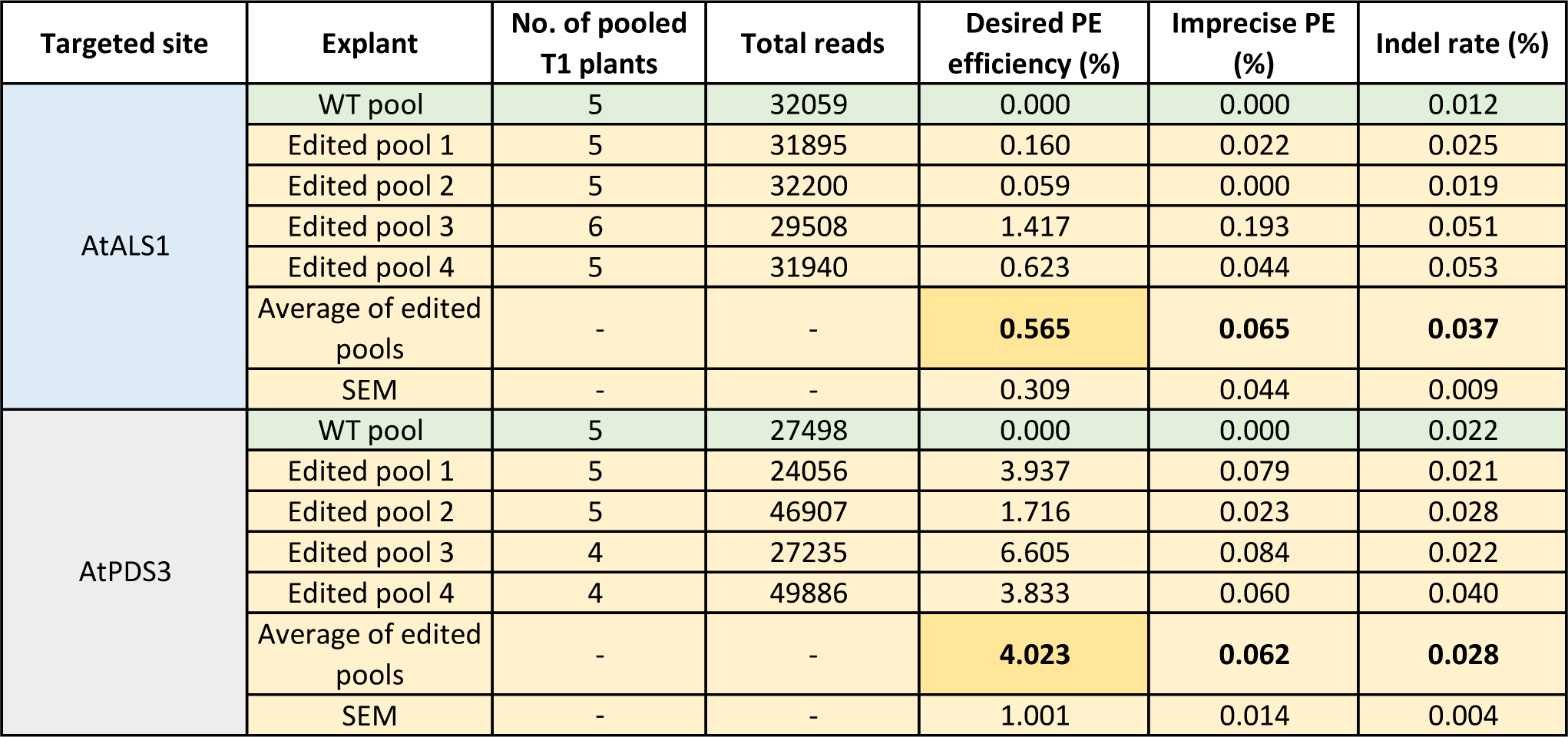
Targeted deep sequencing analysis of pooled T1 plants.

**Supplementary Table 14.**
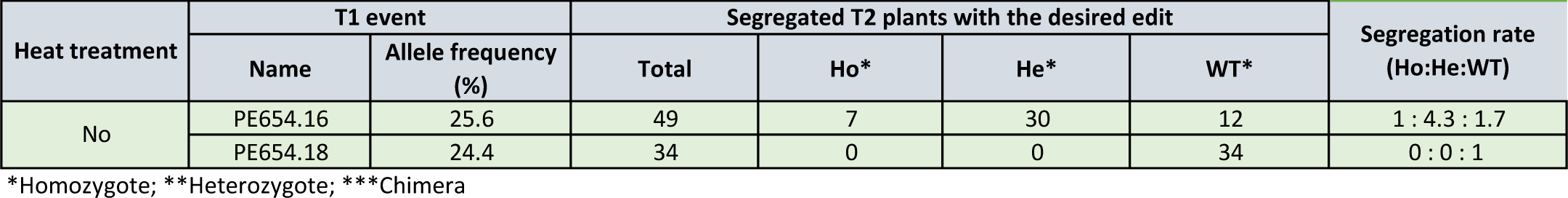
The inheritance of the knockout allele of AtPDS3 in T2 generation.

**Supplementary Table 15.**
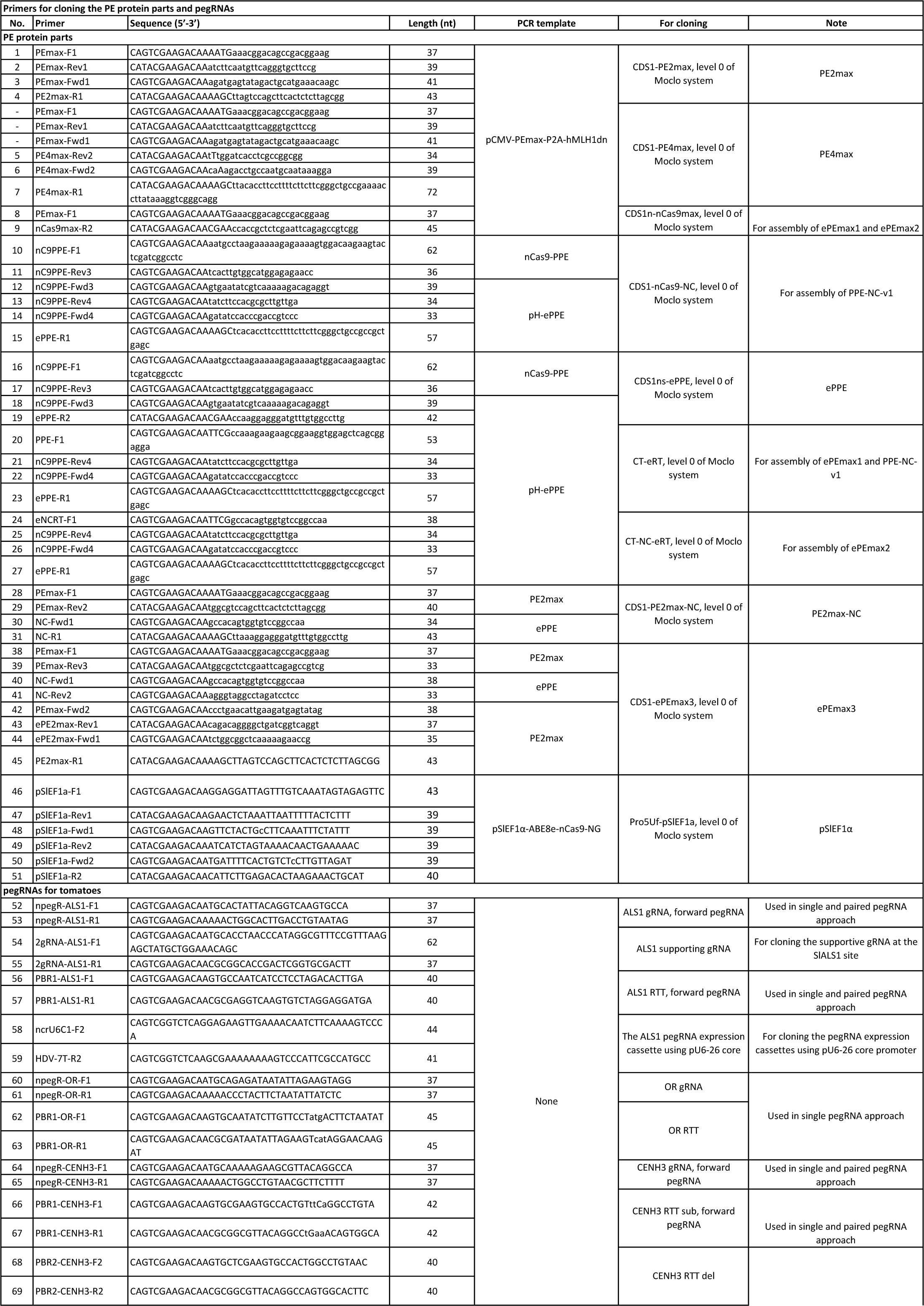
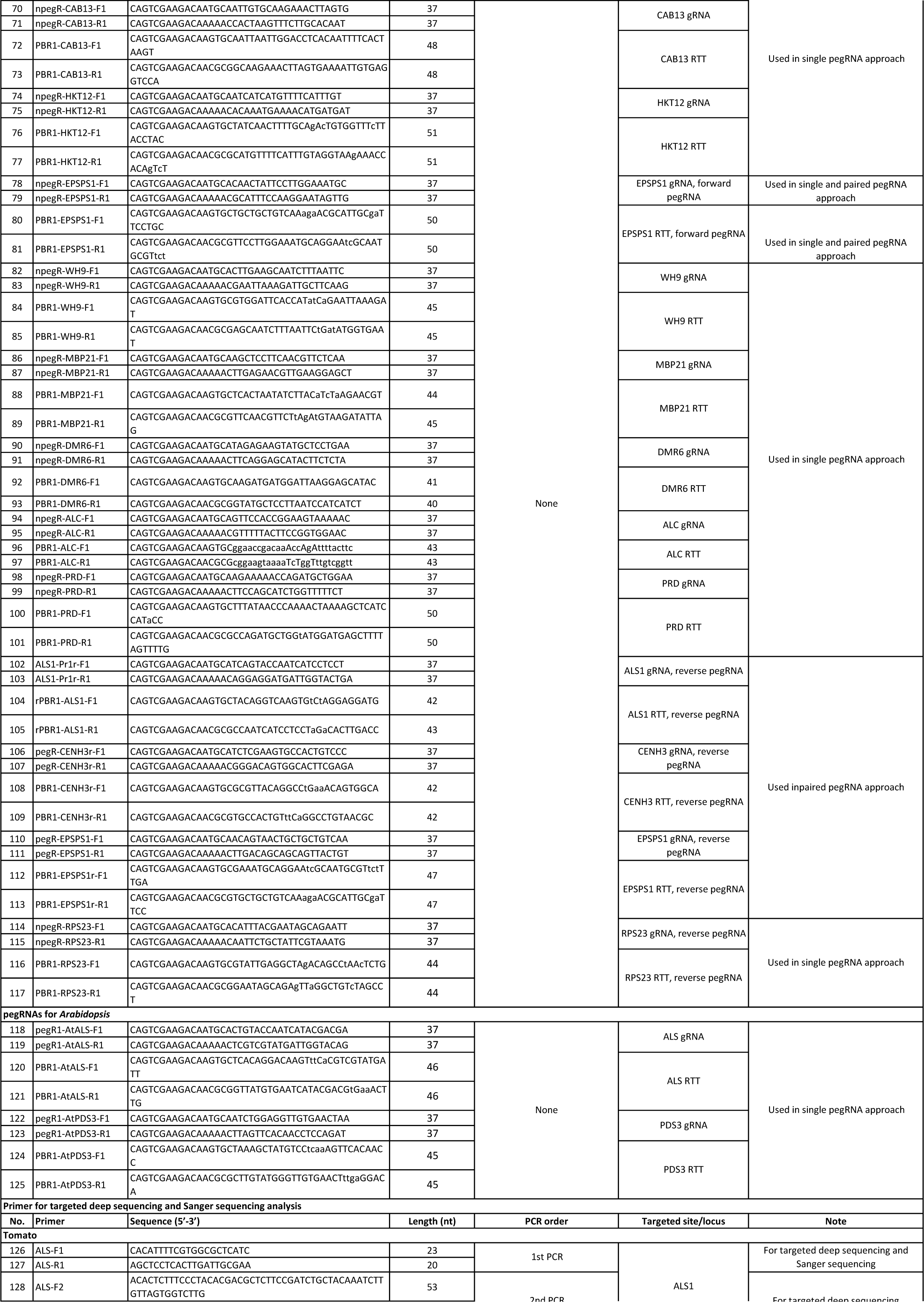
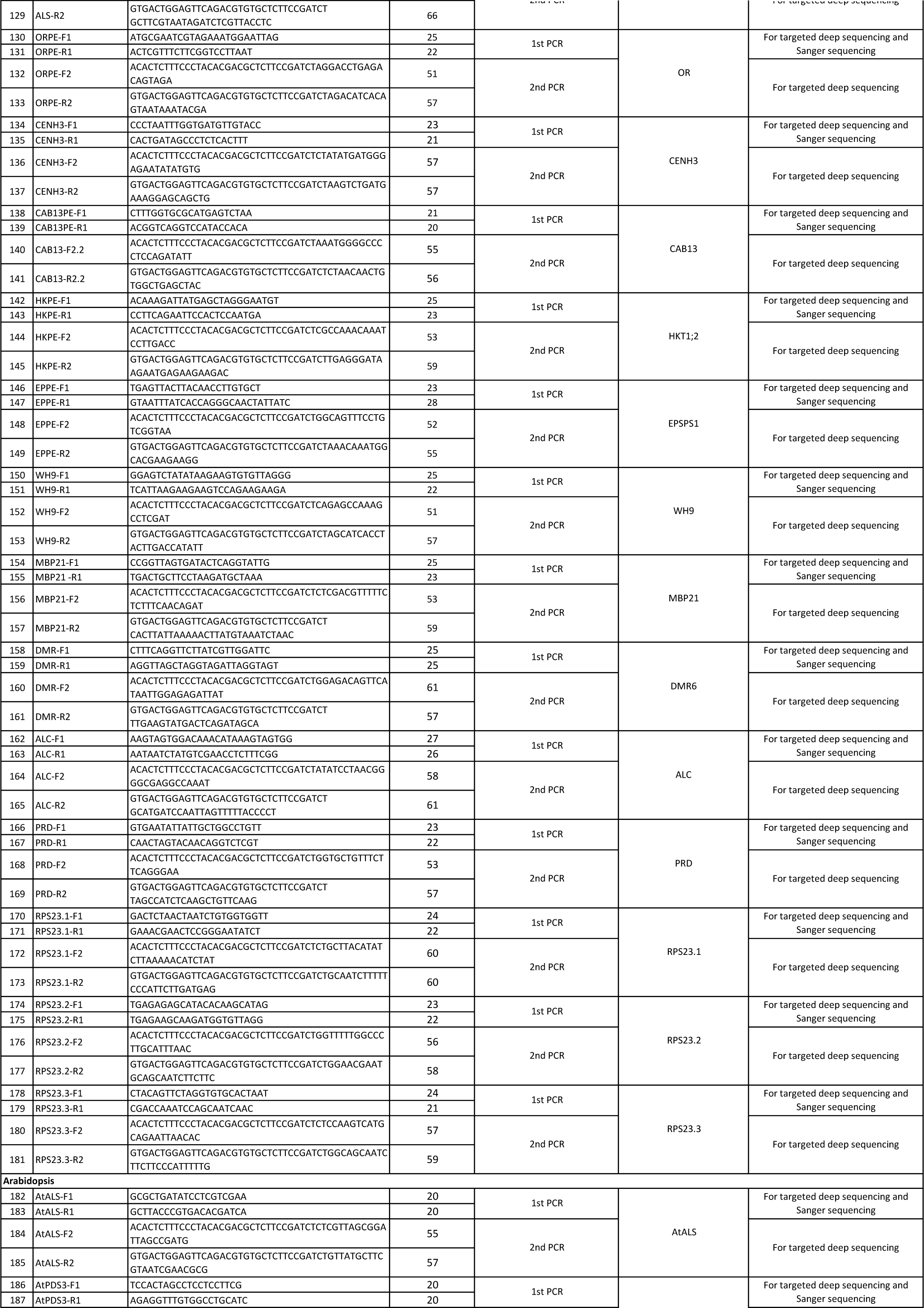
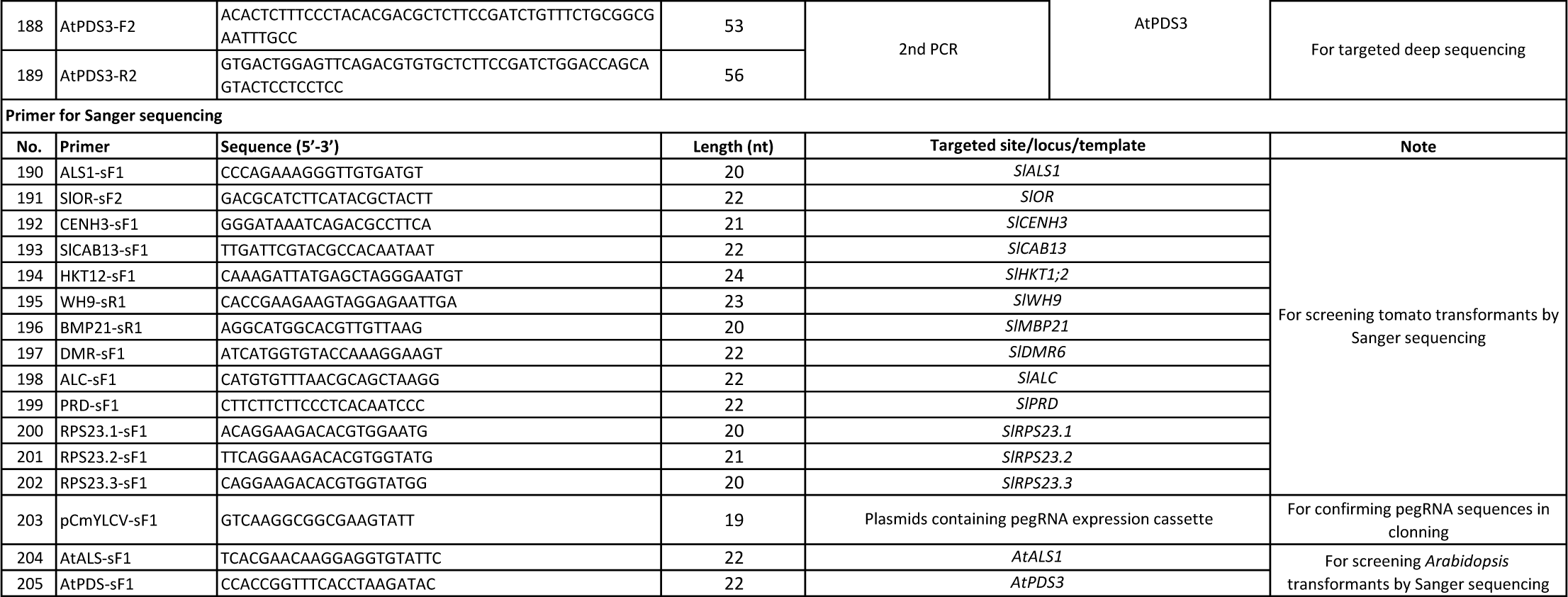
Primers used in the study.

**Supplementary Table 16.**
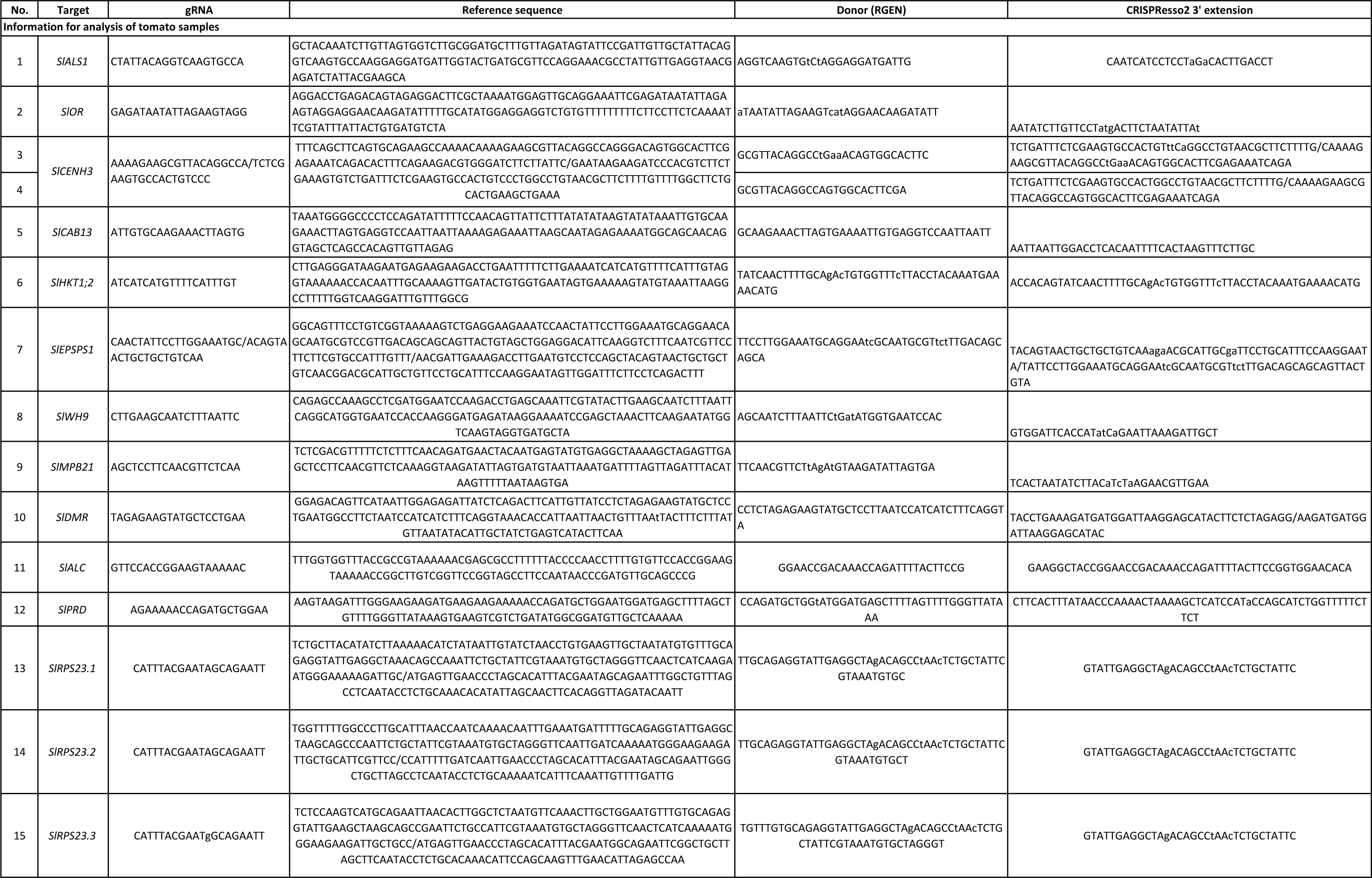
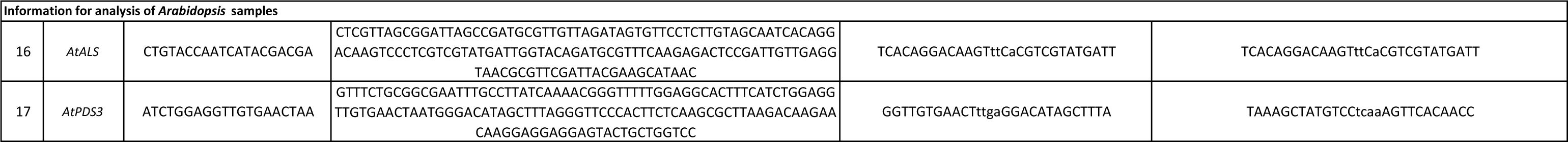
Sequence information used for RGEN and CRISPResso2 analysis of targeted deep sequencing data.

### Supplementary Files

Supplementary file 1: Supplementary Sequence

## SEQUENCES USED IN THE STUDY

### PE2max, cloned from plasmid pCMV-PEmax-P2A-hMLH1dn (Addgene plasmid #174828)

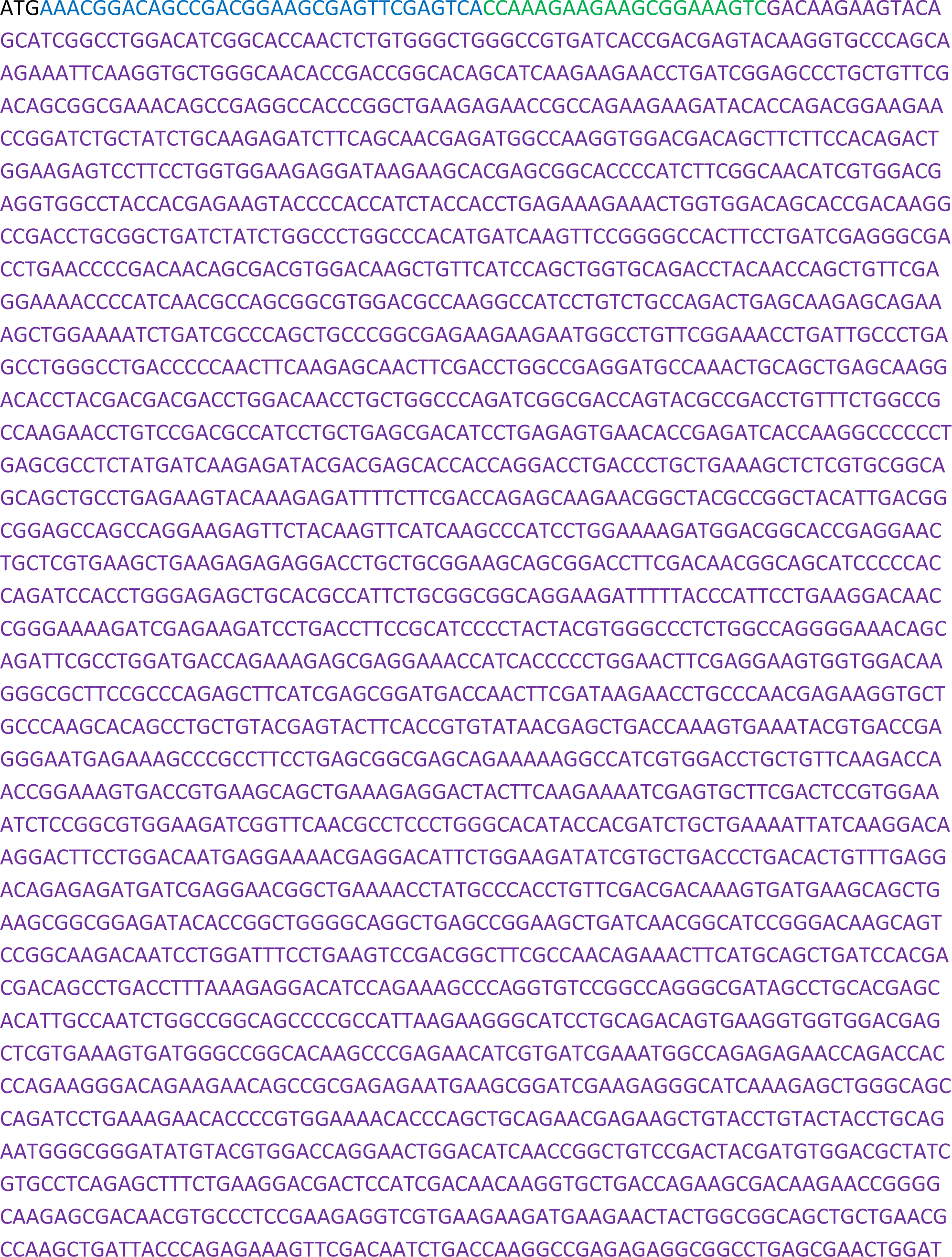
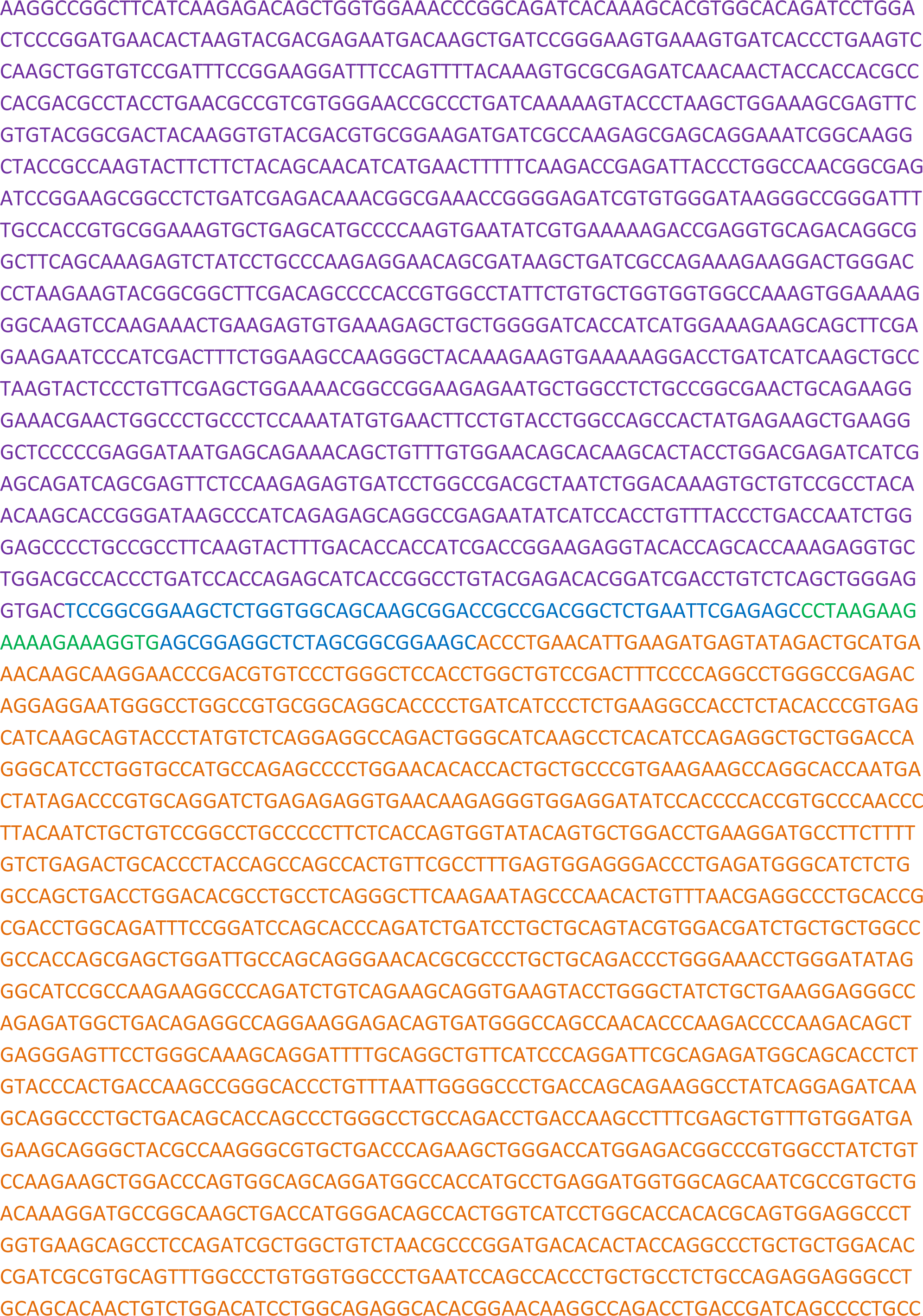
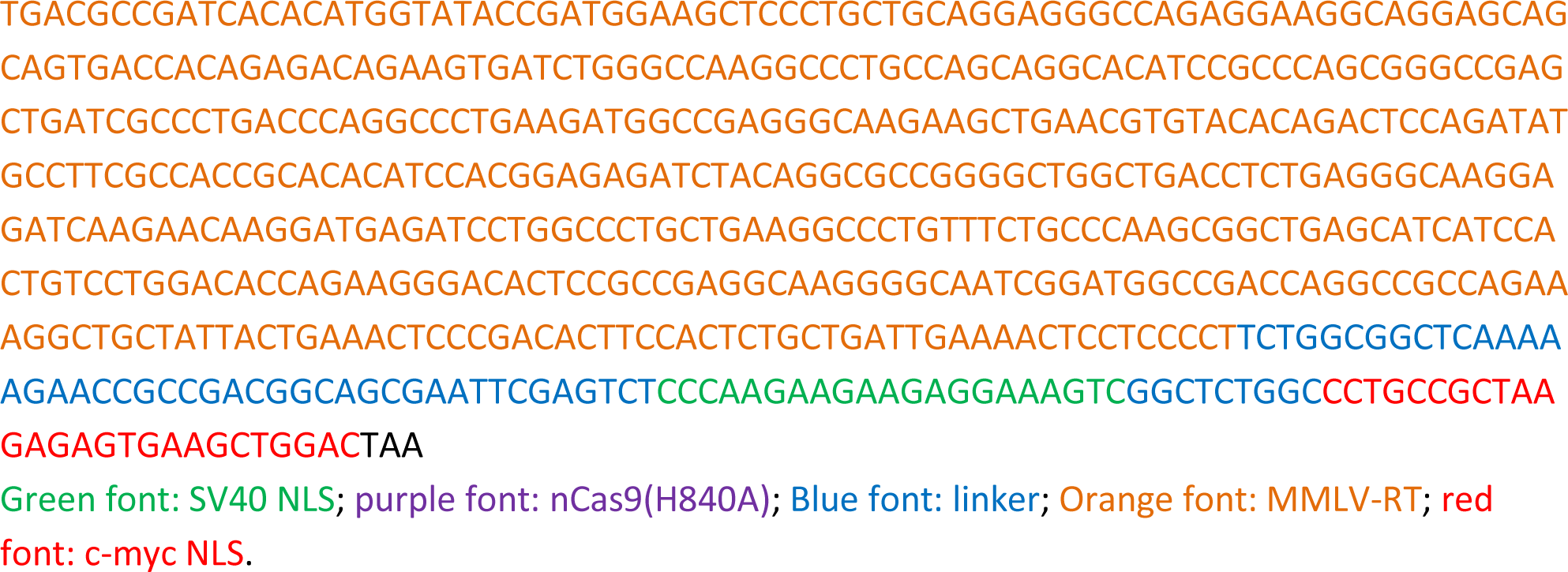

### PE4max, cloned from plasmid pCMV-PEmax-P2A-hMLH1dn (Addgene plasmid #174828)

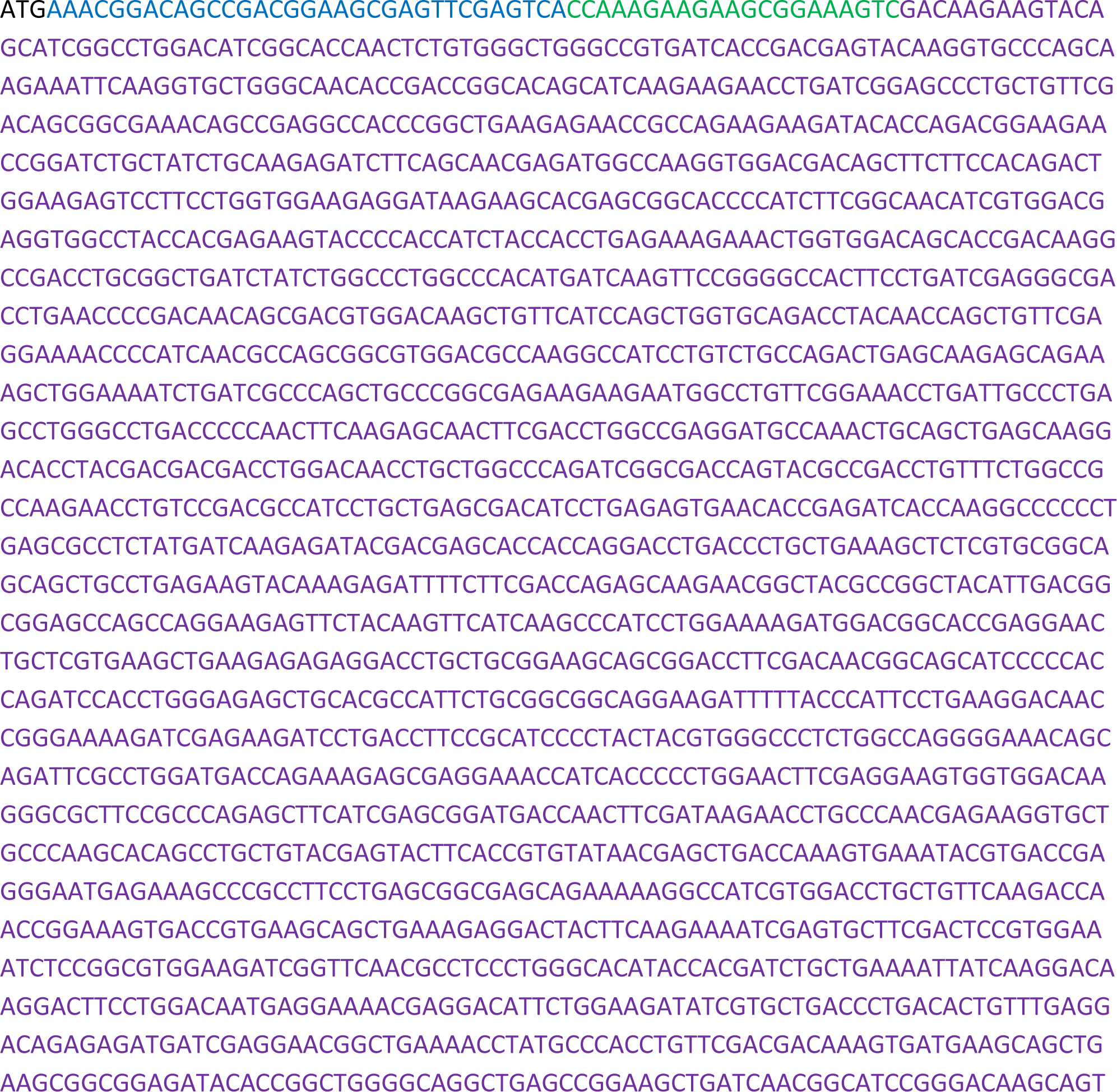
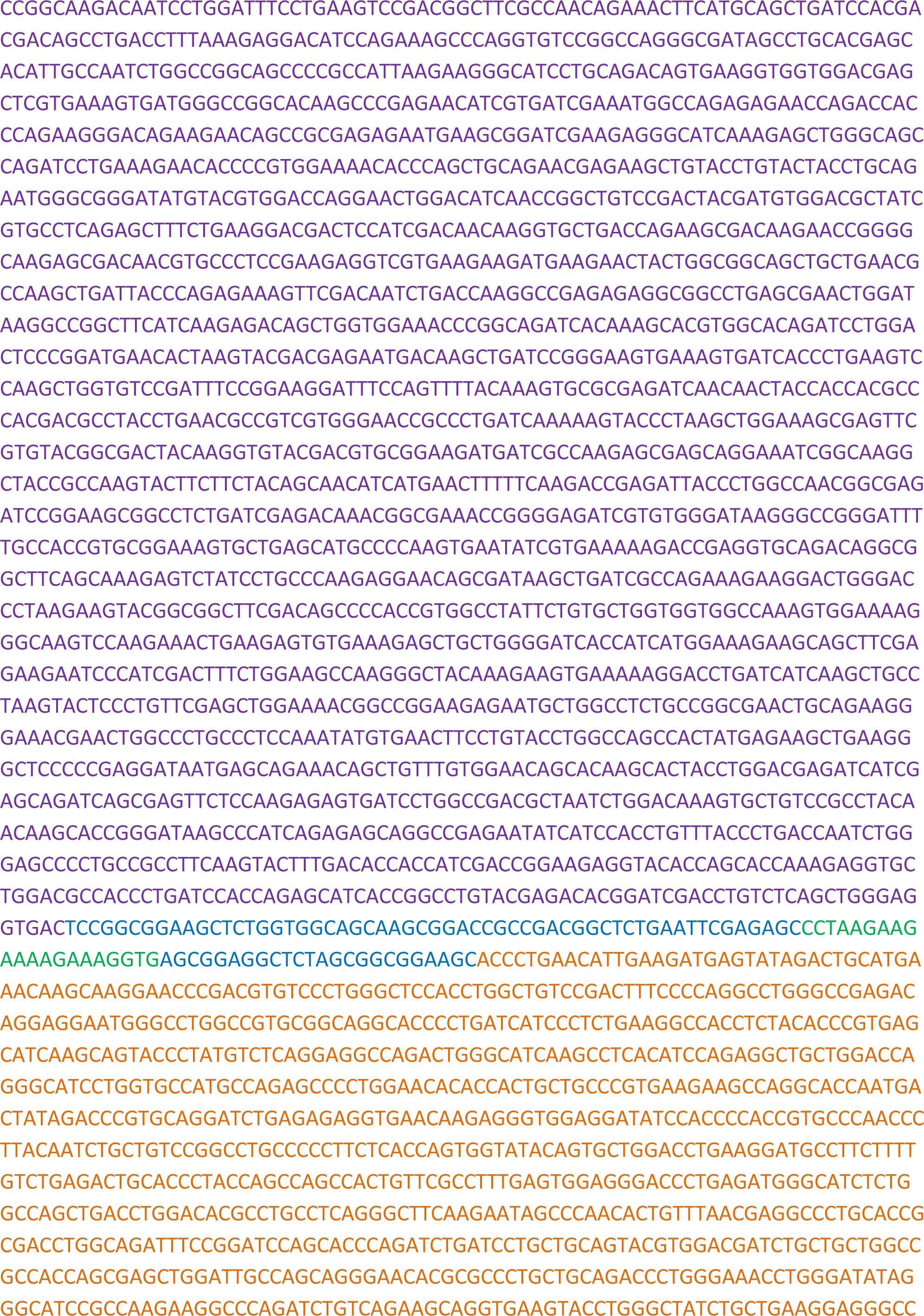
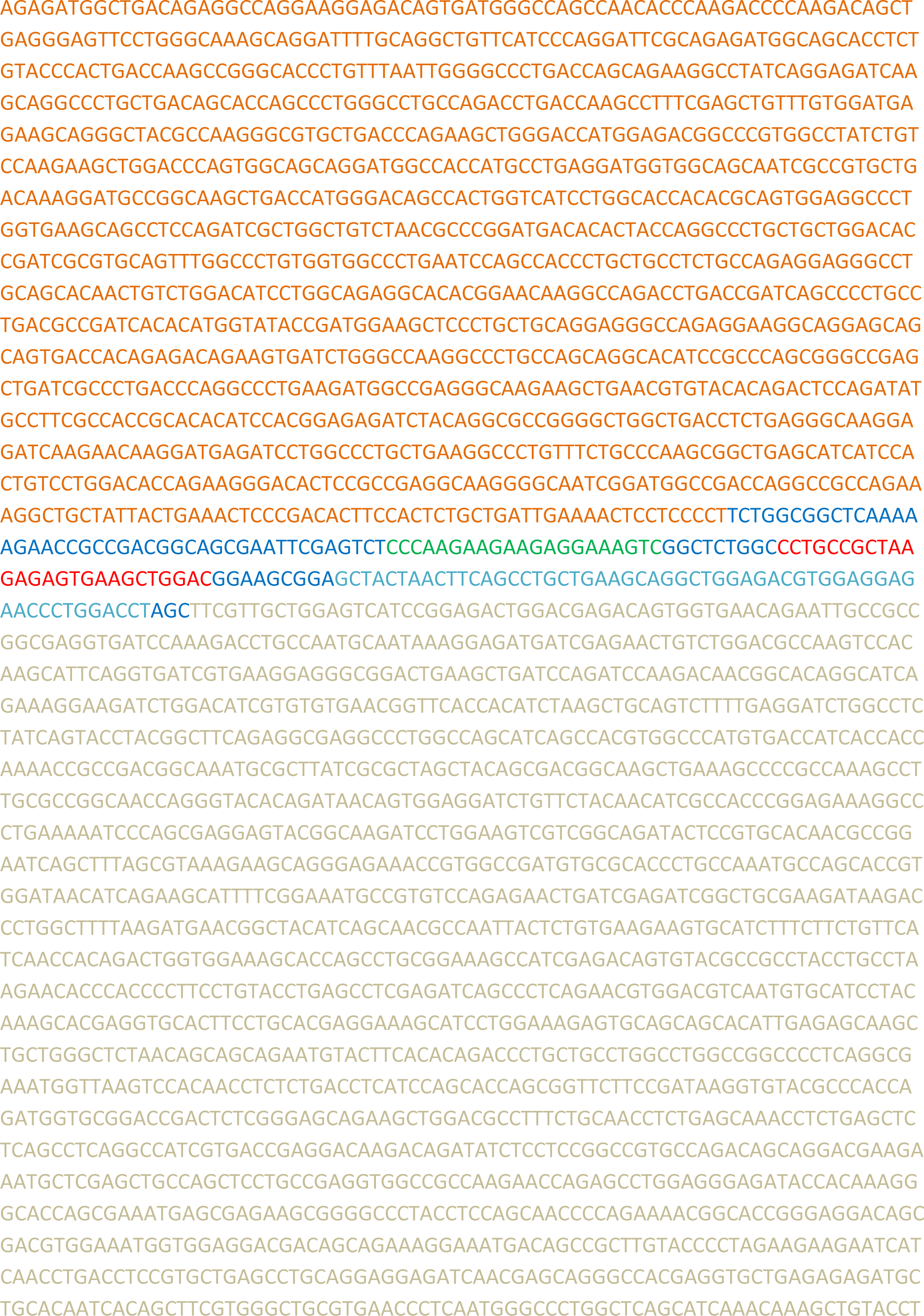
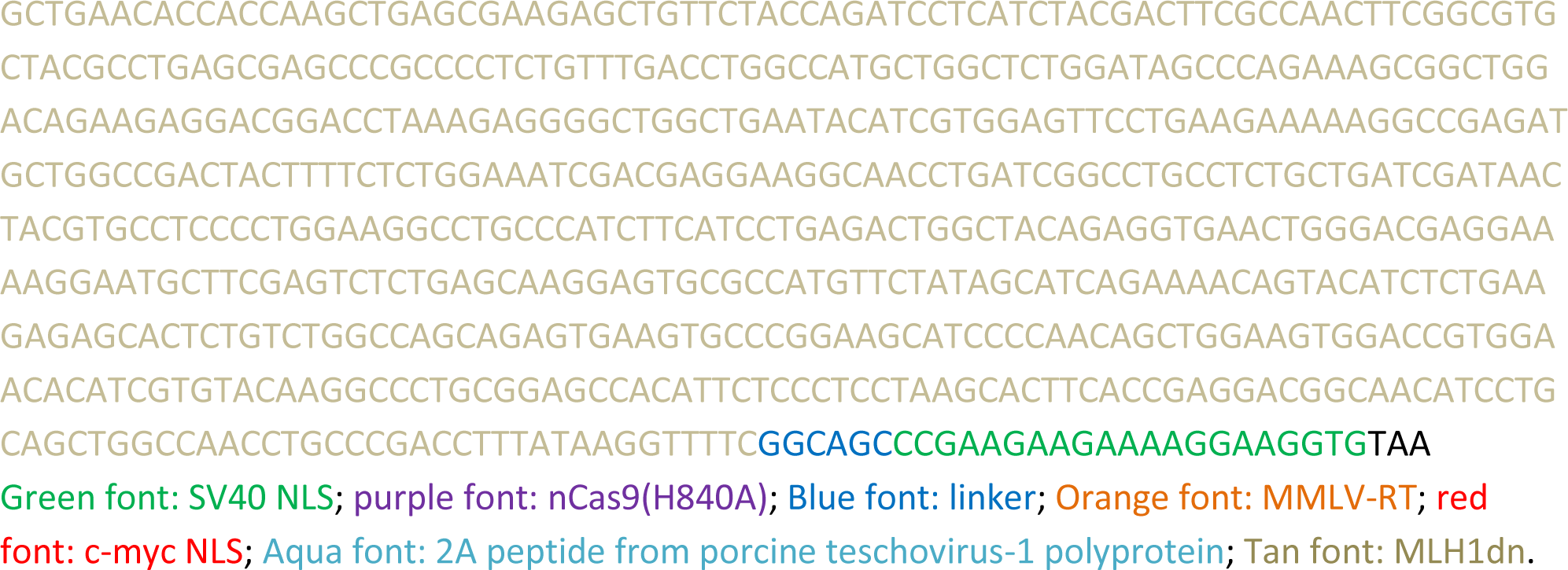

### PPE-NC-v1

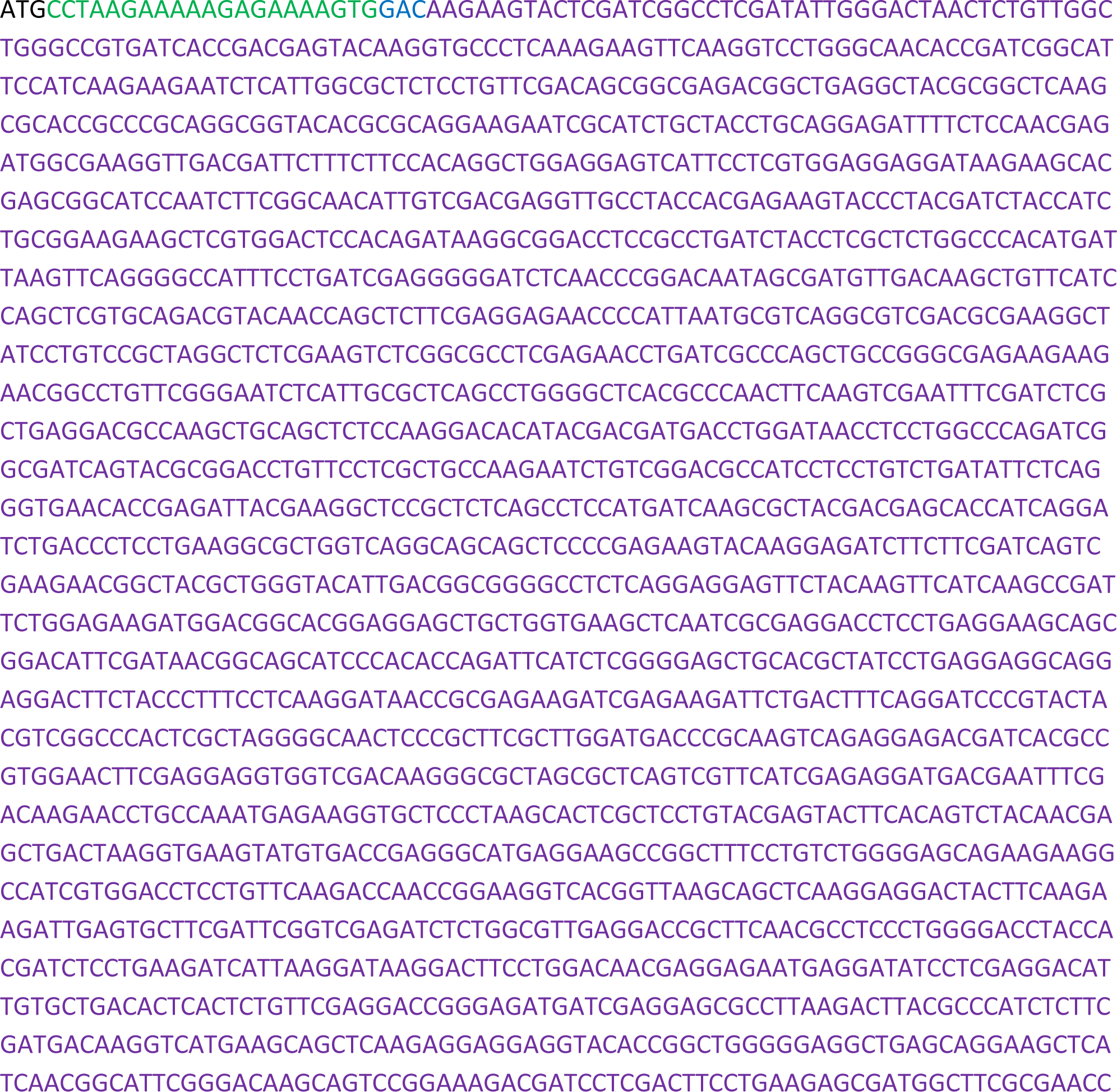
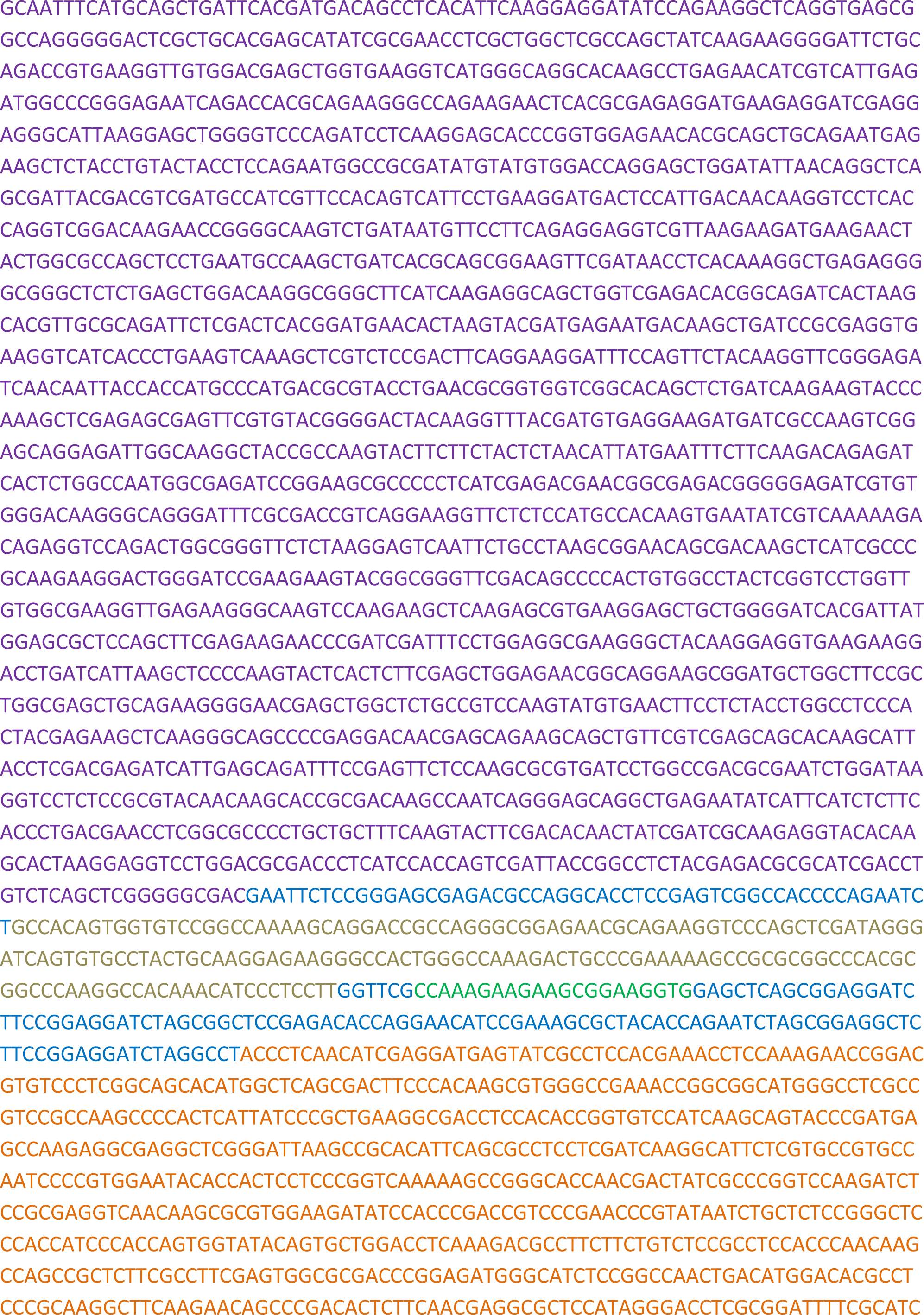
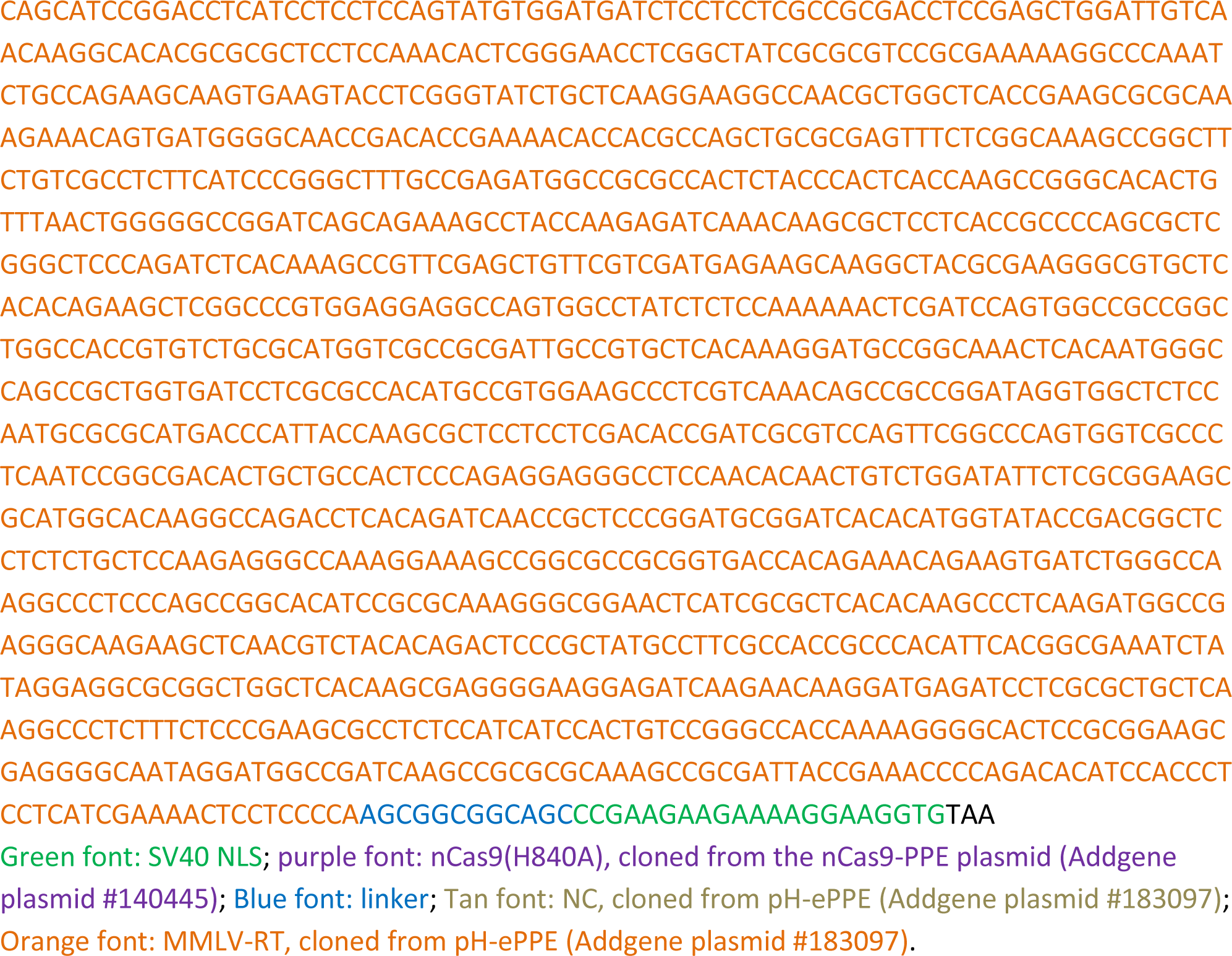

### ePPE, cloned from pH-ePPE (Addgene plasmid #183097)

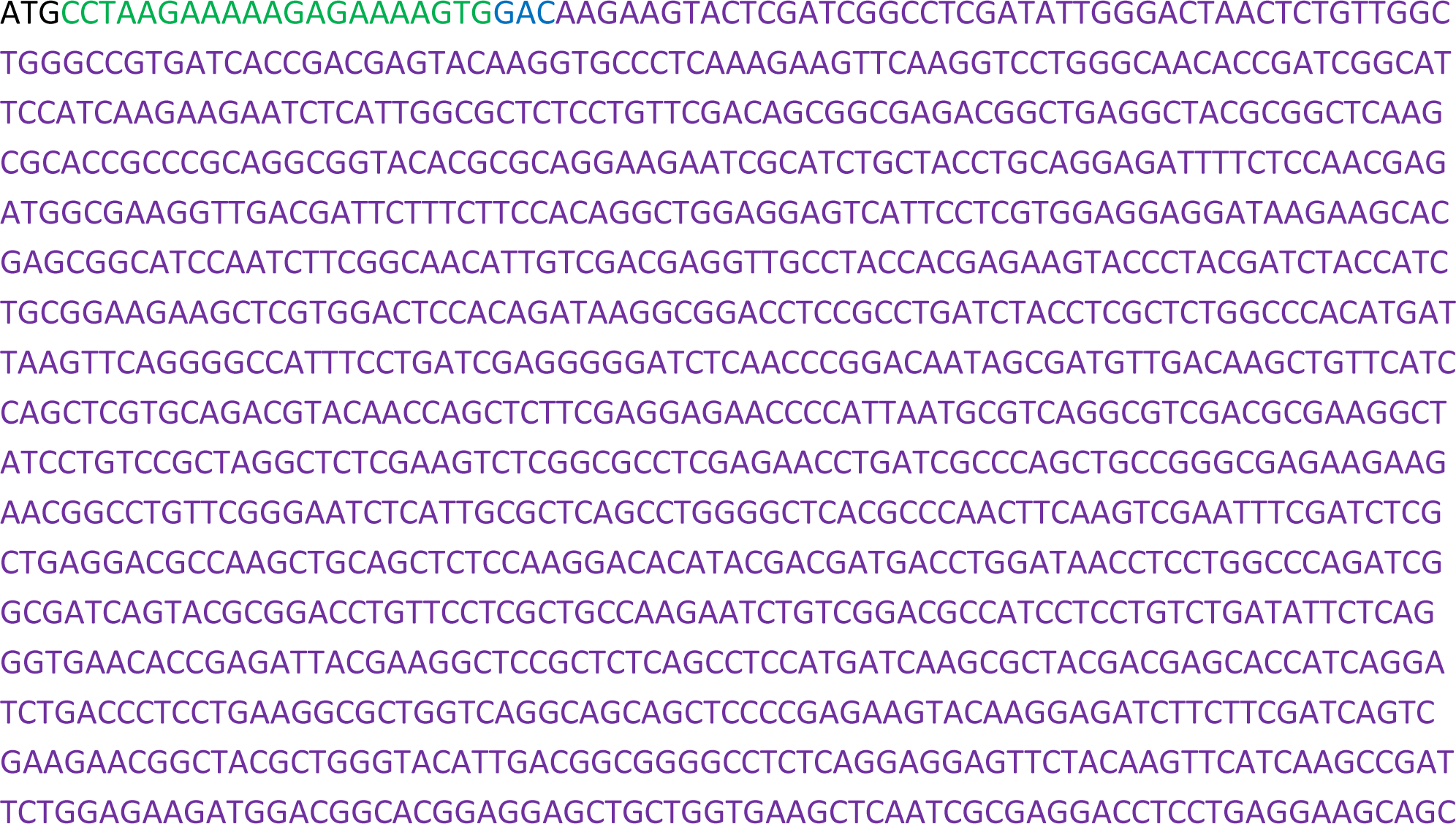
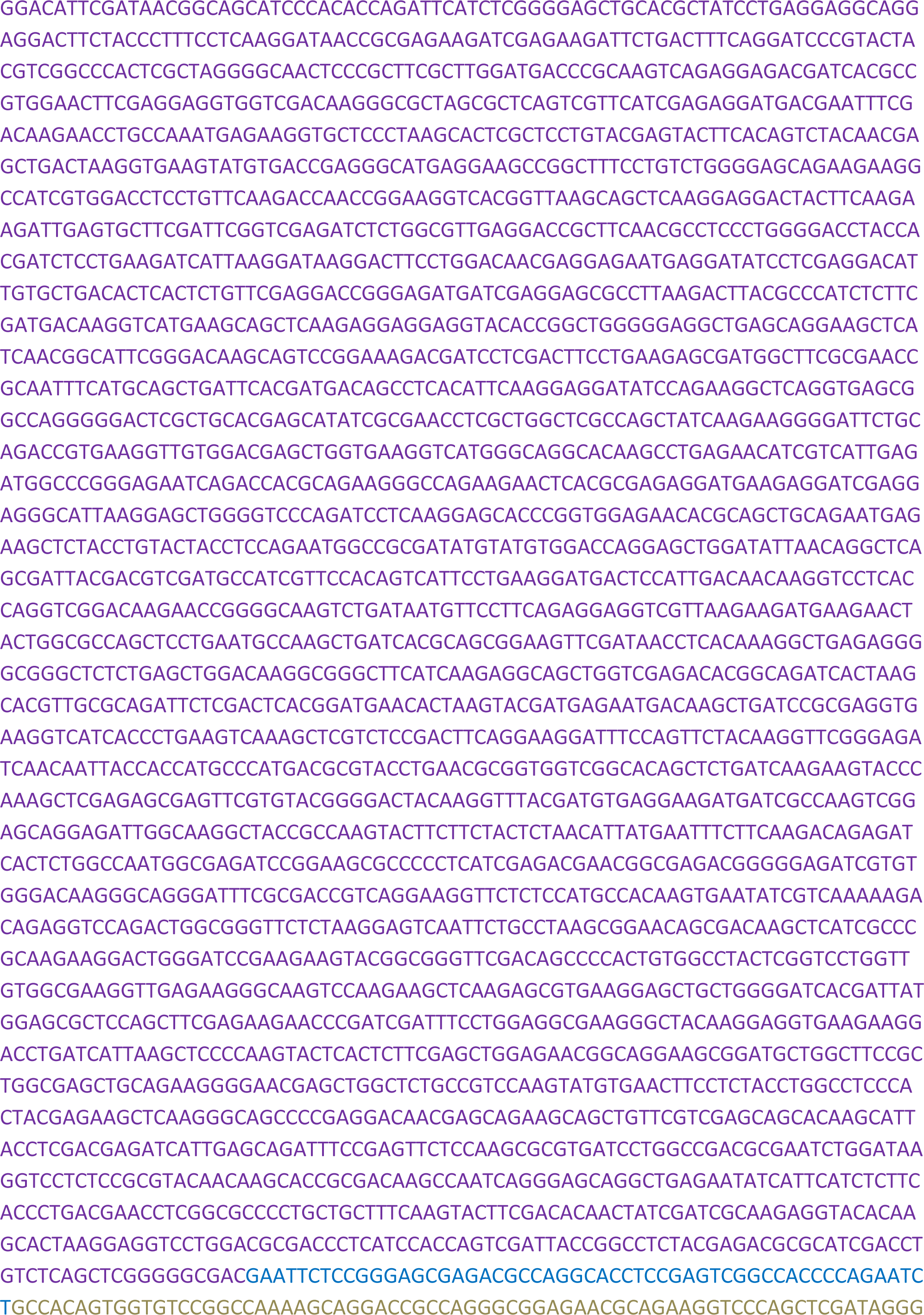
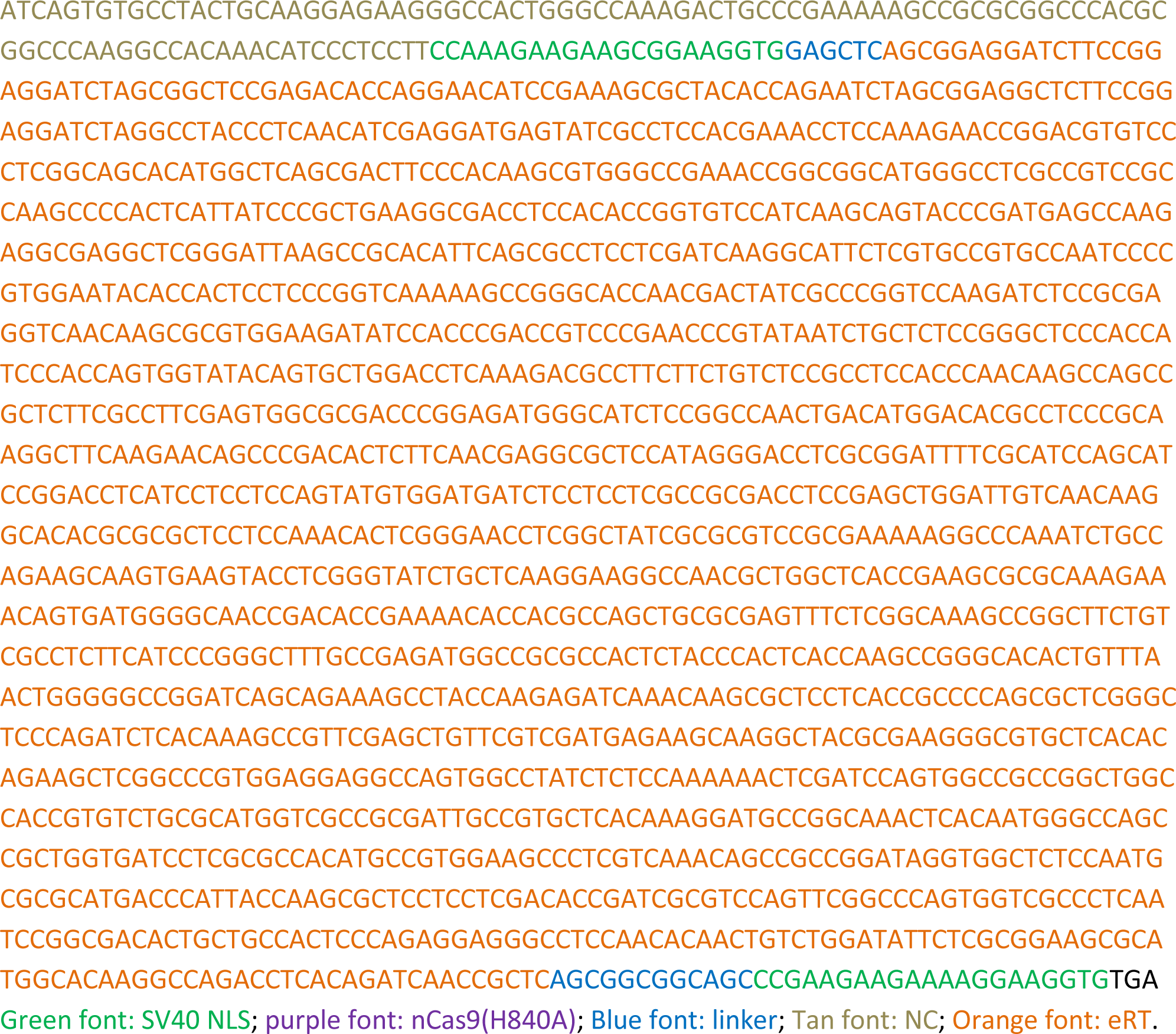

### ePEmax1

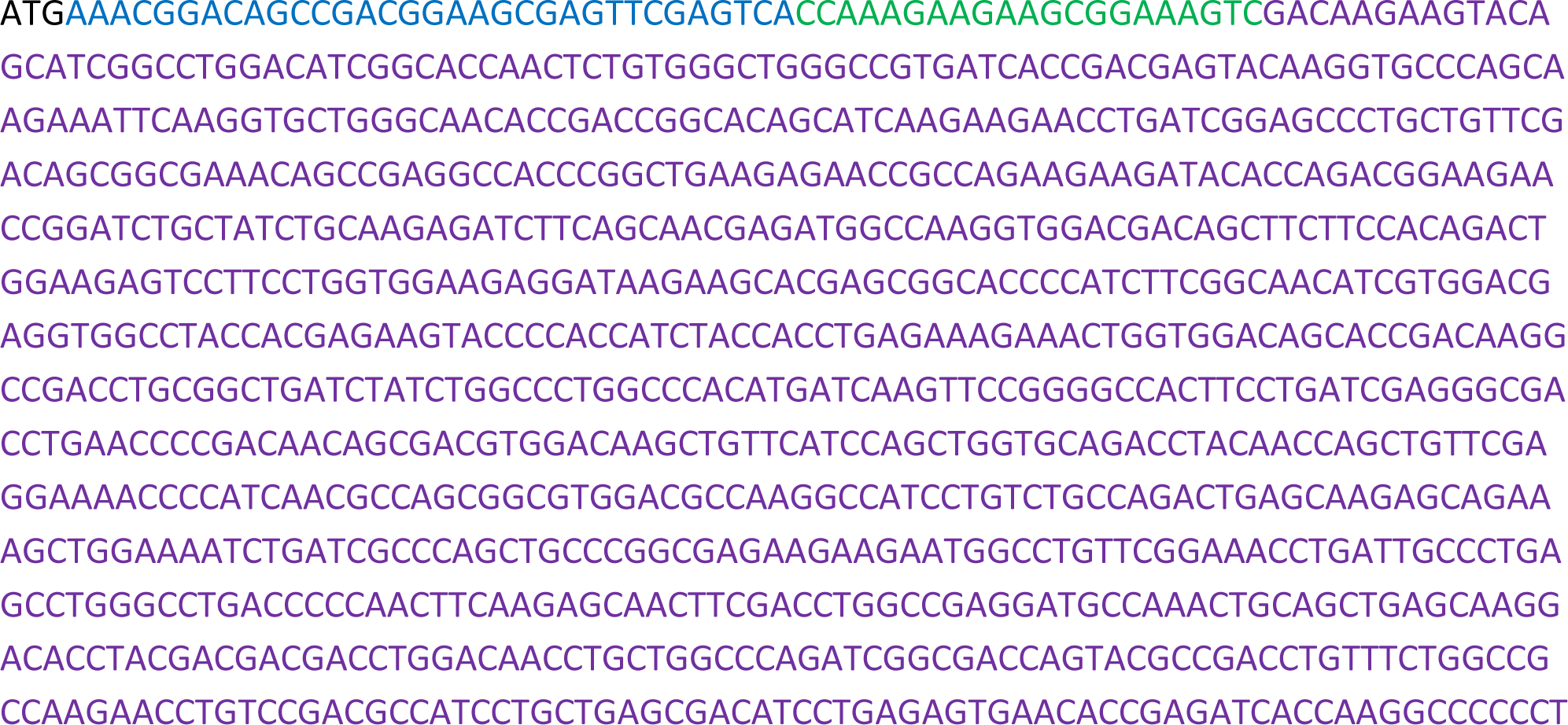
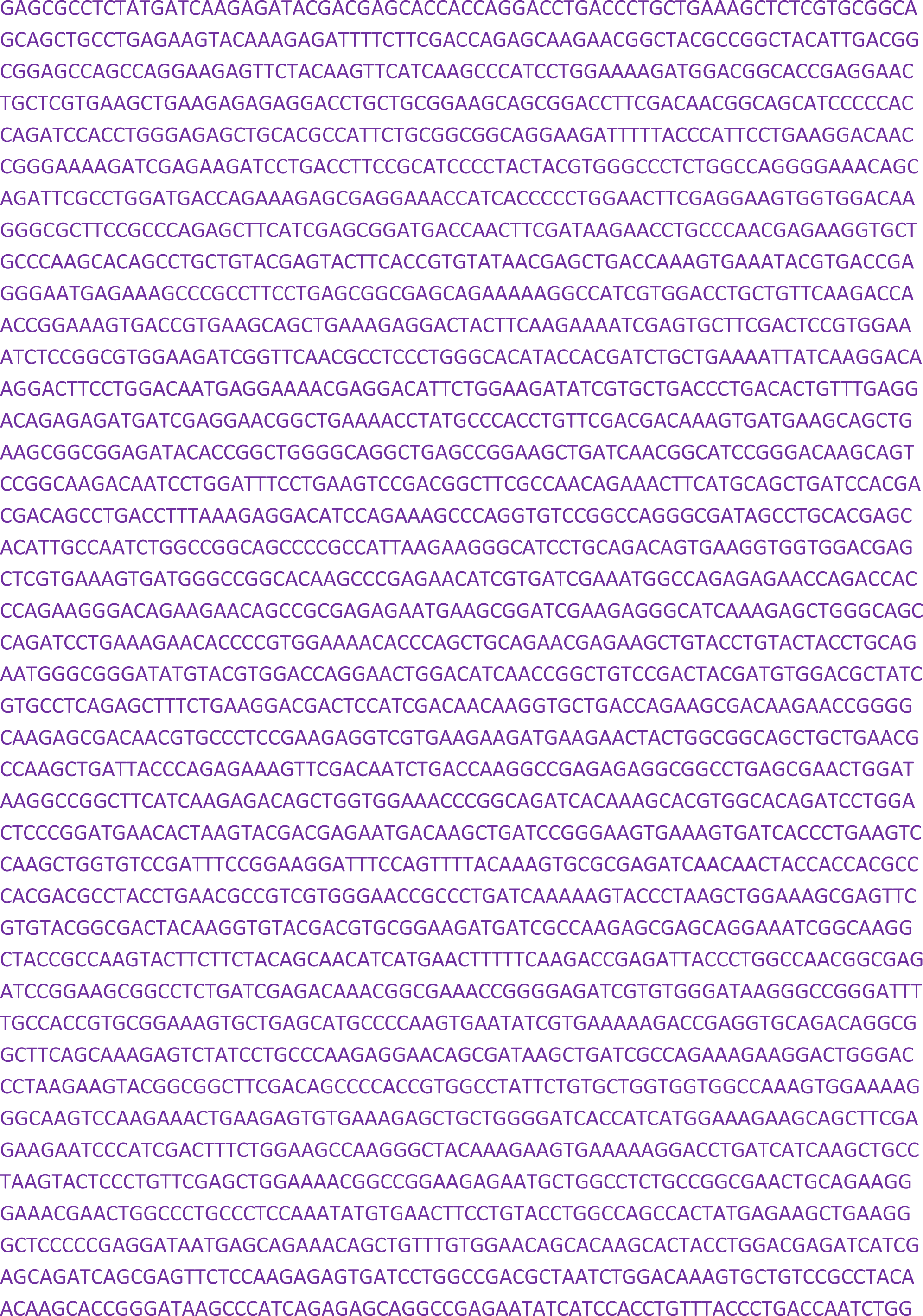
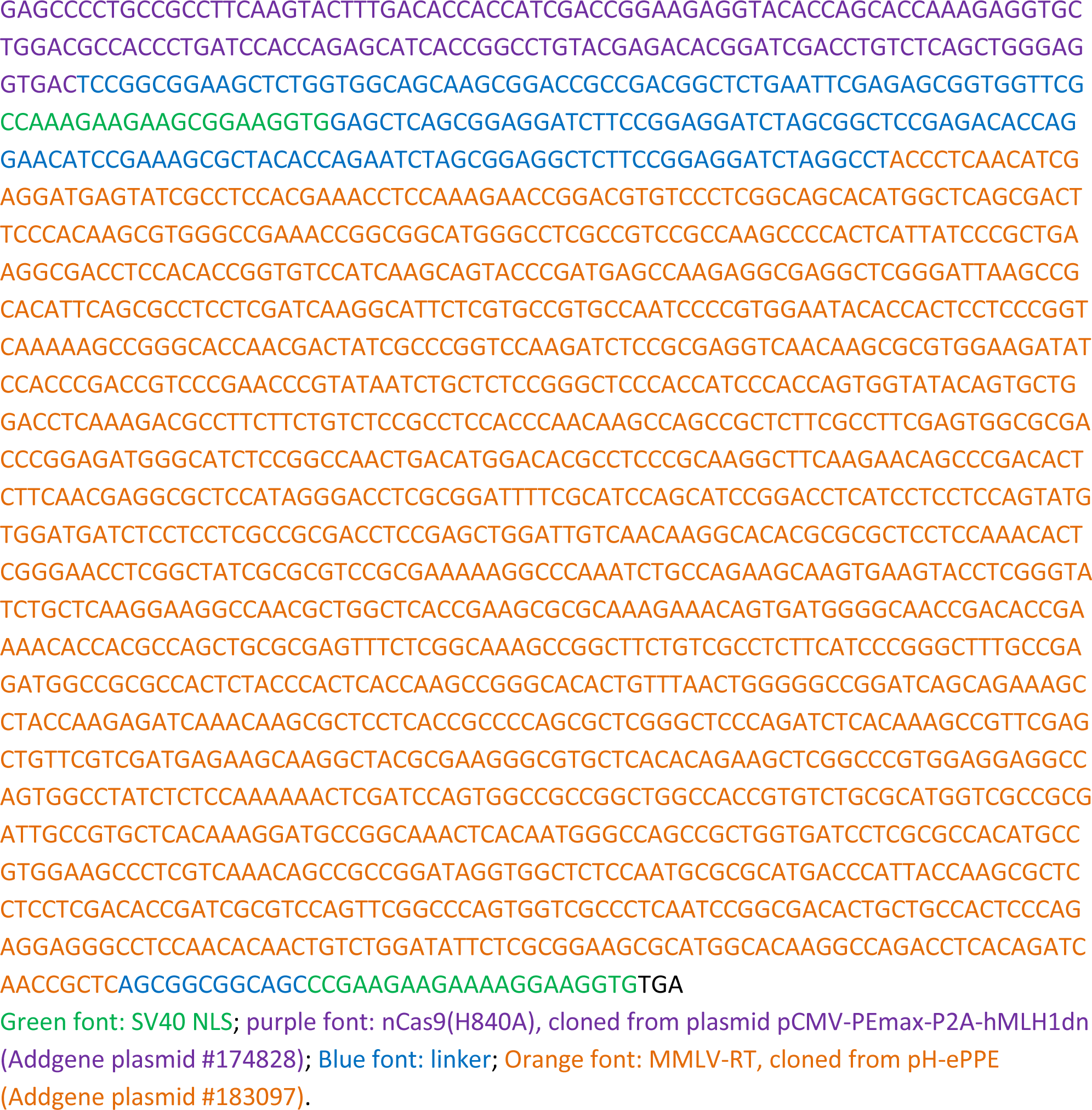

### ePEmax2

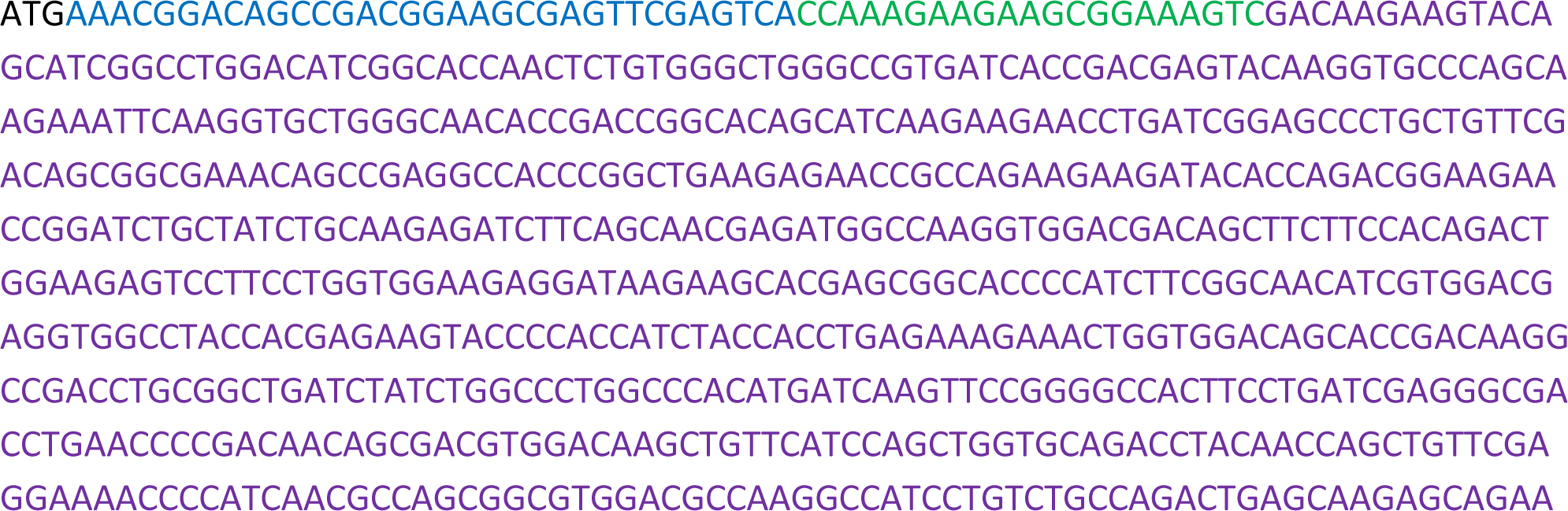
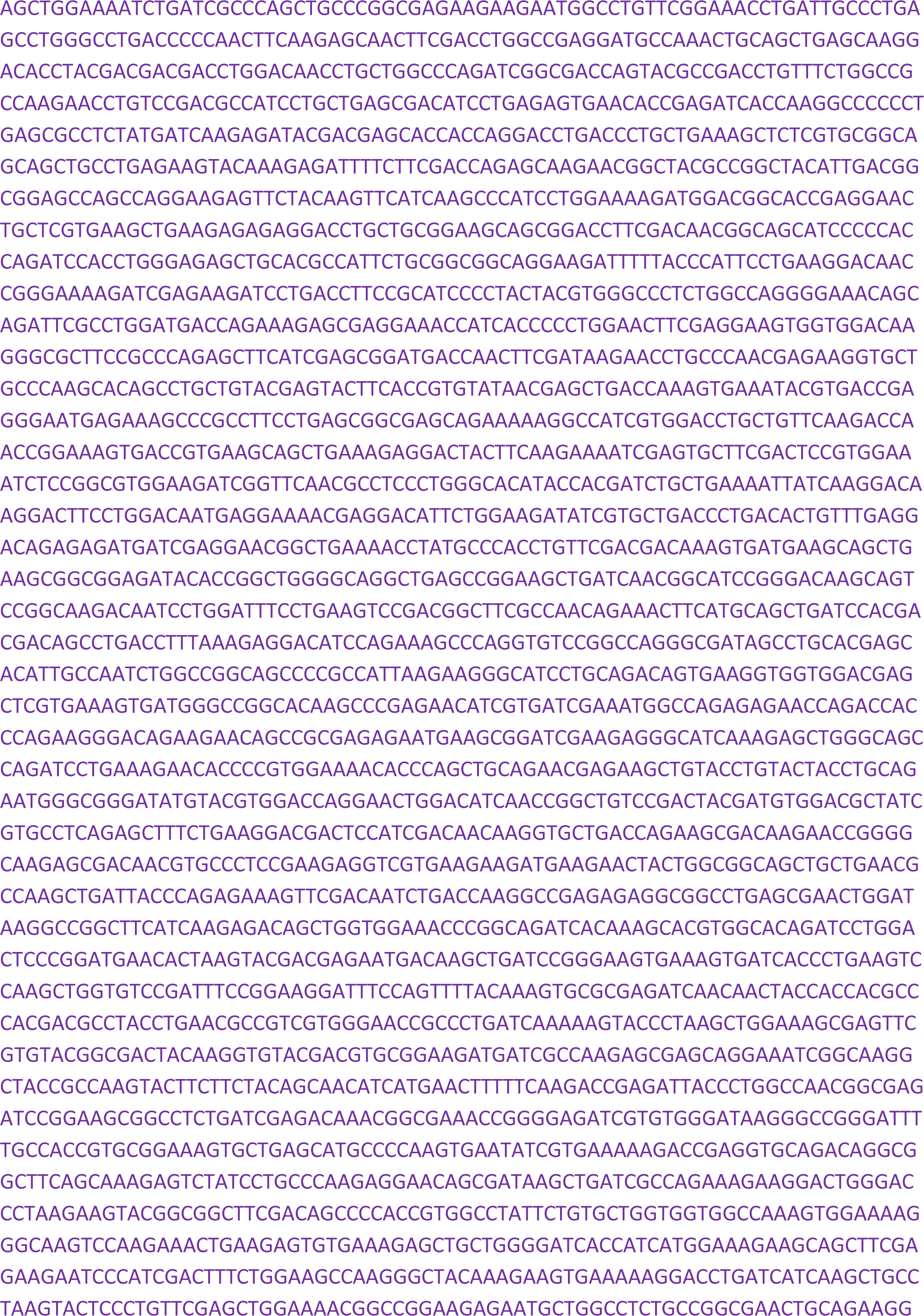
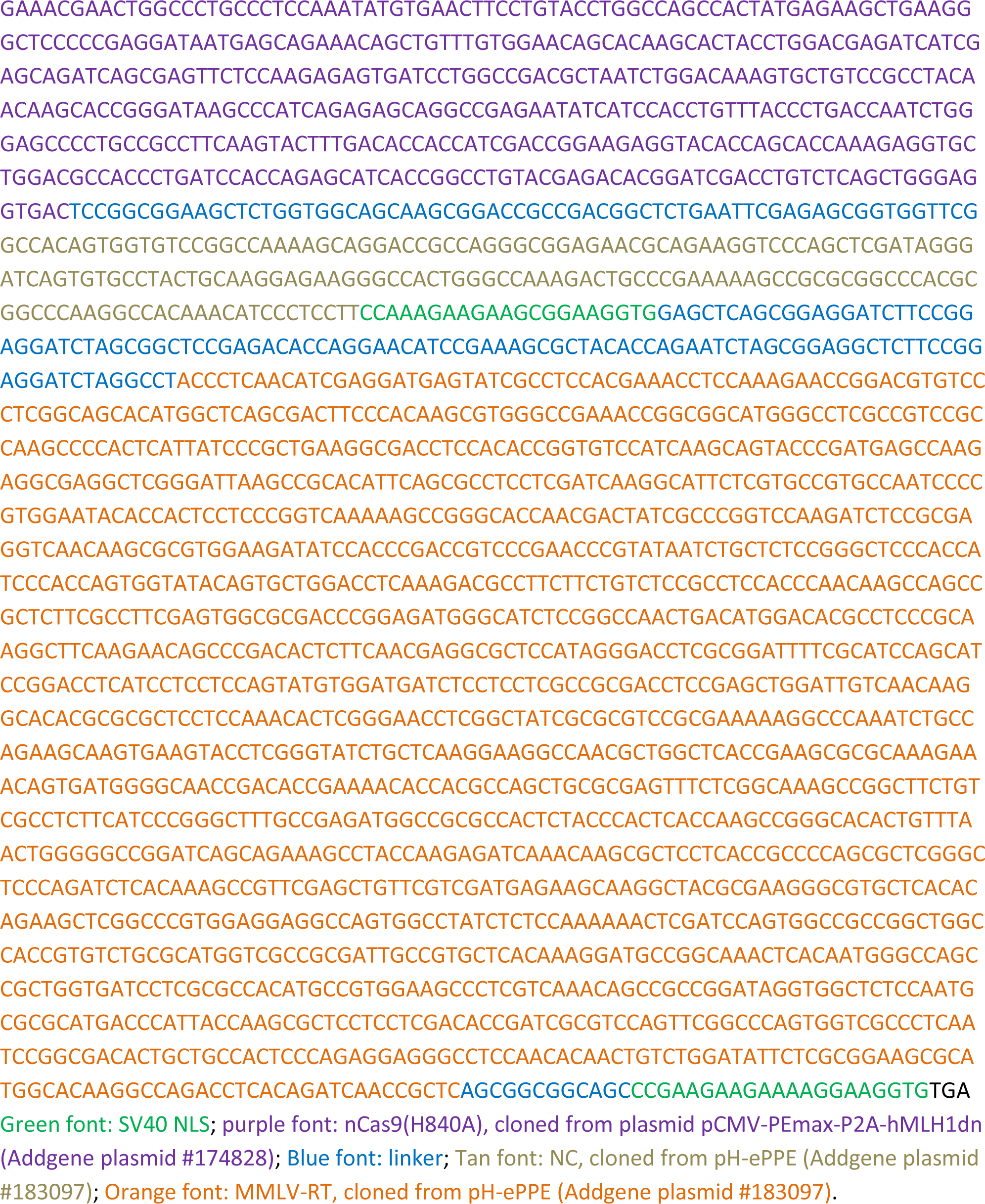

### PE2max-NC

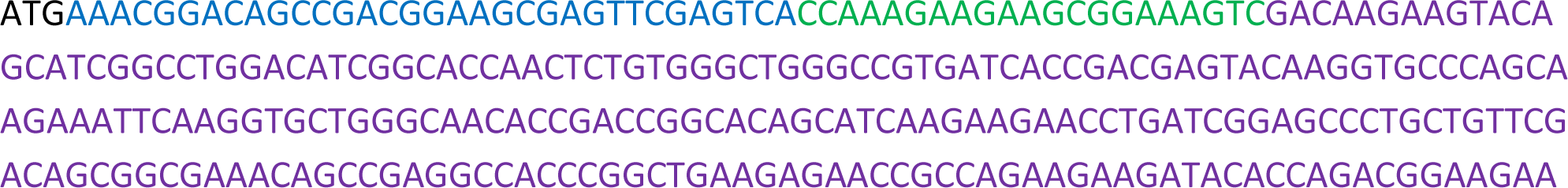
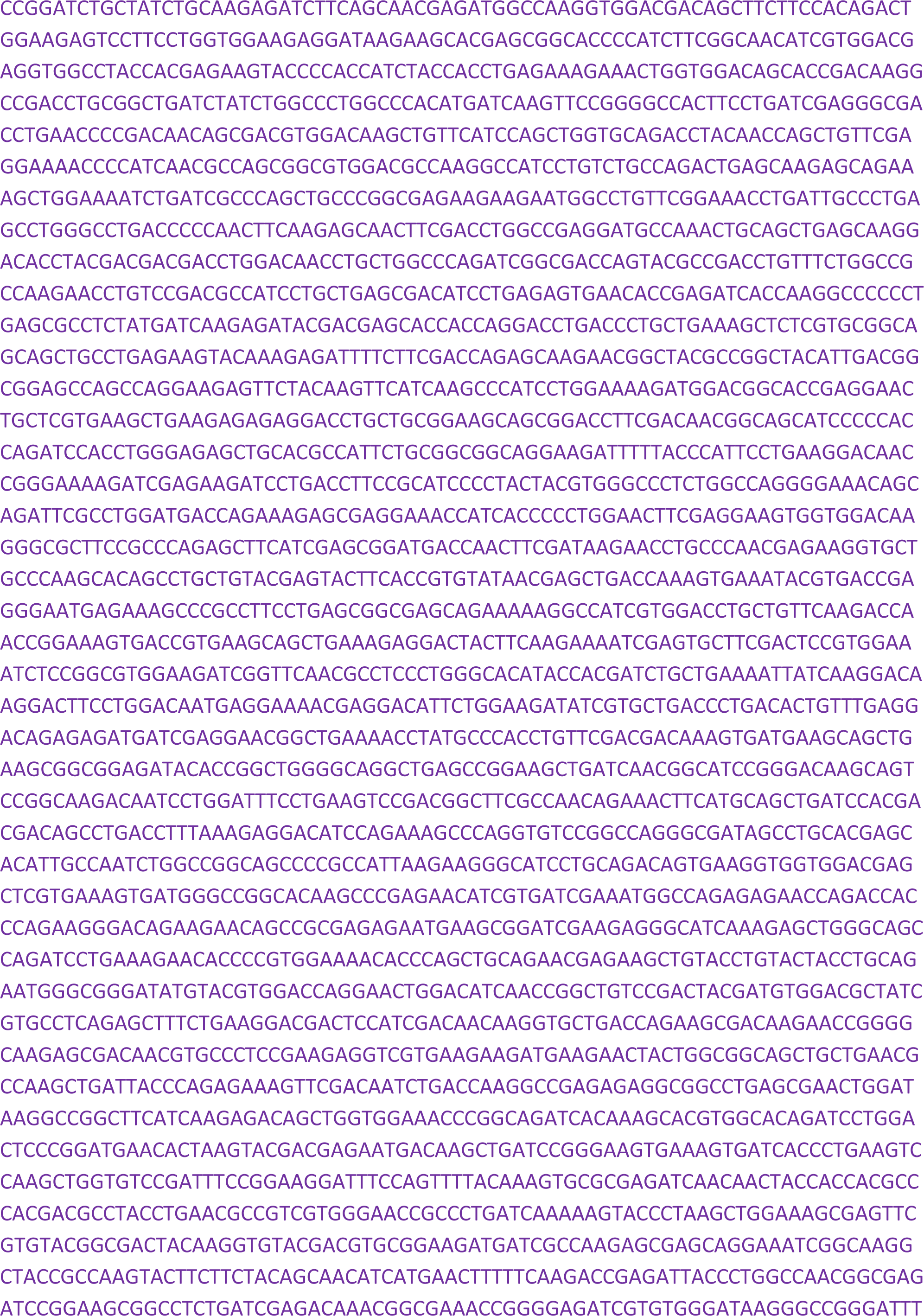
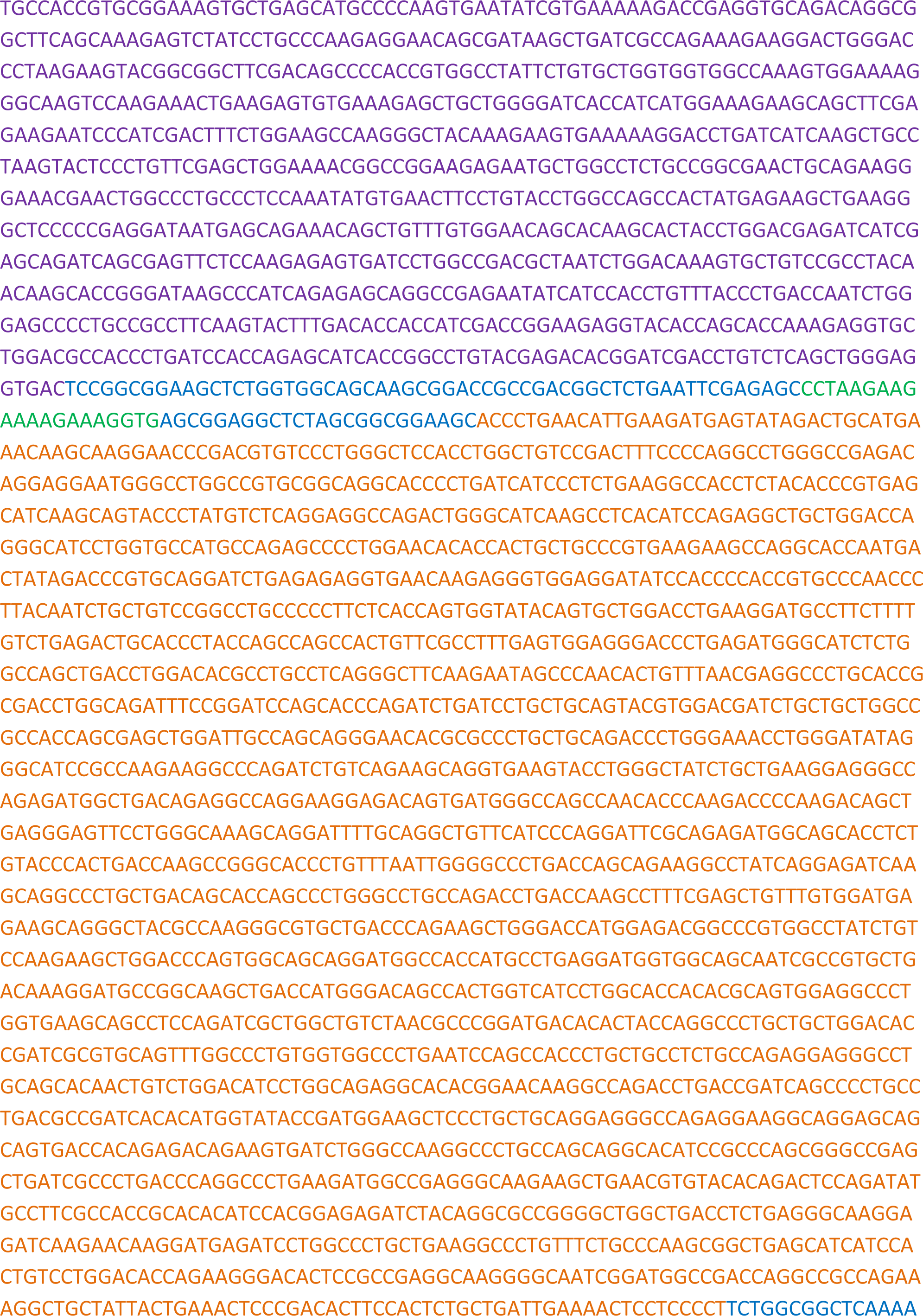
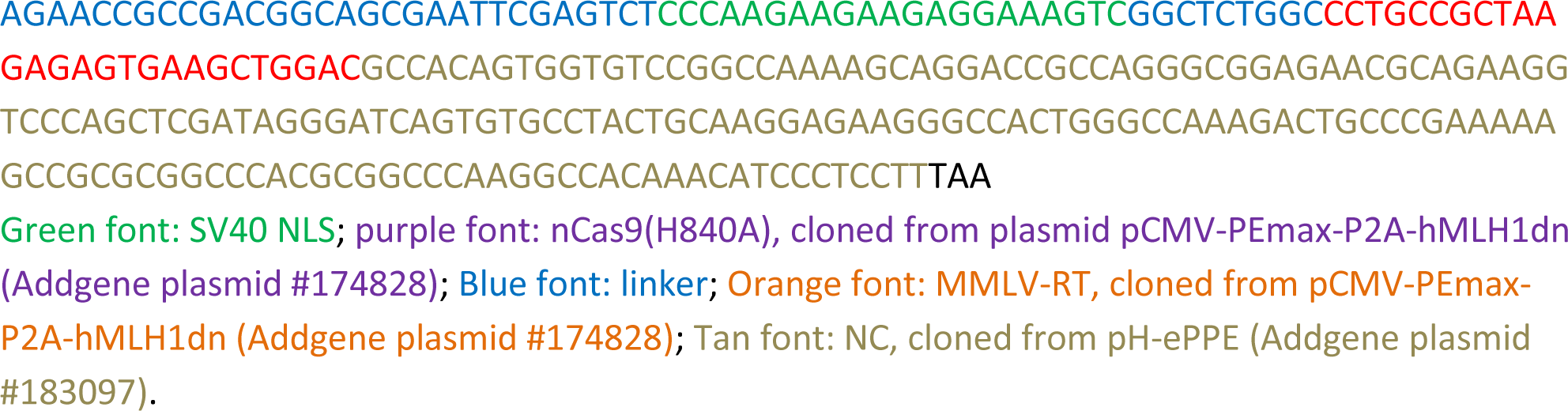

### ePEmax3

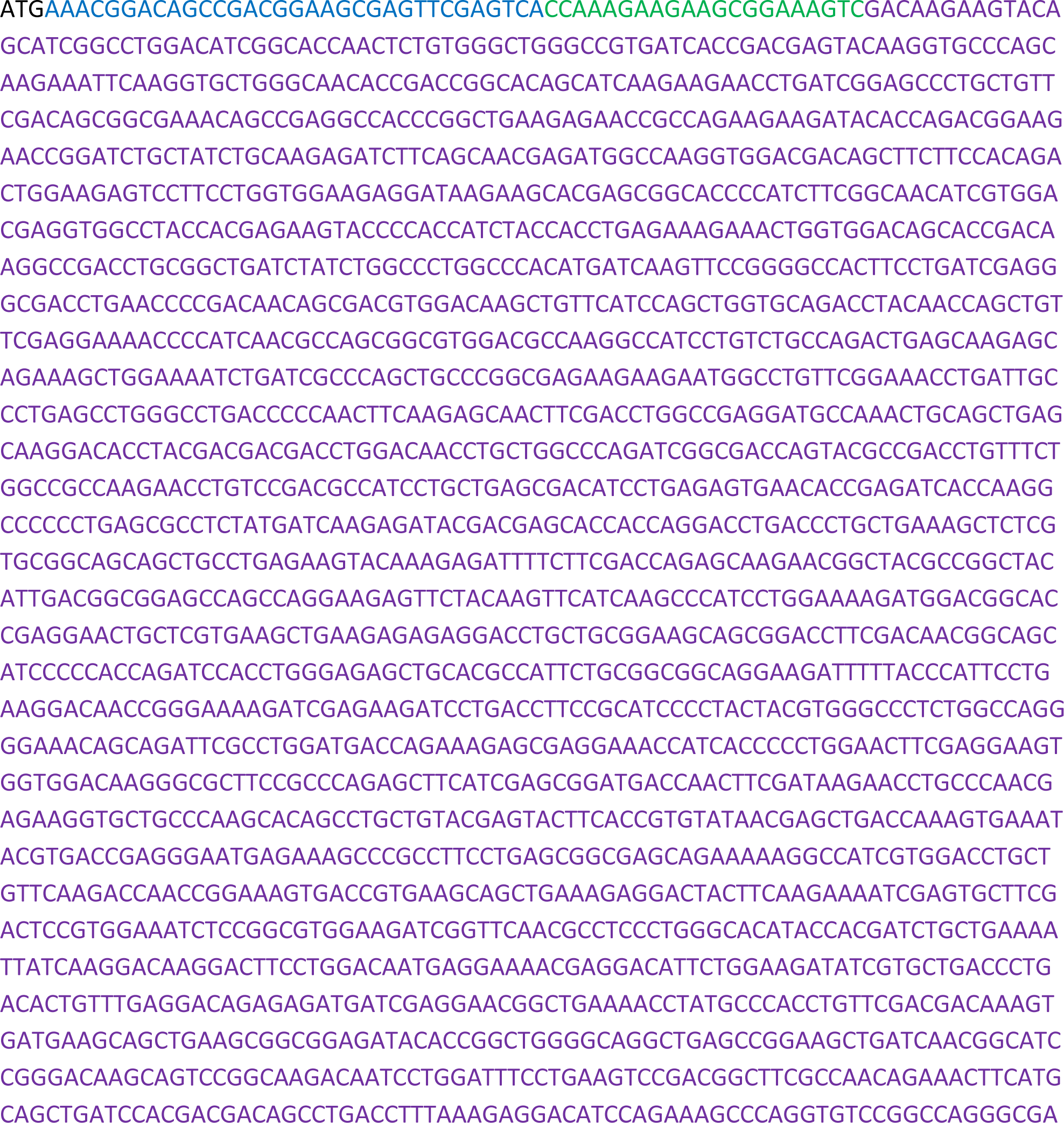
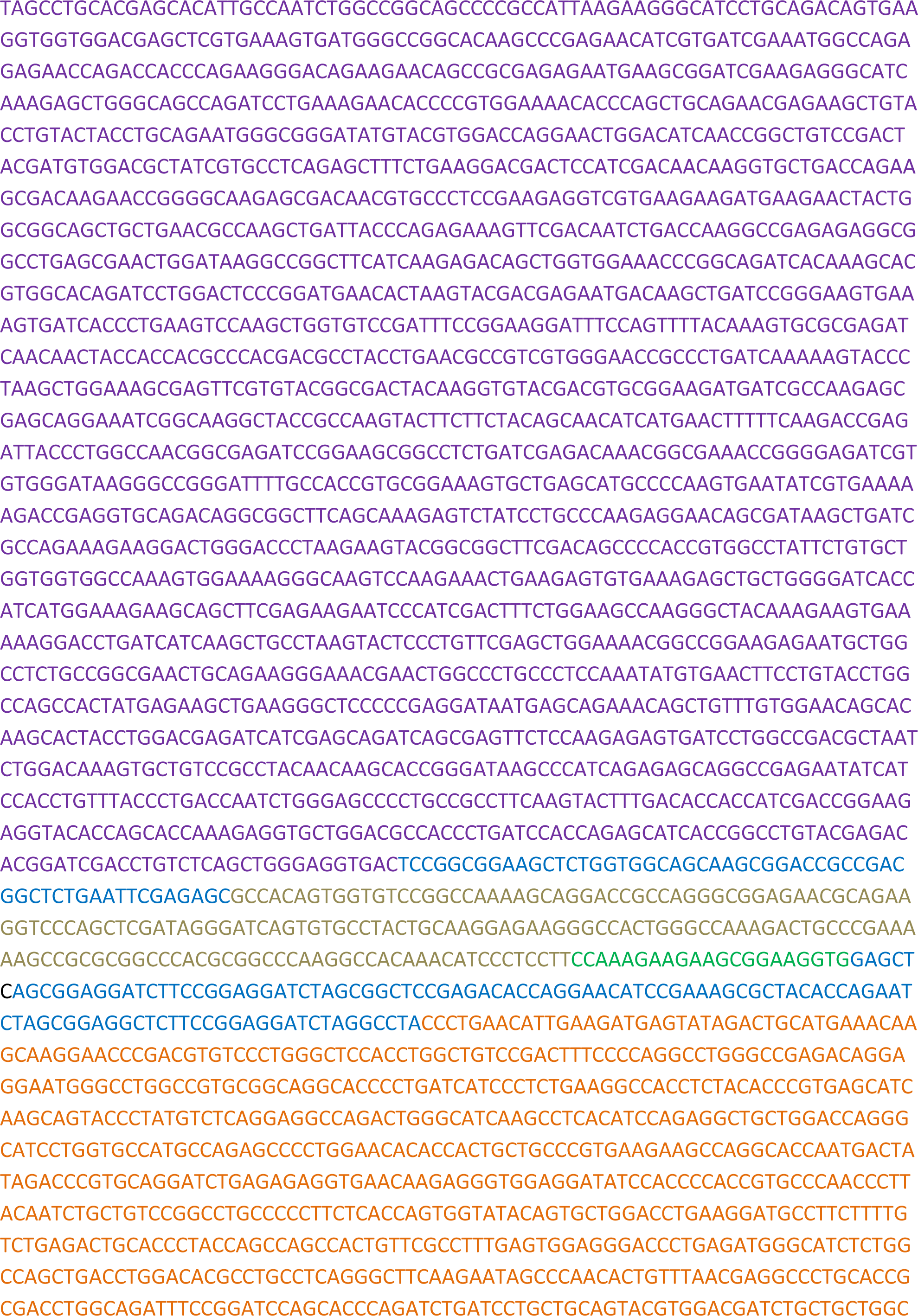
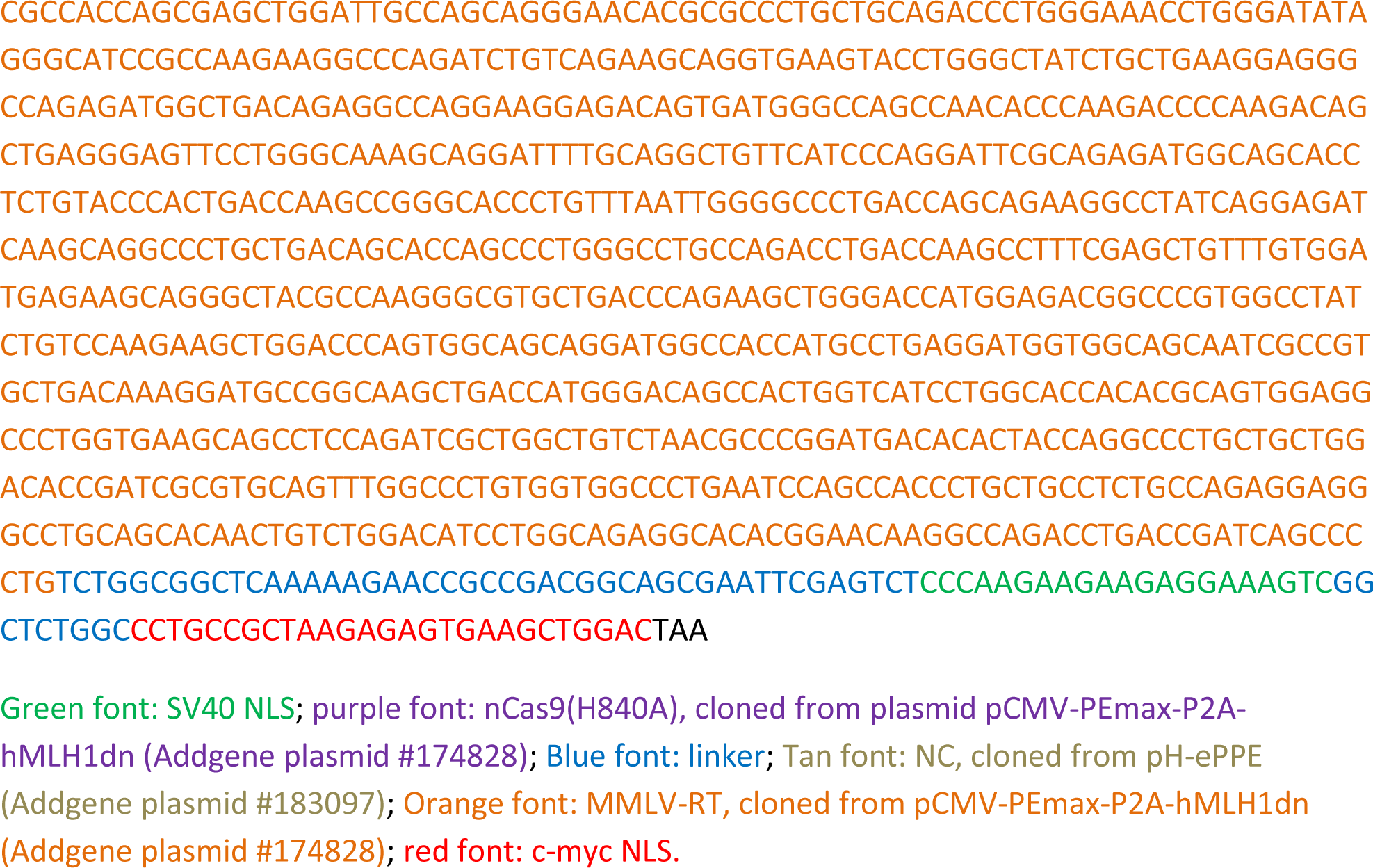

### nCas9max, cloned from plasmid pCMV-PEmax-P2A-hMLH1dn (Addgene plasmid #174828)

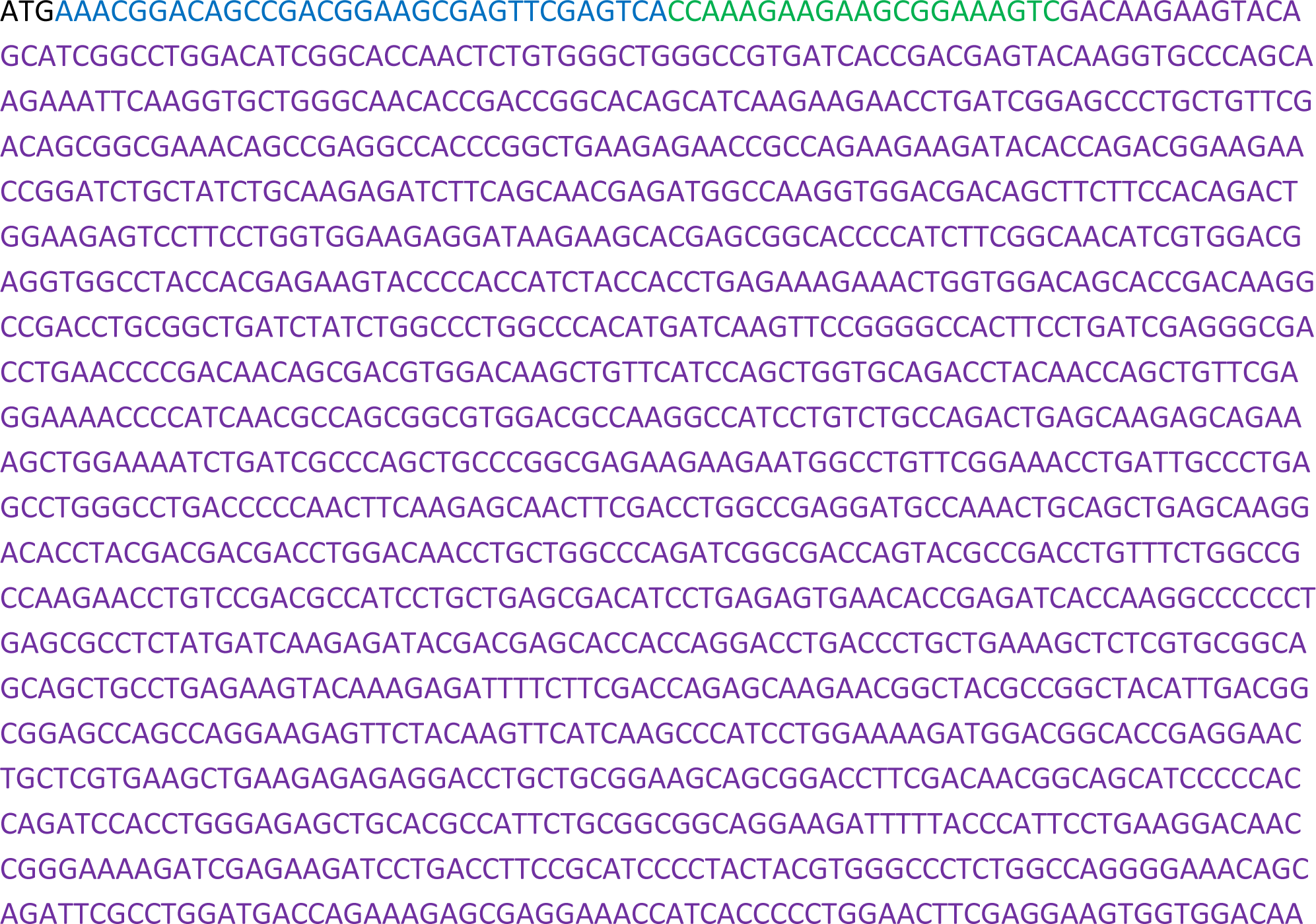
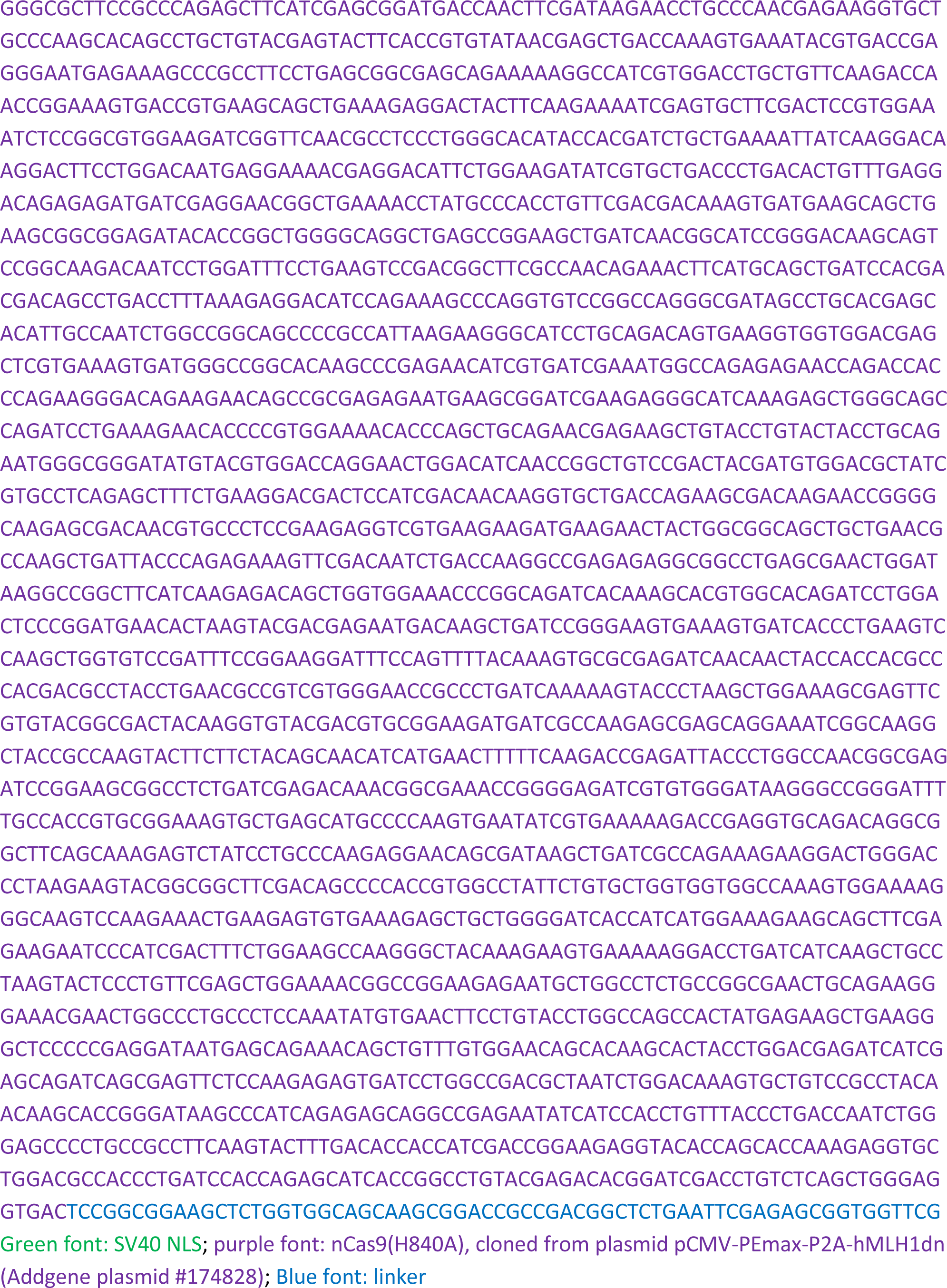

### MCP-MMLV RT, MCP was cloned from pREDIT_MCP-RecT (Plasmid #164803), MMLV RT is a tobacco codon-optimized MMLV RT from the PE2

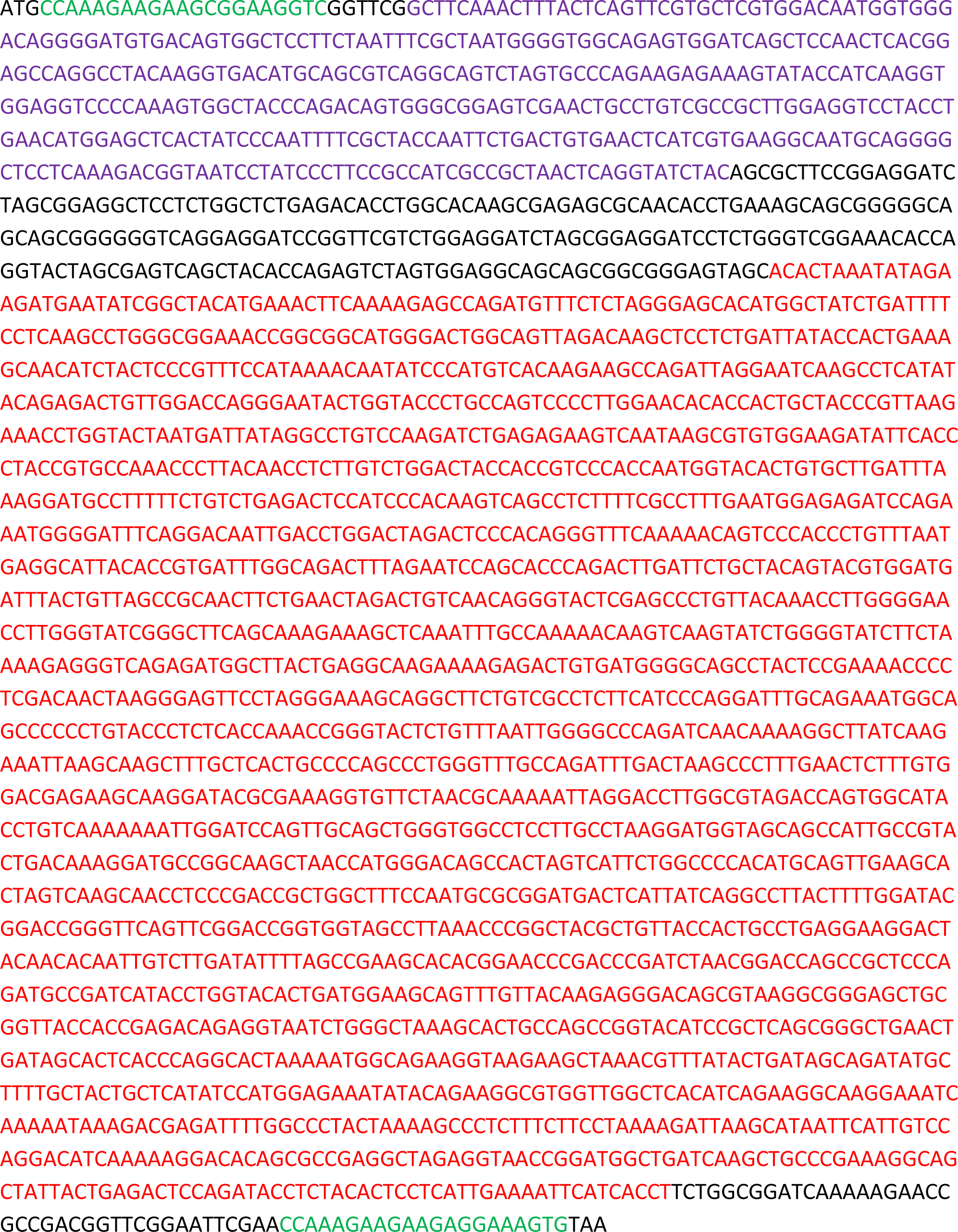
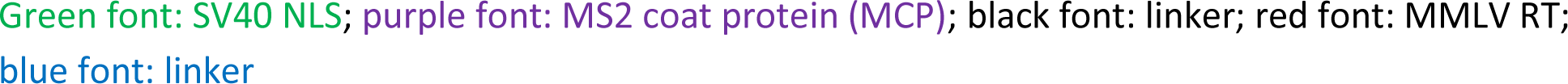

### The CaMV 35S promoter (p35S) was cloned from plasmid pICH51266 (Addgene plasmid #50267)

### EU + Rb7 double terminator (EURb7), synthetic sequence

AAAGCAGAATGCTGAGCTAAAAGAAAGGCTTTTTCCATTTTCGAGAGACAATGAGAAAAGAAGAAGAAGAAGAAGAAGAAGAAGAAGAAGAAAAGAGTAAATAATAAAGCCCCACAGGAGGCGAAGTTCTTGTAGCTCCATGTTATCTAAGTTATTGATATTGTTTGCCCTATATTTTATTTCTGTCATTGTGTATGTTTTGTTCAGTTTCGATCTCCTTGCAAAATGCAGAGATTATGAGATGAATAAACTAAGTTATATTATTATACGTGTTAATATTCTCCTCCTCTCTCTAGCTAGCCTTTTGTTTTCTCTTTTTCTTATTTGATTTTCTTTAAATCAATCCATTTTAGGAGAGGGCCAGGGAGTGATCCAGCAAAACATGAAGATTAGAAGAAACTTCCCTCTTTTTTTTCCTGAAAACAATTTAACGTCGAGATTTATCTCTTTTTGTAATGGAATCATTTCTACAGTTATGACACCAACTCGGTCCATTTGCACCCCTAATCATAATAGCTTTAATATTTCAAGATATTATTAAGTTAACGTTGTCAATATCCTGGAAATTTTGCAAAATGAATCAAGCCTATATGGCTGTAATATGAATTTAAAAGCAGCTCGATGTGGTGGTAATATGTAATTTACTTGATTCTAAAAAAATATCCCAAGTATTAATAATTTCTGCTAGGAAGAAGGTTAGCTACGATTTACAGCAAAGCCAGAATACAAAGAACCATAAAGTGATTGAAGCTCGAAATATACGAAGGAACAAATATTTTTAAAAAAATACGCAATGACTTGGAACAAAAGAAAGTGATATATTTTTTGTTCTTAAACAAGCATCCCCTCTAAAGAATGGCAGTTTTCCTTTGCATGTAACTATTATGCTCCCTTCGTTACAAAAATTTTGGACTACTATTGGGAACTTCTTCTGAAAATAGTG

### U6 composite promoter, reported by Jiang and coworkers in 2021, synthetic sequence

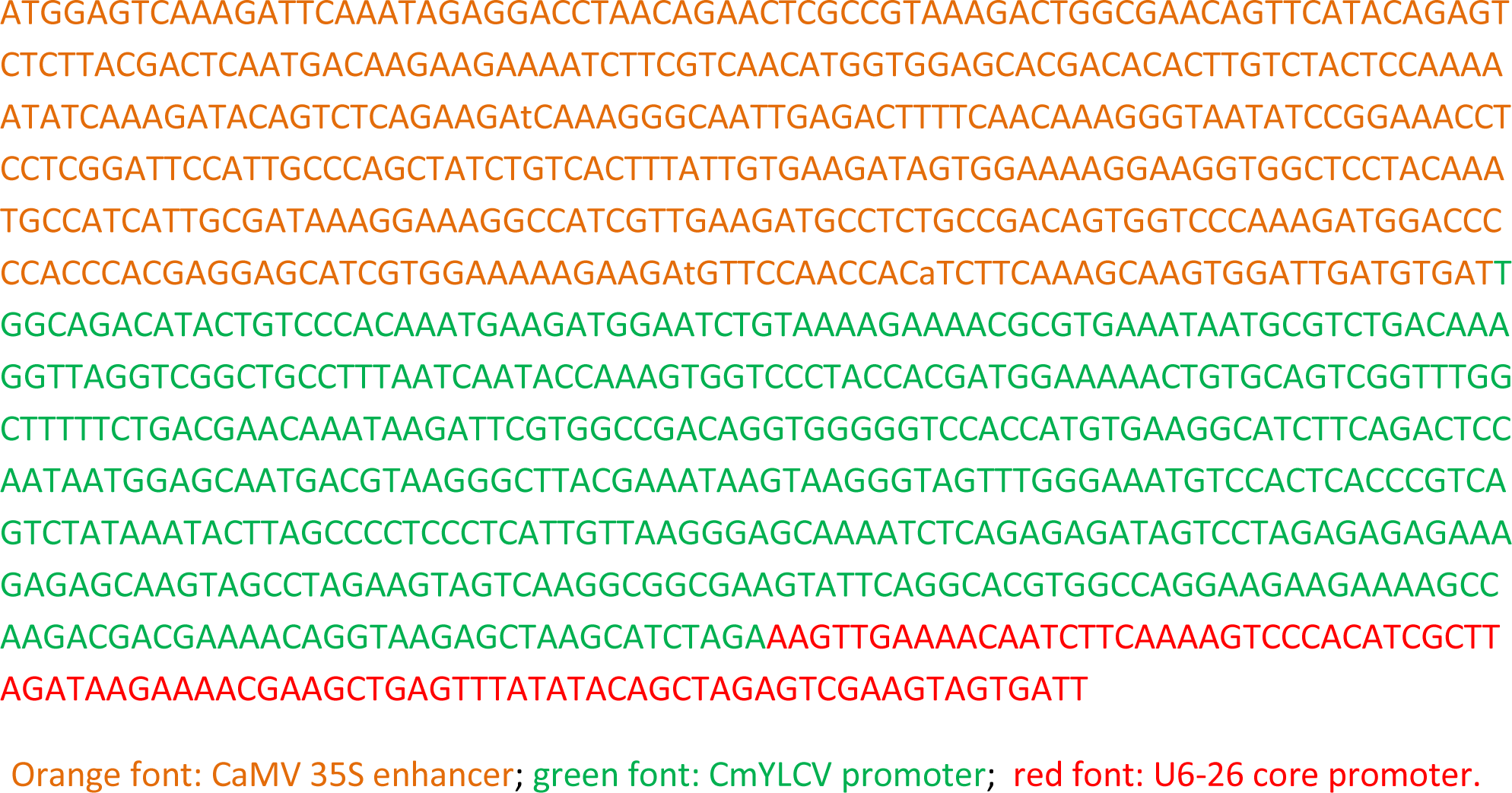

### AtU6-26 core promoter (pU6-26c), cloned from the U6 composite promoter

AAGTTGAAAACAATCTTCAAAAGTCCCACATCGCTTAGATAAGAAAACGAAGCTGAGTTTATATACAGCTAGAGTCGAAGTAGTGATT

The termination sequence used with the U6 promoter was TTTTTTT.

### Altered epegRNA scaffold, synthetic sequence

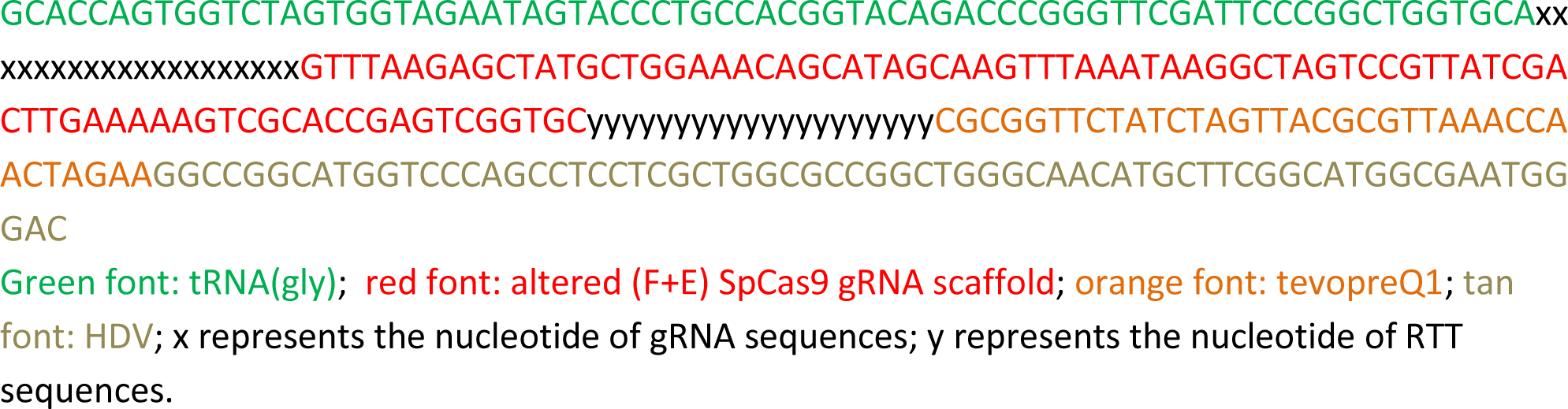

### Supportive gRNA (Sg) sequence

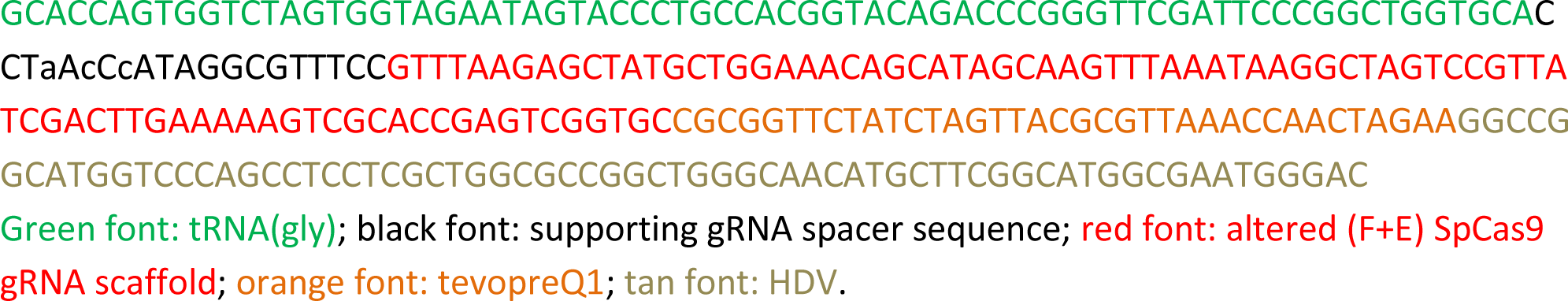

### pegR1.6-MS2 sequence

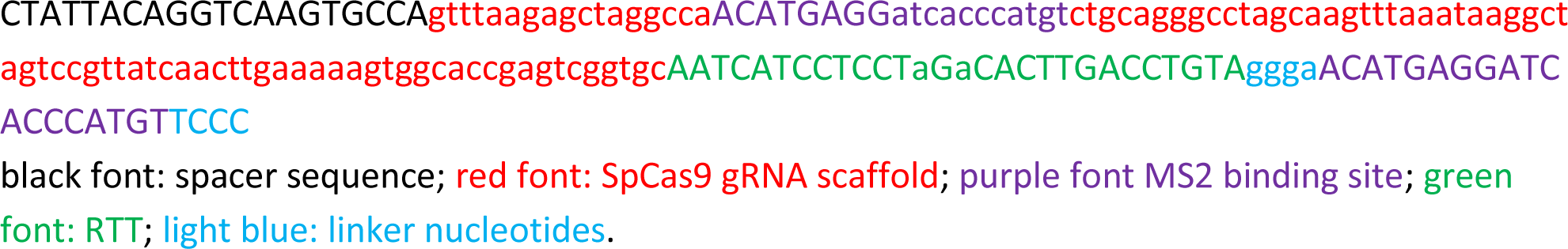

### Plant selection marker: pNOS-NptII-tOCS cloned from pICSL11024 (pICH47732::NOSp-NPTII-OCST) (Addgene Plasmid #51144)

## References

1. Gao, C. Genome engineering for crop improvement and future agriculture. Cell 184, 1621–1635 (2021).

2. Anzalone, A.V., Koblan, L.W. & Liu, D.R. Genome editing with CRISPR-Cas nucleases, base editors, transposases and prime editors. Nat Biotechnol 38, 824-844 (2020).

3. Chen, P.J. et al. Enhanced prime editing systems by manipulating cellular determinants of editing outcomes. Cell 184, 5635–5652 e5629 (2021).

4. Vu, T.V., Nguyen, N.T., Kim, J., Hong, J.C. & Kim, J.-Y. Prime editing: Mechanism insight and recent applications in plants. Plant Biotechnology Journal 22, 19–36 (2024).

5. Jiang, Y.Y. et al. Prime editing efficiently generates W542L and S621I double mutations in two ALS genes in maize. Genome Biol 21, 257 (2020).

6. Jin, S., Lin, Q., Gao, Q. & Gao, C. Optimized prime editing in monocot plants using PlantPegDesigner and engineered plant prime editors (ePPEs). Nat Protoc 18, 831-853 (2023).

7. Li, J. et al. Prime editing-mediated precise knock-in of protein tag sequences in the rice genome. Plant Commun 4, 100572 (2023).

8. Lin, Q. et al. High-efficiency prime editing with optimized, paired pegRNAs in plants. Nat Biotechnol 39, 923–927 (2021).

9. Lin, Q. et al. Prime genome editing in rice and wheat. Nat Biotechnol 38, 582–585 (2020).

10. Qiao, D. et al. Optimized prime editing efficiently generates heritable mutations in maize. J Integr Plant Biol 65, 900–906 (2023).

11. Sun, C., et al. Precise integration of large DNA sequences in plant genomes using PrimeRoot editors. Nat Biotechnol Online ahead of print. (2023).

12. Zong, Y., et al. An engineered prime editor with enhanced editing efficiency in plants. Nat Biotechnol 40, 1394-1402 (2022).

13. Li, J., et al. Development of a highly efficient prime editor 2 system in plants. Genome Biology 23, 161 (2022).

14. Lu, Y. et al. Precise genome modification in tomato using an improved prime editing system. Plant Biotechnol J 19, 415–417 (2021).

15. Vu, T.V. et al. The obstacles and potential solution clues of prime editing applications in tomato. BioDesign Res 2022, 0001 (2022).

16. Perroud, P.F. et al. Prime Editing in the model plant Physcomitrium patens and its potential in the tetraploid potato. Plant Sci 316, 111162 (2022).

17. Wang, L. et al. Spelling changes and fluorescent tagging with prime editing vectors for plants. Front Genome Ed 3, 617553 (2021).

18. Zou, J. et al. Improving the efficiency of prime editing with epegRNAs and high-temperature treatment in rice. Sci China Life Sci 65, 2328–2331 (2022).

19. Vu, T.V. et al. Highly efficient homology-directed repair using CRISPR/Cpf1-geminiviral replicon in tomato. Plant Biotechnol J 18, 2133–2143 (2020).

20. Nelson, J.W. et al. Engineered pegRNAs improve prime editing efficiency. Nat Biotechnol 40, 402–410 (2022).

21. Diamos, A.G. & Mason, H.S. Chimeric 3’ flanking regions strongly enhance gene expression in plants. Plant Biotechnol J 16, 1971–1982 (2018).

22. Anzalone, A.V. et al. Search-and-replace genome editing without double-strand breaks or donor DNA. Nature 576, 149–157 (2019).

23. Doman, J.L., et al. Phage-assisted evolution and protein engineering yield compact, efficient prime editors. Cell 186, 3983-4002 e3926 (2023).

24. LeBlanc, C. et al. Increased efficiency of targeted mutagenesis by CRISPR/Cas9 in plants using heat stress. The Plant Journal 93, 377–386 (2018).

25. Malzahn, A.A. et al. Application of CRISPR-Cas12a temperature sensitivity for improved genome editing in rice, maize, and Arabidopsis. BMC Biology 17, 9 (2019).

26. Chen, P.J. & Liu, D.R. Prime editing for precise and highly versatile genome manipulation. Nat Rev Genet 24, 161–177 (2023).

27. Martinez-Miguel, V.E. et al. Increased fidelity of protein synthesis extends lifespan. Cell Metab 33, 2288–2300 e2212 (2021).

28. Chen, K., Wang, Y., Zhang, R., Zhang, H. & Gao, C. CRISPR/Cas Genome Editing and Precision Plant Breeding in Agriculture. Annu Rev Plant Biol 70, 667–697 (2019).

29. Gao, C. The future of CRISPR technologies in agriculture. Nat Rev Mol Cell Biol 19, 275–276 (2018).

30. Zhang, X. & Mason, H. Bean Yellow Dwarf Virus replicons for high-level transgene expression in transgenic plants and cell cultures. Biotechnol Bioeng 93, 271–279 (2006).

31. Cermak, T., Baltes, N.J., Cegan, R., Zhang, Y. & Voytas, D.F. High-frequency, precise modification of the tomato genome. Genome Biol 16, 232 (2015).

32. Li, X. et al. Highly efficient prime editing by introducing same-sense mutations in pegRNA or stabilizing its structure. Nat Commun 13, 1669 (2022).

33. Weber, E., Engler, C., Gruetzner, R., Werner, S. & Marillonnet, S. A modular cloning system for standardized assembly of multigene constructs. PLoS One 6, e16765 (2011).

34. Nonaka, S., Someya, T., Kadota, Y., Nakamura, K. & Ezura, H. Super-Agrobacterium ver. 4: Improving the Transformation Frequencies and Genetic Engineering Possibilities for Crop Plants. Front Plant Sci 10, 1204 (2019).

35. Park, J., Lim, K., Kim, J.-S. & Bae, S. Cas-analyzer: an online tool for assessing genome editing results using NGS data. Bioinformatics 33, 286–288 (2016).

36. Clement, K. et al. CRISPResso2 provides accurate and rapid genome editing sequence analysis. Nat Biotechnol 37, 224–226 (2019).

37. Conant, D. et al. Inference of CRISPR Edits from Sanger Trace Data. Crispr j 5, 123–130 (2022).

38. Lichtenthaler, H.K. & Buschmann, C. Chlorophylls and Carotenoids: Measurement and Characterization by UV-VIS Spectroscopy. Current Protocols in Food Analytical Chemistry 1, F4.3.1–F4.3.8 (2001).

